# Time-dependent bistability leads to critical slowing down during floral transition in Arabidopsis

**DOI:** 10.64898/2025.12.09.692976

**Authors:** G. Rodríguez-Maroto, K. Wang, P. Casanova-Ferrer, M. Cerise, G. Coupland, P. Formosa-Jordan

## Abstract

Developmental transitions occur in the life cycles of all multicellular organisms. Despite their fundamental relevance, the underlying dynamics remain poorly understood. In plants, floral transition is a key developmental process whereby the shoot apical meristem changes from producing leaves to forming flowers. Using quantitative imaging, developmental genetics and dynamical systems theory, we show that a time-dependent bistable switch between expression of *APETALA2*, a key floral inhibitor, and the floral activators *SUPPRESSOR OF OVEREXPRESSION OF CONSTANS 1* and *FRUITFULL*, can explain the dynamics of floral transition in Arabidopsis. Notably, we detect a slowing down of the inhibitor dynamics, consistent with the system crossing a critical point of a bistable switch and transiently experiencing a ghost attractor. We demonstrate that this time-dependent bistability is essential to generate the range of dynamical behaviours measured across genotypes, including oscillations in the inhibitor, and that it also confers robustness in the transition. Collectively, our work provides quantitative evidence of time-dependent bistability underlying floral transition, which introduces a new timescale to this developmental process.

## Introduction

Biological systems progress through their developmental programmes via a succession of distinct states. These transitions between distinct states are fundamental across life, and include, for example, transitions at the cellular level, such as cell-fate decisions^1^, transitions at the multicellular level, such as phase transitions in morphogenetic processes^2^, and transitions at the organism level, such as the metamorphosis in amphibians^3^. Long-standing questions remain concerning the dynamics and timing of these transitions, and how they generate appropriate responses against changing stimuli. Conrad Waddington was a pioneer in investigating the theoretical basis of these questions^4^. In his iconic illustration known as the Waddington landscape, the state of a developing system is represented by a ball rolling down an undulated surface. Each valley represents a distinct developmental route, and the system’s trajectory through a valley represents the development of the system4. A key feature of Waddington’s landscape is that it can be dynamic — genetic or environmental signals can alter its topography^4,5^ (Figure 1A). This time-dependent nature is particularly relevant in the context of developmental transitions, where understanding the system’s transient behaviour is essential. Furthermore, Waddington’s landscape can be connected to bifurcation theory in dynamical systems, where it is defined as a potential or quasi-potential function derived from the underlying dynamical system equations^5–8^.

**Figure 1.**
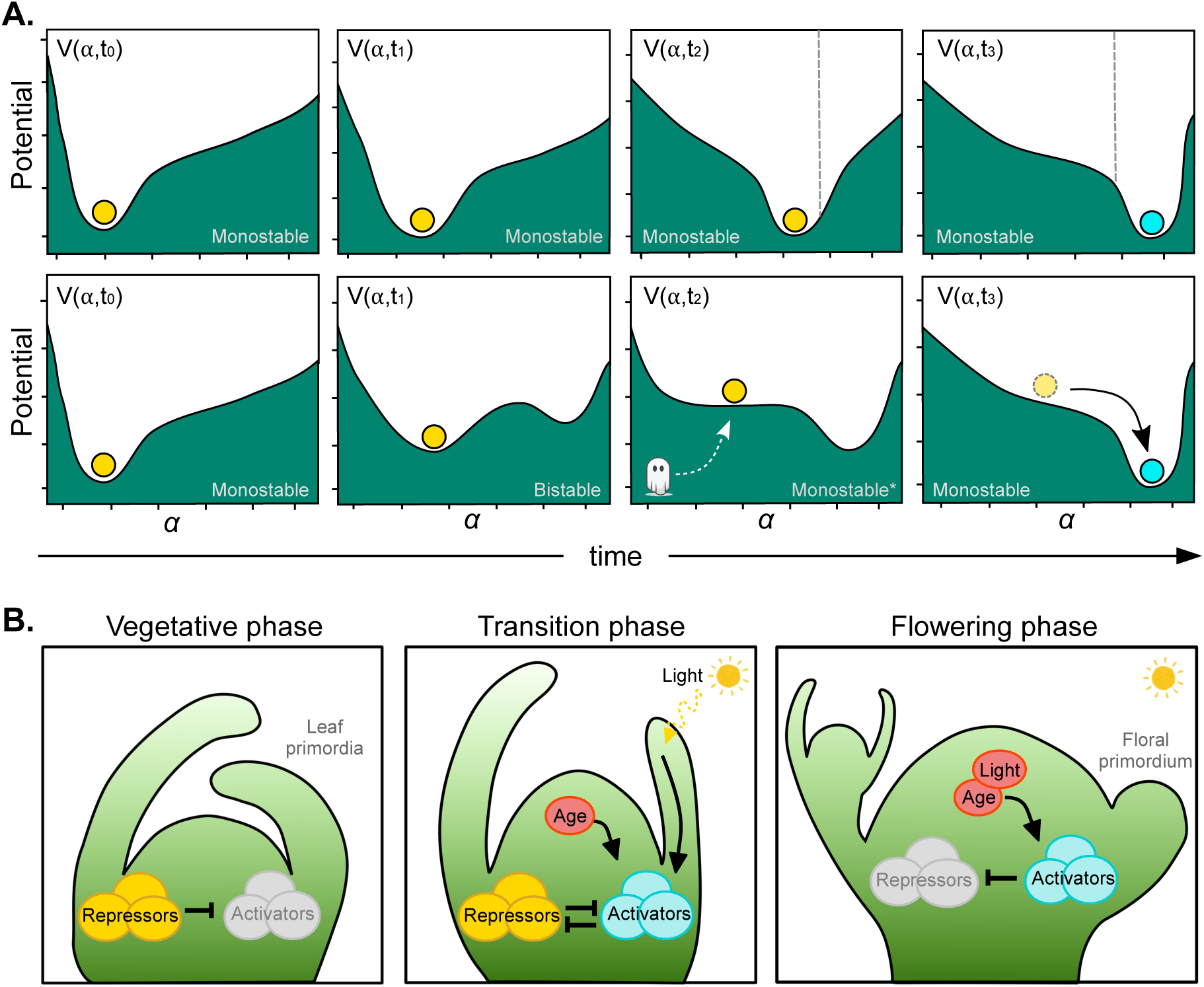
Monostable versus bistable transitions in the context of floral transition at the SAM. A) Waddington/quasi-potential landscape cartoons illustrating a monostable (top row) and a bistable (bottom row) transition. The system’s state is depicted as a marble that is attracted to stable steady states (stable attractors), located at the bottom of the valleys. In the monostable transition (top row), the only existing valley changes over time. A threshold (grey dashed line) arbitrarily separates developmental state A (yellow) from developmental state B (blue). In the bistable transition (bottom row), two different valleys coexist for some time, each containing a stable steady state representing a different developmental state. The system only moves to the new stable attractor when the initial stable attractor disappears. The disappearance of the attractor flattens the landscape, and the system undergoes a slowing down of the dynamics, remaining in a transient state reminiscent of the previous unstabilised state, known as a ghost state. B) Scheme of simplified regulatory network underlying floral transition at the SAM. Light and age factors modulate mutual inhibition between floral activators and inhibitors over time. Normal arrows indicate activation; blunt arrows represent inhibition.

In the context of the Waddington landscape, a developmental transition can be characterised by two main different dynamical scenarios (Figure 1A). One corresponds to a monostable transition, where a single valley contains a stable steady state (attractor) to which the system converges throughout development (Figure 1A, top row). In this scenario, the valley can shift its position and change shape over time, with the developmental system’s change in identity being represented either by a continuous shift or a sharp change upon crossing a positional threshold (Figure 1A, top row). The second scenario involves the coexistence, at some point of development, of two valleys, each containing a stable steady state. This is known as bistability^7–10^ and we refer to the switch from one stable state to the other as a bistable transition (Figure 1A, bottom row). Assuming the developmental trajectory is deterministic, i.e., the system trajectory does not undergo random fluctuations, this sharp and discontinuous transition occurs when the initial stable attractor disappears, forcing the system’s trajectory out of the first valley, and eventually converging on the second stable attractor^7^ (Figure 1A, bottom row). The change in the number or type of attractors (i.e., stable and unstable steady states) when the system is perturbed (i.e., changes in a model parameter) is known as a bifurcation^11,12^ (saddle-node bifurcation, in the case above) and the state at the bifurcation point itself is referred to as a critical state. Thus, a bistable transition would involve undergoing a saddle-node bifurcation over time, passing through criticality, which has important dynamic consequences. After the stable attractor disappears, its influence on the system’s dynamics persists due to the flattened landscape surface, called a ghost attractor^13–17^ (Figure 1A, third panel in lower row). This phenomenon, which results in a slowing down of the system’s dynamics, is known as critical slowing down^13–17^, and has been proposed to occur in several biological processes, from neural spiking^18,19^, to embryonic development^20,21^ and population dynamics^22^.

Bistability requires a nonlinear positive feedback loop^7,9^ and can confer robustness and hysteresis to the system^5,9,10,23,24^. These emerging properties make bistability a key dynamical mechanism^10,25^, particularly in contexts such as cell signalling^1,26,27^, and differentiation^28^, where it can explain how an undifferentiated cell progresses to one of the available differentiated cell fates during development^7,29^. Yet, despite its theoretical relevance in biology, proving the existence of bistability in a biological system remains a significant challenge, especially in contexts where it enables developmental transitions. Furthermore, there is still limited experimental evidence supporting critical slowing down and its relevance in biology. Although the gene regulatory networks (GRN) driving key developmental decisions such as the floral transition are increasingly known^30,31^, the underlying dynamical mechanisms that enable robust yet flexible transitions against genetic and environmental variations remain poorly understood.

The plant life cycle includes several developmental transitions, but only a few studies have investigated their underlying dynamic behaviour^32–36^. Floral transition is one of these crucial steps in plant development, marking the progression from vegetative growth to the reproductive stage, and is strongly influenced by endogenous and environmental cues that regulate its timing^37,38^. Only precise spatiotemporal coordination between gene expression dynamics and morphological and physiological changes enables reproductive success and completion of seed development under favourable conditions^37^. In *Arabidopsis thaliana*, these simultaneous processes converge at the shoot apical meristem (SAM) (Figure 1B), which contains a population of stem cells whose progeny give rise to aerial organs^39^: leaves in the vegetative state and lateral branches and flowers in the inflorescence phase, also known as the flowering phase. The SAM is also the tissue where relevant changes in GRN dynamics controlling floral transition occur^40–43^. During the vegetative phase, flowering is strongly repressed by APETALA2 (AP2) and APETALA2-LIKE (AP2-LIKE) transcription factors (TFs). These repressors inhibit the transcription of the floral activators *FRUITFULL* (*FUL*) and *SUPPRESSOR OF OVEREXPRESSION OF CONSTANS 1* (*SOC1*), both encoding MADS box TFs, and *MIR172* genes, which encode microRNA172 (miR172) that targets the mRNAs of *AP2* and *AP2-LIKE* genes^44,45^ (Figure 1B, left panel). Exposure to inductive long-day (LD) conditions (16 h light and 8 h darkness) activates the photoperiodic pathway, which triggers *FLOWERING LOCUS T* (*FT*) transcription in leaf vasculature. FT protein is then translocated to the SAM, where it binds to the b-ZIP transcription factor FD to form the florigen activation complex (FAC)^46,47^ and activates the transcription of genes such as *FUL* and *SOC1*^48^. The ageing pathway also contributes to the activation of these floral activator genes^49,50^. As plants age, expression of the developmental timer miR156 decreases^49,50^. This attenuates the inhibition of a set of *SQUAMOSA PROMOTER BINDING PROTEIN*-*LIKE* (*SPL*) genes, whose encoded TFs induce in turn the expression of *FUL*, *SOC1*, and *MIR172*. Under short-day (SD) conditions (8 h light and 16 h darkness), *SPL15* is particularly important, and floral transition occurs at a slower pace via the ageing pathway^49,51–53^. The combined effect of age and daylength signals thus modifies the network’s dynamics as they induce the transcription of *FUL*, *SOC1* and *MIR172* (Figure 1B, middle panel), which in turn repress *AP2* and *AP2-LIKE* genes transcriptionally^54,55^ and post-transcriptionally^56^. This mutual inhibition between floral activators and repressors defines an intermediate, transition phase. As floral transition progresses, the repression of floral inhibitors becomes more pronounced, thereby weakening the meristem’s vegetative identity and leading to the initiation of floral primordia and the flowering phase.

In this study, we combine experimental data and theory to elucidate the dynamics underlying the SAM transition from a vegetative to an inflorescence state. Specifically, we focus on determining whether bistability is involved in this process, and its dynamical consequences. We provide *in vivo* and *in silico* evidence that a time-dependent bistable switch, modelled as a toggle switch modulated by an age signal over time, underlies floral transition by explaining the temporal expression dynamics in wild type and mutants. We show that late-flowering phenotypes emerge naturally due to the influence of a ghost attractor, which explains long transients associated with disruptions to floral primordium development. Furthermore, we demonstrate that AP2 self-repression can cause fluctuations in its abundance at early stages. Collectively, our results suggest that floral transition at the SAM is driven by a time-dependent bistable switch mechanism, whereby different meristematic regions undergo distinct dynamic regimes.

## Results

### A time-dependent toggle-switch model for the floral transition

To investigate the floral transition mechanism at the SAM under the regulation of the photoperiodic and ageing pathways, we analysed the dynamics of the AP2 inhibitor and the SOC1 activator *in vivo*. We expanded previous reports^41,43,57,58^ by reanalysing existing data for AP2 and SOC1 expression^43^, and by acquiring new data for AP2 expression through imaging the SAM of *AP2::AP2:VENUS* LD-grown plants from 7 to 24LDs (Figures 2A, 2B, S1, and S2A). To improve accuracy, our new computational pipeline accounts for changes in meristem size by defining characteristic expression domains (see Methods and Figure S1). Consistently, both datasets showed a decrease in AP2 abundance over time until 17LDs, when SOC1 also reached its maximum (Figures 2, S2A, and S2D–S2F). From 17LDs onwards, the concentrations of both the activator and inhibitor stabilised (Figure 2C), and mature floral primordia began to appear on the meristem flanks (Figures 2A, 2B, S2B, and S2C).

**Figure 2.**
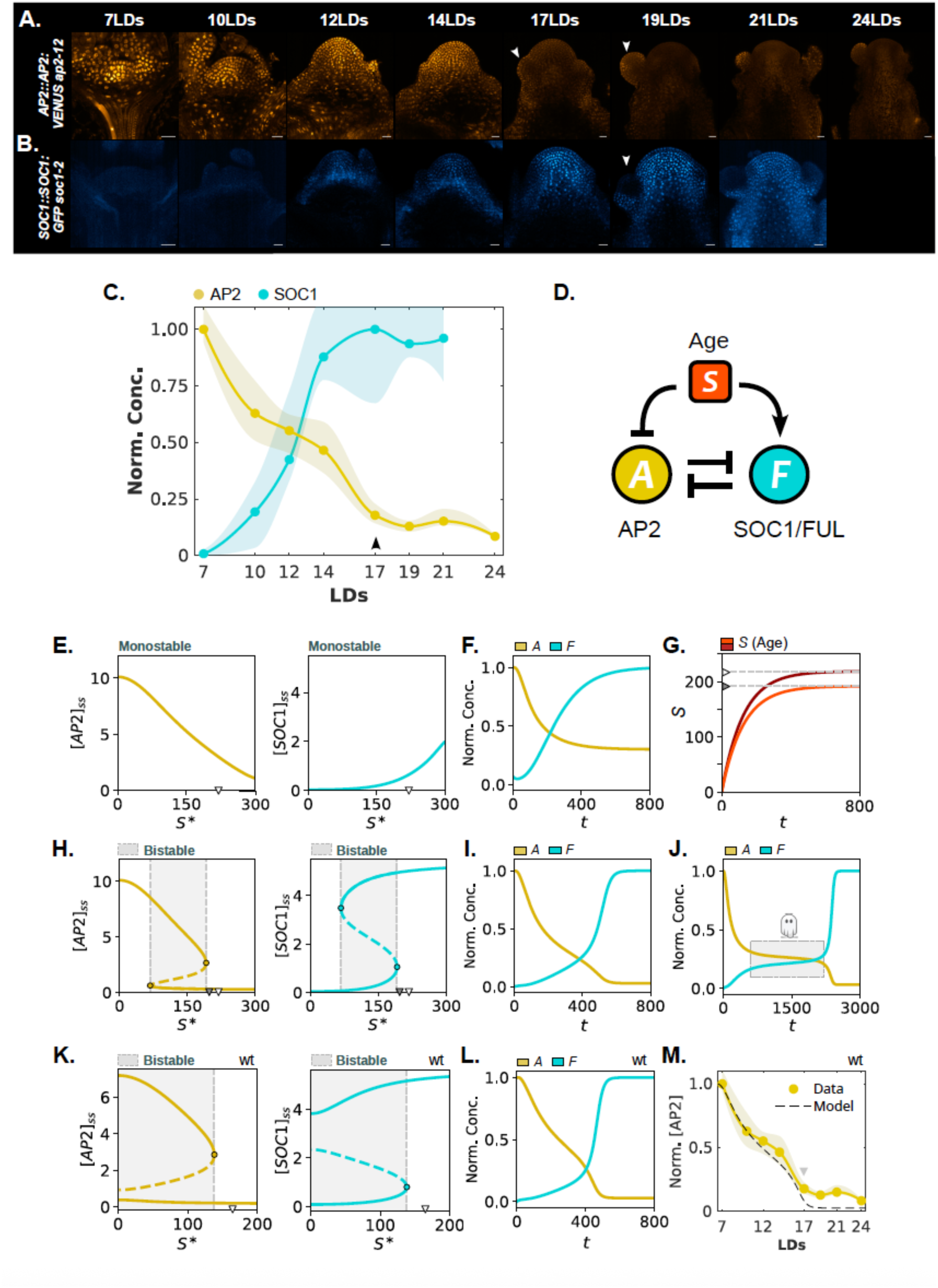
A time-dependent toggle-switch can recapitulate AP2 protein dynamics at the SAM in wt, indicating that time-dependent bistability underlies floral transition. A and B) Expression of AP2::AP2::VENUS (A) and SOC1::SOC1:GFP (B) at the SAM in *ap2-12* and *soc1*-2 backgrounds, respectively, grown under LDs. Scale bars = 20 µm. Arrowheads indicate visible floral primordia. C) Quantification of the temporal dynamics of AP2 (yellow) and SOC1 (cyan) effective concentrations (see Methods). Circles represent normalised median protein concentrations; shaded regions indicate the inter-quartile range (IQR). The arrowhead indicates floral transition completion (visible floral primordial, see Figure S6A). Fluorescence reporters are normalised by its maximum median value, respectively. D) Toggle-switch model network diagram for studying floral transition and the influence of environmental/endogenous signals. *A*: AP2; *F*: FUL and SOC1; *S*: age endogenous signal. E–M) Model analysis. E, H, and K) Bifurcation diagrams for *A* (yellow) and *F* (cyan) as a function of the age-signal (*S*\*) for (E) a monostable transition, (H) a bistable transition, and (K) a bistable transition representing the wt-genotype. Solid lines represent stable steady states, dashed lines represent unstable states. Triangles on the x-axes indicate the steady state value of *S* (*S_ss_*). Bistability occurs in the regions in grey. Coloured circles indicate saddle-node bifurcation points. F, I, J and L) Normalised numerical simulations showing the temporal dynamics of the activator (cyan) and inhibitor (yellow) (F) a monostable transition, associated to (E); (I, J) bistable transitions for two different parameter values of the age signal production, associated to (H), in which *S_ss_* is indicated with a white and black triangle on the x-axis for (I) and (J), respectively; and (L) a bistable transition representing wt, associated to (K). G) Age-signal (*S*) dynamics for (F, I) in red, and for J in dark red. In (J), the saturation of the age signal is closer to the saddle-node bifurcation, resulting in an extremely long transient due to the increased influence of the ghost attractor. M) Comparison between experimental (continuous yellow) and model (dashed black) results for AP2 dynamics. Both curves are individually normalised by their maximum and a temporal rescaling of the model simulation is used. See Suppl. Tables 1 and 3 for more information.

The developmental trajectories of the SAM from the vegetative to the inflorescence state can be inferred from the changes in AP2 and SOC1 concentration (Figure 2C). Here, we propose a time-dependent toggle-switch model^5,25^ to study the dynamics of the GRN underlying the floral transition and the effects of age and light signals (Figure 2D; see Methods). This single-compartment model consists of three ordinary differential equations that capture the mutual repression between floral activators (*SOC1*, *FUL; F* variable) and inhibitor (*AP2*; *A* variable) genes (see Methods). This double-negative feedback loop is influenced by a time-varying age signal, *S*, which is independent of *A* and *F*. The variable *S* serves as an external control parameter for this developmental transition, representing the physiological age of the plant. Biologically, *S* reflects the abundance of SPLs, which gradually increase over time as a result of the age-dependent decline of miR156^52^, a prerequisite for floral transition to proceed. In the model, *S* dynamics increases toward saturation, mirroring the mechanism by which SPLs activate *F* (*FUL* and *SOC1*) and *MIR172,* which in turn directly repress *A* (*AP2*)^40,57^. For simplicity, the influence of light in the system was introduced through the activator variable, distinguishing its contribution from that of the ageing pathway.

To analyse the behaviour of the model over time, we treated the age variable *S* as a control parameter, *S\**. In our framework, age represents the accumulation of SPLs at the SAM. Based on this model, we studied the corresponding phase portraits and bifurcation diagrams, whose stable states correspond to the valleys of the Waddington landscape (see Methods). In parallel, we performed numerical simulations of the temporal model dynamics to establish their correspondence with the bifurcation diagrams (Figures 2E–2J, S3, and S4; Methods).

Bifurcation analyses as a function of *S\** showed that a monostable transition with a single decreasing inhibitor stable branch and an increasing activator stable branch can take place (Figure 2E). This behaviour was consistent with the corresponding simulations, showing a smoothly decreasing inhibitor and increasing activator (Figures 2F and S4A). For other parameter values, bistable transitions were observed: bistability exists within a range of age values, and eventually the system becomes monostable, undergoing a saddle-node bifurcation and transitioning discontinuously from a vegetative (high repressor, low activator) to an inflorescence (high activator, low repressor) state (Figure 2H). The strength of both light and age signals determines the appearance of bistability (Figures 2H and S4C–S4H). Numerical simulations in the bistable scenario also showed consistent antagonistic dynamics between the activator and repressor variables (Figures 2I, 2J, S3D–S3I, and S4B). The discontinuity of the transition in the bistable case observed in the bifurcation diagrams (Figure 2H, S3D, and S3G) was not always evident in the numerical simulations, and only appeared for certain model parameters (Figures 2I, 2J, S3F, and S3I). When present, it was accompanied by long transients where the temporal dynamics are slowed down (Figures 2J, S3I, S4B-S4H, S5A, and S5B).

In the bistable case, the system’s dynamics continue to be transiently influenced by the initial state even after its disappearance via a ghost attractor^13–17^, leading to a critical slowing down^14–17^. This phenomenon can cause regulatory variables to remain at a transient concentration plateau, whose duration depends on the distance (Δ*S*) between the final steady state of the age variable (*S*_ss_) and the saddle-node bifurcation (*S_SN_*) (Figures S4C–S4H, S5A, and S5B). When critical slowing down is present, the duration of a transition follows a power law whose characteristic exponent is m = -½ (i.e., the duration of the transition follows *τ*^∝^Δ*S*^m^, with Δ*S* = *S*_ss_-*S_SN_*; Figure S5C). This behaviour is consistent with systems at criticality close to a saddle-node bifurcation, whereby an additional temporal scale in their dynamics appears, which follows a similar power-law behaviour^13–16^.

Model numerical simulations confirmed that higher AP2 production rates (*⍺*_1_) led to delayed floral transition (Figures S4G and S4H), consistent with late-flowering phenotypes observed in the inhibitor overexpression lines^43,57^.

### AP2 protein dynamics at the SAM are consistent with an underlying bistable mechanism during floral transition

To determine whether the experimental data were consistent with a monostable or bistable transition, we closely analysed the reduction in AP2 abundance at the SAM over time (Figure 2C). In the vegetative meristem, AP2 declined steeply between 7 and 10 LDs, but the reduction rate became significantly slower between 10 and 14LDs (Figures 2C and S2D–S2F). At this point (14LDs), leaf primordium formation ceases at the SAM tip, SAM elongation continues, and cauline leaves are detectable (Figures 2A, 2B, S2B, and S2C). Later on, at 17LDs, floral primordia start to appear. We hypothesised that this fast-slow-fast decline could be a signature of time-dependent bistability in the system (Figure 2I), where the slow dynamics result from the critical slowing down, consistent with the flat Waddington landscape after the vegetative state becomes destabilised (Figure 1B). This introduces an intermediate state with prolonged AP2 expression levels before floral primordia appear.

Simulations across the parameter space revealed that these dynamics could only be obtained in the bistable transition scenario (Figures S5D–S5F, see Methods). Although a monostable transition can generate progressively slower decay of the inhibitor, a subsequent re-acceleration was consistently absent (Figures S5D–S5F). Thus, the second rapid decay of the inhibitor is then most likely caused by the saddle-node bifurcation itself, when the system exits the ghost state and is attracted again by the inflorescence state. The discontinuous nature of this bifurcation would explain the measured second rapid reduction in the inhibitor (Figure 2C). Hence, a wild-type SAM is expected to be in a region of the parameter space where time-dependent bistability exists. As the age signal, *S*, increases, if age is treated as a parameter, the high inhibitor and low activator steady state concentrations smoothly decrease and increase, respectively (Figure 2K). Numerical simulation of system dynamics confirmed that the deceleration in AP2 reduction occurs exactly at the value predicted by the bifurcation analysis (Figures S3D, S3G, S5A, and S5B), and that AP2 is fully depleted only once the saddle-node is crossed, in conjunction with a completed floral transition.

For wt, numerical simulations qualitatively recapitulated the experimental data (Figures 2L, 2M, and S5G), showing an initial rapid reduction in AP2, followed by a transient, slower decreasing phase (10-14LDs Figure 2M), and subsequently by another strong reduction in AP2 expression (14-17LDs Figure 2M). Taken together, these data suggest that the slowdown and subsequent acceleration of AP2 reduction at the SAM constitute a distinct signature of bistability and a feature demonstrably absent in monostable transitions. This evidence provides compelling support for a bistable switch mechanism underlying floral transition.

### Single activator mutants exhibit delayed floral transition due to prolonged AP2 expression and the influence of the ghost attractor

To test the bistable nature of the floral transition, we studied how mutations in floral activator genes (*FUL* and *SOC1*), known to cause delayed flowering times^43,59,60^, affect the critical slowing down dynamics associated with the bistable transition. From confocal images, we quantified AP2 abundance at the SAM of *ful* mutant plants grown under LDs (Figure 3A). In addition, we re-analysed previously published data on AP2 expression in *soc1* mutants^43^ using our new computational pipeline (Figure S1). Confocal images of AP2 translational reporter lines in both *ful* and *soc1* demonstrated that AP2 expression at the SAM persists longer than in wt under LDs, consistent with their delayed flowering phenotypes (Figures 3A and 3B). In *ful,* the decrease in AP2 expression was very similar to wt, although complete AP2 depletion was slightly delayed compared to wt (19LDs in *ful* versus 17LDs in wt, Figure 3C), consistent with its slight late-flowering phenotype under LDs^59^. These results contrast with those of *soc1*, in which AP2 expression persists much longer than in wt (depletion of AP2 in *soc1* occurs at 24LDs versus 17LDs in wt; Figure 3C), in agreement also with the late-flowering phenotype of *soc1* compared with *ful* and wt under LDs^43^. Despite a similar initial decline in expression, the reduction in AP2 at the SAM in *soc1* ceases from 17LDs until 21LDs, and only at 24LDs is AP2 completely depleted from the SAM (Figure 3C). This prolonged expression of AP2 in *soc1* mutants directly correlated with severely delayed floral primordia development, a robust proxy for floral transition completion, with visible floral primordia appearing only from 24LDs onwards (Figures 3B, S6A, and S6B). Such an extremely late formation of floral primordia was thus preceded by the quantified plateau phase in decline of AP2 expression (Figure 3C). These data collectively suggest that AP2 depletion at the SAM is necessary for floral primordium formation.

**Figure 3.**
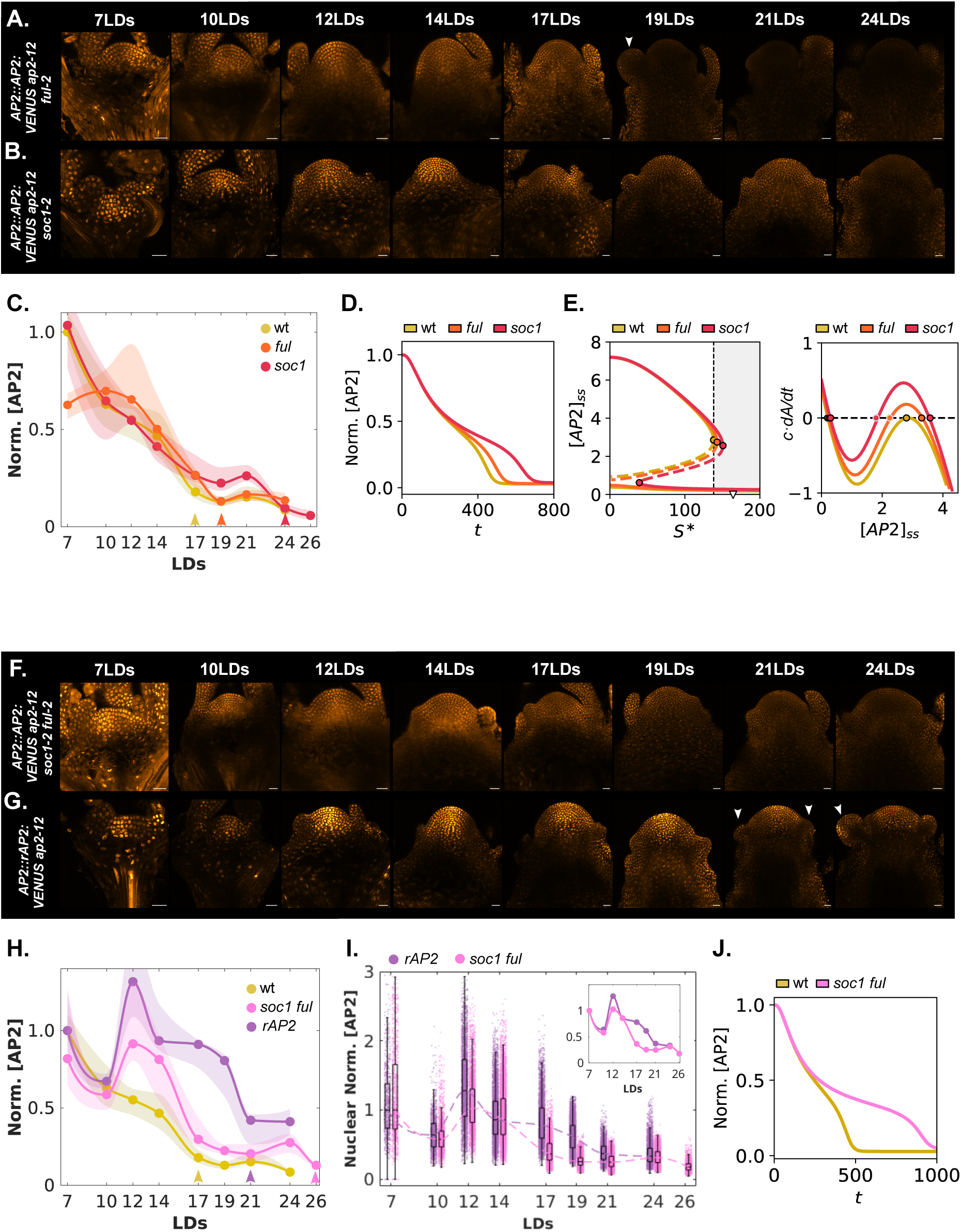
Mutations of floral activator genes prolong the transient-expressing phase of the inhibitor and reveal its fluctuating behaviour. A and B) Expression pattern of AP2:AP2::VENUS in (A) *ap2*-*12 ful-2* and (B) *ap2*-*12 soc1*-2 at the SAM of plants grown under LDs. Scale bars = 20 µm. C) Quantification of the temporal dynamics of AP2 concentration at the SAM in wt (yellow), *ful* (orange), and *soc1* (red). Circles and shaded regions are defined as in Figure 2C. Arrows indicate floral transition completion (visible floral primordia, see Figure S6). D) Normalised numerical simulations for wt, *ful*, and *soc1* showing the *AP2* model variable dynamics. E) AP2-age bifurcation diagram (left) and nullclines (right) for wt, *ful,* and *soc1*. The bifurcation diagram is represented as in Figure 2. F and G) Expression pattern of (F) AP2:AP2::VENUS in *ap2*-*12 soc1-2 ful-2* and (G) AP2::rAP2:VENUS *ap2-12* at the SAM under LDs. Scale bars = 20 µm. H) Quantification of the temporal dynamics of AP2 concentrations at the SAM in wt (yellow), *soc1 ful* (pink), and *rAP2* (purple). I) Single-cell nuclear quantification of AP2 protein expression in *soc1 ful* (pink) and *rAP2* (purple) at the SAM during floral transition, for the same SAMs analysed in (H). The inset shows the median trends. J) Normalised AP2 model temporal dynamics for wt (yellow) and *soc1 ful* (pink). See Suppl. Table 1 for parameter values. In (C) and (H), fluorescence reporters are normalised by the maximum median value in wt, except for *rAP2*, which is individually normalised with respect to the median at 7LDs due to a different transgene insertion. In (I), single-cell nuclear concentrations in *soc1 ful* and *rAP2* are normalised individually by the maximum median value and the median value at 7LDs, respectively. See Suppl. Tables 1 and 3 for more information about parameters and normalisations.

Numerical simulations of the *ful* and *soc1* mutants, modelled by reduced expression rates in the activator variable, qualitatively recapitulated the AP2 decline shown in the experimental data (Figures 3C and 3D, and S6C–S6E). In these mutants, a stronger age signal (higher SPL levels) was required to cross the saddle-node bifurcation and perform the vegetative-to-inflorescence transition at the SAM (Figure 3E). This shift in the position of the saddle-node bifurcation amplifies the influence of the ghost attractor, naturally explaining the delayed floral transition observed in both mutants (more pronounced in *soc1*) and the longer plateau in the AP2 decline observed in *soc1* (17-21 LDs, Figure 3C), consistent with the critical slowing down predicted by the model.

To quantify the presence of a second timescale during the transition, we performed an analytical fit of AP2-VENUS dynamics in the wt and *soc1* mutant (Figure S7 and Suppl. Tables 5 and 6) using a sum of two arctangent functions. This approach, which is independent of the model, confirmed that both wt and *soc1*-dynamics exhibit a “bottleneck” phase, which is a signature of critical slowing down^11,12^, and that this phase is significantly accentuated in the *soc1* mutant. Specifically, the duration of the critical slowing down (Δ*τ*) more than doubled in *soc1* and the minimum speed of AP2 reduction (∣v_min_∣) was reduced by more than 50% compared with wt (Figures S7H and S7H). This second timescale, which appears due to criticality, provides a basis for the prolonged AP2 expression observed experimentally (Figure 3C).

Prolonged transient dynamics, linked to impaired development of floral primordia in late-flowering mutants such as *soc1*, are therefore naturally derived from the influence of the ghost attractor. Taken together, these results support the hypothesis of floral transition proceeding via a bistable-switch mechanism.

### *soc1 ful* double activator mutants and plants carrying an *AP2* transgene resistant to miR172 display fluctuations in AP2 levels in the vegetative meristem

To further address our hypothesis, the dynamics of AP2 expression were assessed under LDs in two extreme genotypes: the *soc1 ful* double mutant and an *AP2::rAP2:VENUS ap2-12* transgenic line that expresses a version of AP2 resistant to miR172 (*rAP2*) (Figures 4F–4I). In *soc1 ful*, AP2 depletion at the SAM was even more delayed compared with that in single mutants (AP2 complete depletion only occurred at 26LDs in *soc1 ful*; Figures 3F, 3H, S6H, and S6I), consistent with its extremely late-flowering phenotype^60,61^. Similarly to *soc1*, *soc1 ful* showed a long plateau phase in AP2 reduction between 17 and 24LDs (Figures 3F, 3H, S6H, and S6I), which was also associated with a severe delay in floral primordium initiation (Figures 3F, S6F, and S6G). AP2 was depleted from the SAM at 26LDs (Figures 3F, 3H, and S6H–S6J), and floral primordia were only observed at 29LDs (Figures S6F, S6G, and S6J). This extended plateau in AP2 expression before complete depletion of the inhibitor was reflected in the numerical simulations of *soc1 ful* (Figure 3J). In this genotype, the bistable region expands towards increased age-signal values, which reduces the gap between the saddle-node point and the final age value, thereby amplifying the influence of the ghost attractor (Figure S8A and S8B). Consequently, the simulated *soc1* and *soc1 ful* mutants show a clear increase in the time for which the system is trapped by the ghost attractor (*τ*_g_) compared with the time needed to reach the saddle-node bifurcation (*τ*_0_) in wt or *ful* (Figures S8C–S8F), as well as a further decrease in the predicted rate of reduction in AP2 (Figure S8G).

**Figure 4.**
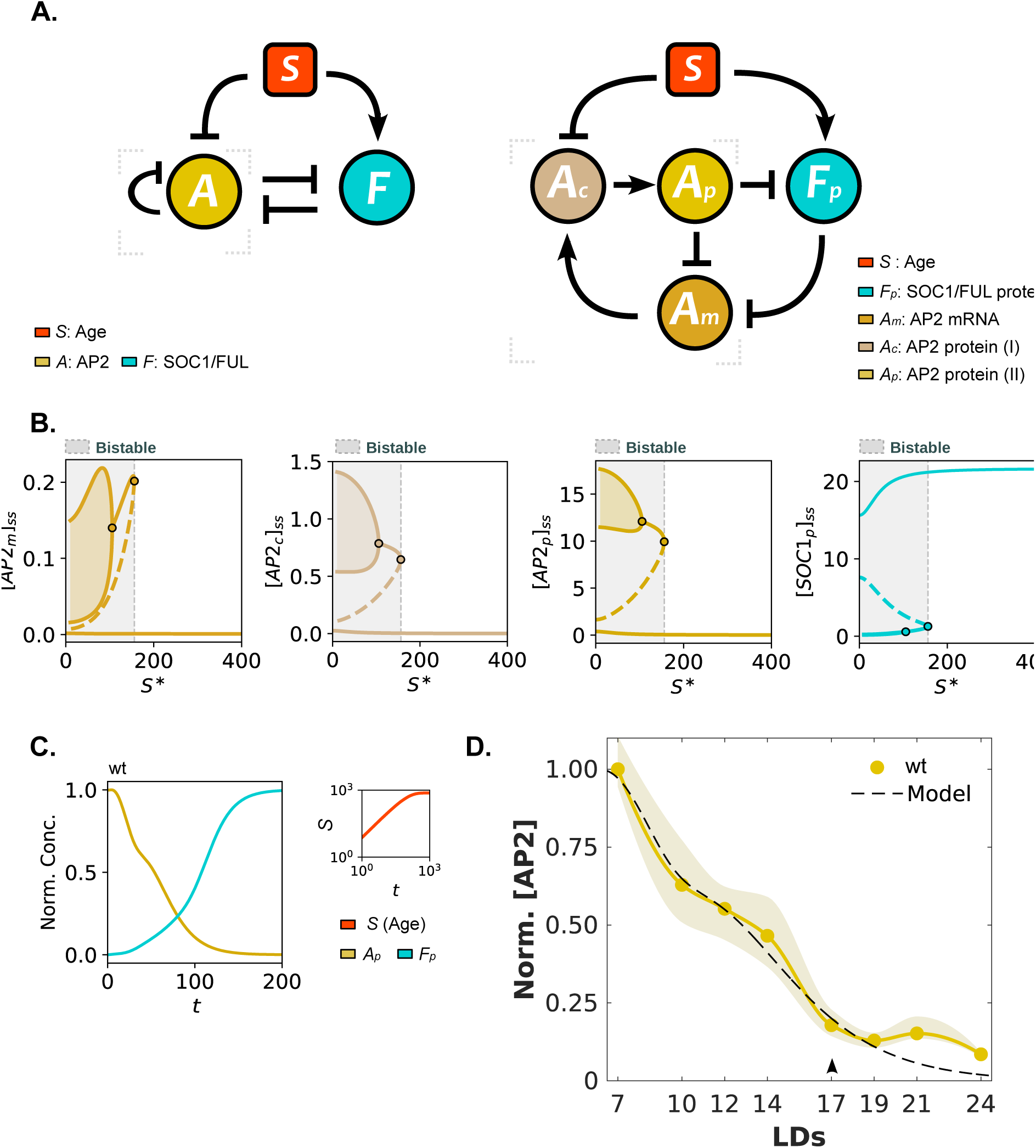
An extended toggle-switch model incorporating AP2 autoinhibition driving oscillations can still recapitulate the wt-dynamics during floral transition. A) Extended toggle-switch network-model diagram incorporating AP2-autoinhibition (left) and a detailed version of the diagram (right), introducing new variables to describe the AP2 autoinhibition, to ensure sufficient time delay to drive oscillations. B) Bifurcation diagrams for model variables *Am* (orange-brown), *Ac* (light brown), *Ap* (dark yellow) and *Fp* (cyan) as a function of the age-associated signal (*S*\*) for wt. Bifurcation diagrams are represented as in Figure 2; here, circles represent saddle-node and Hopf bifurcations. C) Numerical simulations for wt showing the temporal dynamics of the activator (Fp, cyan) and inhibitor (Ap, yellow) proteins, individually normalised by its maximum. *S* dynamics are in red in the top-right corner. D) Comparison between experimental (yellow) and the extended model’s (dashed black) results for AP2 expression at the SAM. Both curves are individually normalised by their maximum and a temporal rescaling of the model simulation is introduced for comparison. See Suppl. Tables 1 and 3 for more information about parameters and normalisations.

To validate these theoretical predictions, we analysed the experimental AP2 dynamics across all genotypes by calculating the derivative of the interpolated trajectories and the discrete ratios (Δ[AP2]/Δt) between consecutive points. Similarly to the analytical arctangent-fit results for *soc1* (Figure S7), this analysis confirmed both the significantly lower decline velocity in this stagnation period (Figures S8H–S8K) and the corresponding increase in the time the system is trapped by the ghost attractor (Figures S8L and S8M) in activator mutants compared with wt. Collectively, these results demonstrate the emergence of a dominant second timescale through the critical slowing down phenomenon. Furthermore, they strengthen the hypothesis of time-dependent bistability in the system and underscore the key role of the ghost attractor in explaining the long transient dynamics observed in *soc1 ful*.

In *rAP2*, AP2 expression at the SAM effectively declined at 21-24LDs, but persisted even when mature floral primordia were clearly visible (Figures 3G, 3H, S6F, and S6G). Notably, rAP2 maintained comparatively high AP2 abundance even at later stages of the floral transition (17-19LDs in Figures 3H, S6H, and S6I). To further understand the rAP2 dynamics, we compared rAP2 expression levels across different SAM domains, such as the CLV3-expressing central zone (CZ) and the peripheral zone (PZ), where new primordia are initiated. We found that rAP2 signal intensity at the adjacent region of newly formed primordia was consistently lower than that at the central region (Figures 3G, S10A–S10C). These preliminary observations suggest that the PZ region undergoes the strongest reduction in AP2 levels during the floral transition, potentially facilitating floral primordium initiation despite high overall AP2 expression in the CZ. Such a spatial gradient suggests a distinct mode of action of AP2 in different regions of the SAM.

Both *soc1 ful* and *rAP2* genotypes, however, exhibited a feature in their AP2 dynamics during early vegetative stages (7–12LDs) that was very different from that in wt and single mutants (Figures 3C and 3H). In *rAP2* and *soc1 ful*, AP2 protein levels increased significantly between 10 and 12 LDs, and only decreased again at 12LDs (Figure 3H). To confirm that this fluctuating behaviour between 10 and 14 LDs was not due to an artifact of the quantification pipeline or an uncontrolled event, we quantified single-cell nuclear AP2 concentration in SAMs from both genotypes during floral transition (Figures 3I, S10D–S10F, S11, and S12). Single-cell quantification confirmed the overall increase in AP2 expression between 10 and 12 LDs in *soc1 ful* and *rAP2* genotypes (Figures 3I, S11D, S11E, S12D, and S12E) and the absence of fluctuations in AP2 levels at early vegetative stages in wt (Figures S11A and S12A) and single activator mutants (Figures S11B, S11C, S12B, and S12C). However, these initial fluctuations in AP2 dynamics could not be replicated in *rAP2* or *soc1 ful* numerical simulations in our model (Figures 3J and S8B). This discrepancy raised the question: how can this fluctuating behaviour be theoretically generated, and which underlying biological interactions could explain it?

### A negative feedback loop in AP2 resolves the fluctuating behaviour and still recapitulates wt dynamics

A negative feedback loop with delay is sufficient to generate oscillations in a system^11,62–64^. Based on this principle, we hypothesised that autoinhibition in the repressor variable could explain the fluctuating behaviour in AP2 expression dynamics observed in *soc1 ful* and *rAP2* backgrounds (Figure 4A). Indeed, AP2 protein can bind to the *AP2* gene^65,66^, and its self-repression has been proposed^44,55,65,66^. However, the physiological relevance of this regulation remains unknown. We propose that the influence of this autoinhibition on the system’s dynamics is attenuated in wt under LDs due to the strong repression of AP2 by both miR172 and FUL/SOC1, and therefore becomes relevant only in deregulated contexts in which AP2 dynamics are severely impaired, such as in *soc1 ful* or *rAP2* backgrounds.

The delay in AP2 self-repression was implemented in the model using three variables mediating the feedback: *AP2* mRNA (*Am*), an intermediate AP2 protein state (*Ac*), and the final AP2 transcription factor (*Ap*), which represses its own transcription and SOC1-FUL variable (*Fp*) (Figure 4A; see Methods). As a result of this autoinhibition in the repressor, oscillations emerged in the system for certain values of the age signal, both in a monostable and a bistable scenario (Figures S13A–S13G) due to the appearance of a stable limit cycle^11,12^, which is an attractor that presents sustained oscillations. In the bistable scenario, the stable limit cycle can coexist with the stable inflorescence attractor (Figures 4B, S13A–S13C, S13F, and S13G). As age increases, the limit cycle eventually becomes a non-oscillatory vegetative attractor through a Hopf bifurcation, followed by a saddle-node bifurcation (Figure 4B). The amplitude of these oscillations, determined by the amplitude of the limit cycle, is significantly higher in the repressor variables (*Am*, *Ac* and *Ap*) than in the activator variable (Figure 4B), as determined by the initial low concentration of this variable in a vegetative SAM.

Can oscillations function as an alternative mechanism to recapitulate the experimentally observed AP2-wt dynamics? To answer this question, we first examined with numerical simulations whether monostable transitions having a limit cycle –with or without a Hopf bifurcation– could recover the slowing down and re-acceleration of the AP2 decay. Our simulations in those cases showed either sustained or damped oscillations (Figures S13D, S13E, S13H, and S13I). In a monostable scenario with a Hopf bifurcation, damped oscillations turned into a non-oscillatory behaviour after a short period of time for very high age signal steady state values (which led to a very rapid age increase; *S_ss_* = 2000 in Figures S11H and S11I). Conversely, oscillatory behaviour could easily be prevented in the presence of bistability, as the age signal steady state increased (Figures S13J and S13K). Noticeably, although a monostable transition with a Hopf bifurcation could emulate the wt AP2 dynamics for high age steady state values (Figure S13H), this scenario failed to recapitulate the dynamics of the activator mutants that were experimentally measured, such as their late flowering phenotypes, the increase in length of AP2 stagnation or the fluctuations in the inhibitor in *soc1 ful* (Figures S14A and S14B). This lack of flexibility in the monostable scenario contrasts with the variety of dynamic behaviours yielded by a bistable mechanism (Figures S13J and S13K).

Consequently, the wt-genotype in this extended model remains within a bistable transition parametric region (Figure 4B). Only this transition simultaneously recovers its fast–slow–fast dynamics and enables the possibility to generate oscillations detected in other genotypes for other parameter values (Figures S13J and S13K). Numerical simulations showed opposite non-oscillatory temporal dynamics in both the activator (*Fp*) and the active form of the inhibitor (*Ap*) and recapitulated the wt behaviour (Figure 4C and 4D). Consistent with the experiments, the effect of AP2 self-repression in wt was attenuated. The slowing down observed in the decay of the inhibitor *Ap* continues to reflect the influence of the ghost attractor (*Ap* ⪞ 0.5 in Figure 4C). Overall, the extended model shows a good qualitative agreement between theory and experiments for wt (Figure 4D; differences from 19LDs onwards are attributed to the background AP2 signal in the experiments). These results also demonstrate the context-dependent relevance of AP2 autoinhibition, key in *soc1 ful* and *rAP2* and minimised in wt and single mutants, highlighting the role of bistability in providing robustness necessary to suppress oscillations during the transition in wt.

### AP2 dynamics during floral transition in different genotypes can be explained by the extended toggle-switch model

The dynamics of AP2 expression in different mutant genotypes were assessed analytically and numerically with the extended model, using the previously modelled wt genotype as reference (see Methods and Figure 5). Figure 5A shows the position of each genotype as a function of the inhibitor (*A_m_*) and activator (*F_p_*) basal production rates (α_1_ and α_0_, respectively). For each pair of parameters (α_0_, α_1_), we computed the dynamical regimes crossed as the age signal variable increases to the steady state reached in the numerical simulations. The resulting diagram is coloured to distinguish regions by crossed regimes (B: bistable, M: monostable), their order, and the presence or absence of a limit cycle (oscillations). The reduction in the activator expression rate to emulate floral activator mutants shifts the system in the parameter space towards the M-B-M region (Figure 5A), moving the Hopf and saddle-node bifurcation towards higher age-signal values (Figure 5B). This shift has two consequences in the temporal dynamics: it expands the range of the stable limit cycle and extends the upper branch of the saddle-node after the Hopf bifurcation, increasing the likelihood of observing oscillations and the influence of the ghost attractor (Figures 5B and S14C–S14G).

**Figure 5.**
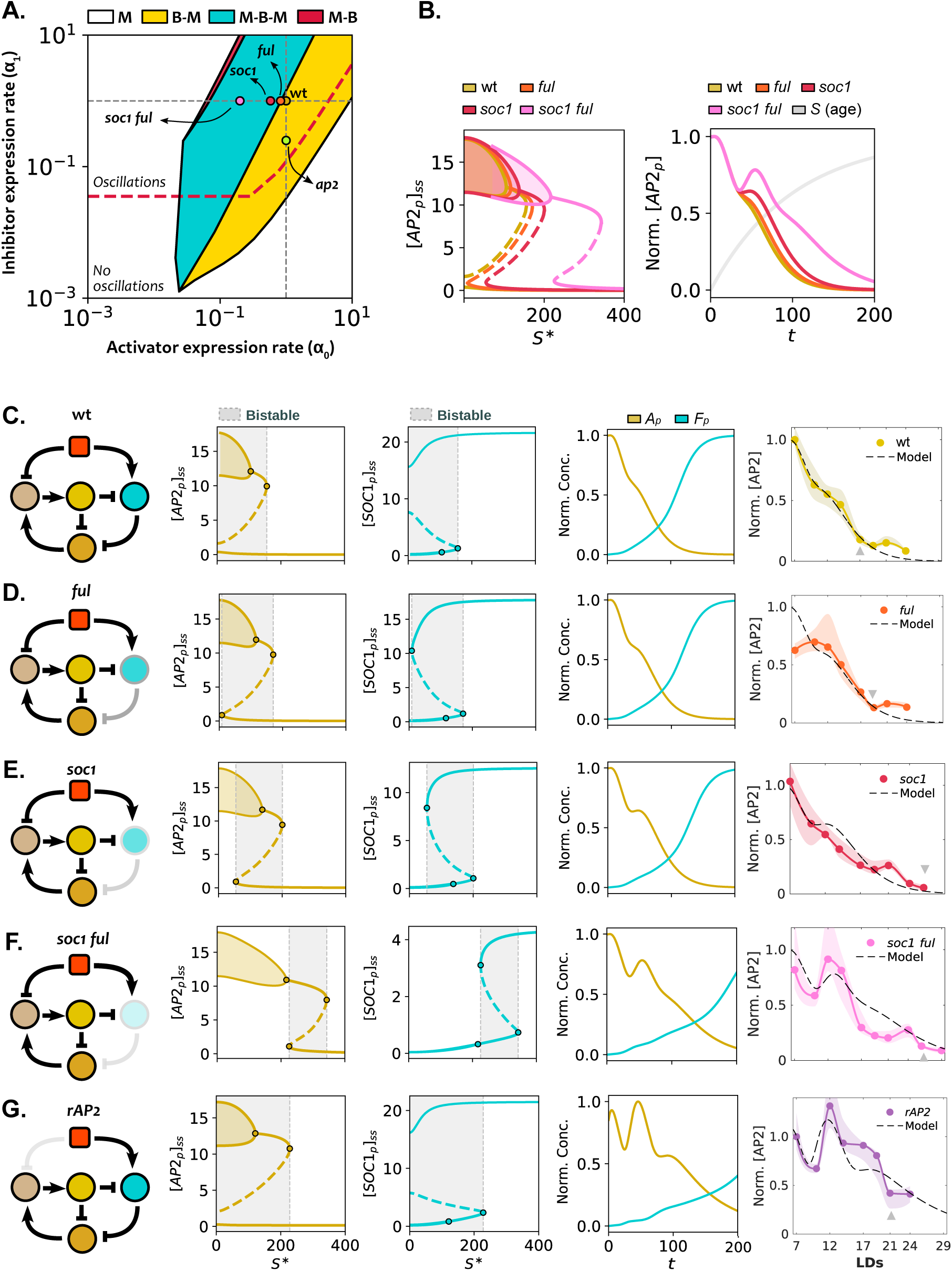
The extended model explains genotype-specific dynamics and qualitative features of floral transition. A) Stability diagram illustrating the stability regimes (M: monostable, B: bistable) crossed for a given (*⍺*_0_,*⍺*_1_) parameter pair (representing *Fp* and *Ap* expression rates, respectively) across the *S*\* range [0, 750]. Genotypes are represented by coloured dots. Stability regimes include: M, where the system is always monostable; B-M, where the system starts bistable and transitions to monostable; M-B-M, involving two saddle-node bifurcations; and M-B, where *S\** is insufficient to reach the second saddle-node. The red line marks the boundary above which oscillations occur. B) *Ap* bifurcation diagrams (left) for wt, *ful*, *soc1,* and *soc1 ful* as a function of *S*\*. Normalised numerical simulations (right) show AP2 and *S* dynamics across genotypes. Curves are normalised individually. C–G) Comparison between the modelled genotypes and the experimental data for (C) wt, (D) *ful,* (E) *soc1*, (F) *soc1 ful,* and (G) *rAP2*. Network diagrams (first column), bifurcation diagrams (second and third columns), numerical simulations reached for *S_ss_* = 750 (fourth column), and comparison between experimental (solid line) and simulated (dashed black line) AP2 dynamics (fifth column). Both curves are normalised by the maximum value reached in wt and a temporal rescaling of the model simulation is used. For *rAP2*, experimental data and model simulations are normalised by their initial values. See Suppl. Tables 1 and 3 for more information about parameters and normalisations. Bifurcation diagrams are represented as in Figure 4B.

Numerical simulations consistently reproduced the delayed AP2 reduction and late-flowering phenotypes of the single mutants (Figures 5C–5E; the inflorescence SAMs appear at 17LDs, 17-19LDs, and 24LDs for wt, *ful* and *soc1,* respectively). The simulated *ful* mutant exhibited a more pronounced plateau during AP2 reduction (Figures 5B and 5D), which was further accentuated in the *soc1* mutant (Figures 5B and 5E).

In the *soc1 ful* double mutant, these features were amplified, resulting in a very high age-signal requirement to reach the saddle-node bifurcation and a remarkably long phase with oscillations (Figures 5B, 5F, and S14E). Numerical simulations of the double mutant effectively captured both the fluctuating behaviour of AP2 decline in a vegetative SAM and its extreme late-flowering phenotype (Figures 5B, 5F, and S14E). These dynamics were also recapitulated in the simulated *rAP2* genotype, in which an initial oscillation in AP2 levels is followed by a plateau at very high inhibitor values (Figure 5G). Such a plateau at high AP2 expression in *rAP2* is possible due to the absence of the repression of this inhibitor by the age signal (see Methods and Figure 5G).

To assess the robustness of the bistable mechanism underlying floral transition, we performed stochastic numerical simulations for each genotype under both intrinsic (Figures S15 and S16) and extrinsic noise (Figure S17 and S18) to model molecular and environmental fluctuations, respectively^67^. Overall, the temporal dynamics of model variables recapitulated the behaviour of the associated deterministic simulations, confirming that the proposed bistable switch and AP2 autoinhibition mechanism are robust to molecular and environmental fluctuations. As expected, increasing noise intensity resulted in greater variability across all genotypes (Figures S15, S16, S17 and S18) and in simulations in which the age variable saturated closer to the saddle node bifurcation (Figure S19). On this second analysis, increased noise effectively destabilised the ghost attractor and led to earlier transitions from vegetative to inflorescence states (see median curves in Figures S19A and S19B), illustrating how stochastic fluctuations modulate transient dynamics at criticality, near the saddle node^67^. Furthermore, the quartile coefficient of variation (QCV) identified extended periods of maximum variability for AP2p and SOC1p variables (Figure S19C), showing that, when stochasticity is taken into account, variability across a population can be amplified at criticality^68^.

These results demonstrate that the time-dependent bistable switch model with autoinhibition can better recapitulate the different behaviours across all studied genotypes (Figures 5 and S14F-S14K). The absence of different dynamic behaviours across the numerically simulated genotypes in the monostable scenario highlights the fundamental role of bistability in floral transition. Bistability ensures flexibility of the system upon perturbations via the expanding range of possible dynamical behaviours. It is this bistability that allows the new model to simultaneously account for the stagnation and late flowering phenotypes in wt and single floral activator mutants, and also the oscillatory regime in *soc1 ful* and *rAP2* genotypes, underscoring the relevance of AP2 autoinhibition in contexts in which this inhibitor is severely deregulated.

### Spatial modelling reveals distinct dynamic regimes along the SAM

AP2 and SOC1 expression patterns also exhibit spatial heterogeneity within the SAM during floral transition (Figure 2). To accurately capture and analyse this phenomenon, we quantified their spatial patterns of expression throughout the transition (Figures 6A, S20, and S21A; see Methods). At early stages (7-12 LDs), AP2 protein was strongly expressed near the apex (Figures 2A, 6A, and S20A). By contrast, SOC1 expression initiated at a lower SAM position (10LDs), and progressively expanded towards the apex (17 and 19LDs; Figures 2B, 6A, and S20B). Notably, a double peak of expression along the longitudinal axis of the SAM was observed in the SOC1 spatial expression profile at 14LDs (Figures 6A and S20E). AP2 spatial profiles were also quantified in *ful*, *soc1*, *soc1 ful* and *rAP2* genotypes (Figures S20C and S20D). A longer persistence of the inhibitor was observed in all of them within the meristem (Figures S20C and S20D), as well as initial variations in AP2 levels in the meristem in *soc1 ful* and *rAP2* (Figure S20D), as previously described when analysing the global temporal dynamics of the SAM (Figure 3). We therefore investigated whether this observed spatiotemporal heterogeneity in AP2 and SOC1 wt-expression patterns indicates distinct meristematic regions undergoing different dynamic regimes. To test this, we extended the previous single-compartment model into a 1D spatially extended and continuous model representing the longitudinal axis of the SAM (Figure 6).

**Figure 6.**
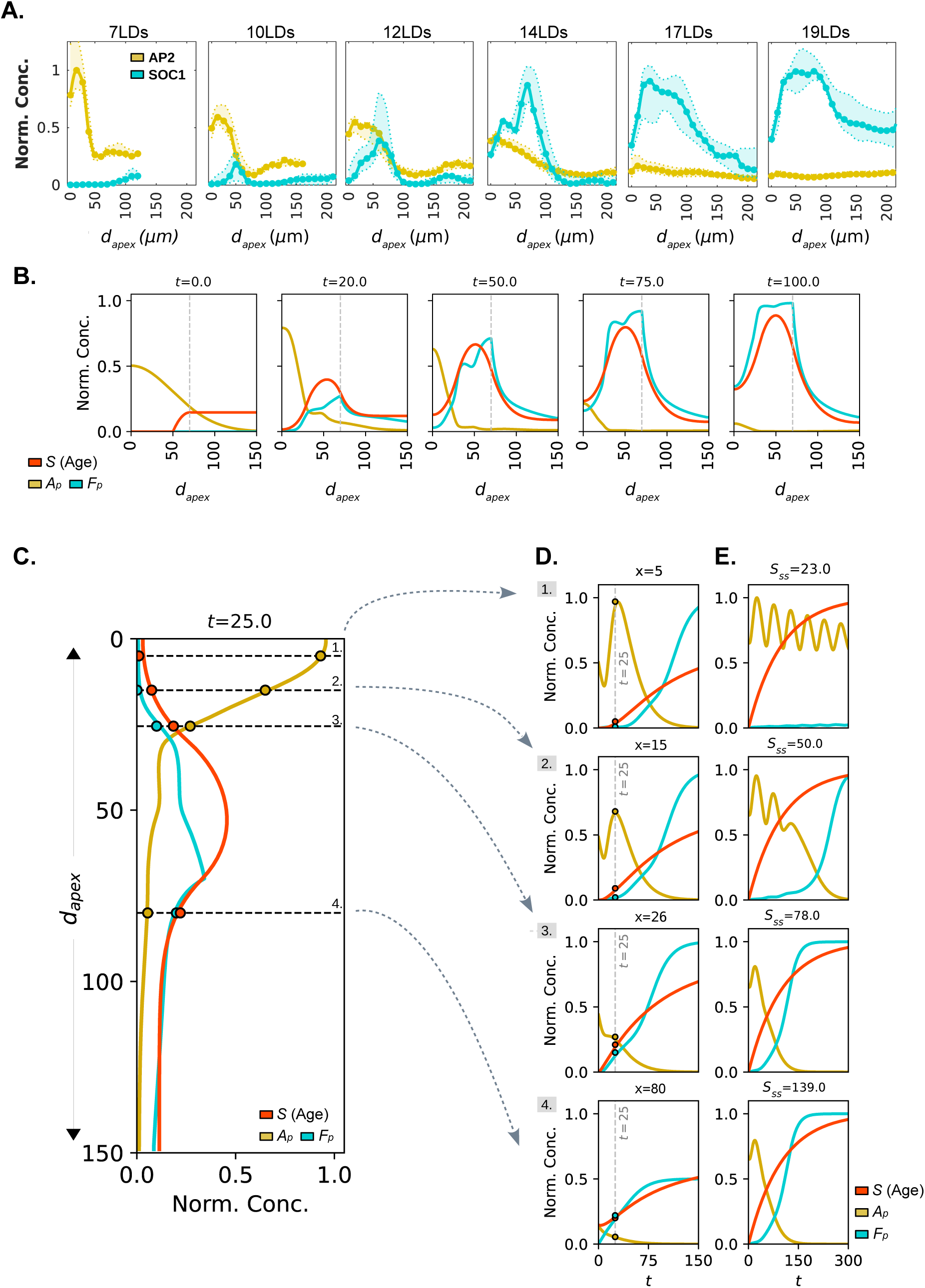
A spatially extended model reveals different dynamical scenarios within the SAM during floral transition. A) Temporal and spatial dynamics of normalised AP2 (yellow) and SOC1 (cyan) expression during floral transition at the SAM in wt. *d_apex_* represents the distance to the SAM apex. Fluorescence reporter concentrations are normalised by its global maximum median value reached along the simulation, respectively. B) Numerical simulations for *Ap*, *Fp,* and *S* wt variables along one spatial dimension at different time points. Only diffusion of the age signal, *S*, has been added to the model analysed in Figures 4 and 5. C, D) Extracted spatial (C) and (D) temporal profiles from (B), showing *Ap*, *Fp,* and *S*. In (C), spatial profiles as a function of the distance to the SAM apex are normalised by their global maximum. In (D), temporal dynamics are shown at different positions along SAM’s longitudinal axis, and coloured dots correspond to those in (C). E) Extracted temporal profiles at different positions along SAM’s longitudinal axis from the spatial model for different *S_SS_*. *Ap* and *Fp* are normalised by their maximum values across the four scenarios displayed; *S* has been normalised within each scenario. The modelled spatial and concentration variables are in arbitrary units. See information about parameters and equations in Methods and in Suppl. Table 1.

The model equations incorporated a diffusion term for the age signal S (see Methods), which is biologically supported by the movement of small microRNAs, such as miR172, through plasmodesmata^40,69^. Accordingly, deeper SAM meristematic regions were initialised with higher age signal levels (Figure S21B), and age signal production was assumed to occur only upon reaching a specific age signal concentration threshold. Although the SAM continuously grows during floral transition, explicit growth or cell division was not included for simplicity. To replicate the attenuation of reporter signals at distant regions from the SAM apex, the expression rates of the activator and inhibitor (*Ap* and *Fp*) were progressively reduced as a function of the distance to the apex (see Methods).

Numerical simulations of the spatial model demonstrated strong qualitative agreement with experimental data. The activator profile emerged close to the apex, subsequently displaying two distinct peaks, recapitulating the double peak found in SOC1 expression (10–12LDs and t=20 in Figures 6A and 6B). Only at later time points was the activator expression profile flattened near the apex, as measured in experiments (Figure 6B). The double peak in the activator is consistent with a coherent feedforward loop motif^70^, in which the age-signal activates directly and indirectly (by repressing AP2) the activator, and the differing strengths of these two activation mechanisms in space generate the two peaks along the SAM longitudinal axis. Theoretical analysis of the spatial model revealed that a non-homogeneous spatiotemporal distribution of the age signal confers distinct dynamic properties to different meristematic regions (Figures 6C–6E and S21B). Regions near the apex initially exhibit transient oscillatory behaviour (section 1 in Figures 6C–6E and S21). As the bifurcation age parameter, *S\**, increases over time, this transient oscillatory phase terminates via a Hopf bifurcation (Figures S21C and S21D). Sustained oscillations would only occur if the final variable value of *S* in the simulations remained within the oscillatory region of the bifurcation diagram. Figure 6E summarises these distinct scenarios comprehensively. For regions farther from the apex, higher age signal levels reduce the likelihood of observing oscillations. This causes the system to pass earlier through the non-oscillatory branch with high AP2 levels after the Hopf bifurcation, and to undergo the saddle-node bifurcation earlier (section 2 in Figures 6C–6E, S21C, and S21D). At even greater distances from the apex, the initial age signal concentration positions the system within a monostable region (sections 3 and 4 in Figures 6C–6E, and S21).

This spatial version of the toggle-switch model with AP2 self-inhibition exhibits qualitative behaviours that are highly consistent with experimental observations, highlighting the robustness and flexibility of the model. Importantly, these results suggest that different meristematic regions can undergo distinct dynamic regimes during floral transition as a consequence of time-dependent bistability influenced by spatial gradients of the age signal.

## Discussion

How biological systems transition between distinct developmental states has long intrigued scientists since Waddington’s seminal work. Although this question has often been studied in the context of cell signalling and differentiation, it has remained, with a few exceptions^32–36^, largely unexplored in plant development. In this context, plant developmental transitions have been conceptually described as shifts between attractors in a dynamical landscape, where the temporal modulation of regulatory parameters ensures robustness of the developmental process^30^. Here, we provide evidence for a time-dependent bistable mechanism underlying the floral transition, whereby the SAM changes between vegetative and inflorescence states. To formulate this model, we comprehensively quantified and analysed the spatiotemporal dynamics of key regulatory elements within the GRN that regulates floral transition at the SAM in response to both the ageing and photoperiodic pathways.

Our analysis of AP2 and SOC1 temporal dynamics consistently supports a time-dependent bistable mechanism underlying floral transition. A toggle-switch model modulated over time by an age signal revealed that the characteristic slowing-down of AP2 reduction measured in wt is the theoretical signature of a bistable transition (Figure 2). This deceleration is caused by the influence of a ghost attractor, which introduces a temporal scale in the transition^13,14,16,21,71^. This phenomenon directly accounts for the prolonged AP2 expression quantified in floral activator mutants (*ful*, *soc1,* and *soc1 ful*). The remarkable correlation between these long transient plateaus in AP2 reduction and the severe impairments in floral primordium development makes the morphological implications of this second timescale visible (Figures 3A–3E). Furthermore, the incorporation of AP2 self-repression in the extended model, supported by previous studies^44,55,65,66^, recapitulated the fluctuations in AP2 levels at early stages in *soc1 fu*l and *rAP2* genotypes (Figures 3H–3J, 5F, 5G, S11, and S12). The model demonstrates that a bistable transition is necessary to ensure the broad range of distinct dynamical behaviours in the system and to provide flexibility upon genetic perturbation, a property not easily achieved with a monostable transition. The broad range of behaviours is also due to the coexistence between a non-oscillatory and an oscillatory attractor^72^, a kind of bistability recently proposed in animal development modelling^64^. But importantly, the critical slowing down regime itself confers dynamic flexibility to prolong the transition, which could be used in certain environmental conditions.

For environmentally sensitive systems such as plants, the mechanism of a developmental transition can determine susceptibility or robustness to environmental perturbations. A bistable-switch-like floral transition could thus explain how the SAM maintains its commitment to the inflorescence state, and does not return to a vegetative state, even under suboptimal light conditions. This robustness against noisy signals is a hallmark of bistability^10,73–76^ that has been investigated in other developmental processes such as seed germination in plants^30,76,77^, and in the patterning of neuronal subtype specification at the vertebrate neural tube^73^. Hysteresis is another property linked to bistable systems, whereby a system’s response depends on its previous behaviour, conferring a “memory effect”^10,73^. It occurs, for instance, when a system remains committed to a certain state upon surpassing an activation threshold (i.e., the SAM retains commitment to flower) and only returns to the initial state (i.e., vegetative state) if the stimulus decreases below a different, lower deactivation threshold. Examples of reversion of floral transition^60,78,79^ and the inflorescence state commitment in a double shift SD-LD-SD experiment^48^ suggest the presence of hysteresis, further supporting the existence of a bistable mechanism. Noticeably, our modelled *soc1 ful* double mutant has an additional saddle-node bifurcation at relatively high age levels that makes the bistability region constrained to a small signal age range, facilitating the reversion to a vegetative state upon a decrease in the age signal (Figure 4G). This feature again connects this time-dependent bistability to the robustness of the transition. A quantitative analysis of the floral transition GRN under perturbations is yet required to demonstrate the hysteretic behaviour of the system.

Previous models of developing animal tissues have proposed that gene regulatory networks and the underlying dynamics can be modulated over the spatial dimension^71,73,74,80^. Our work provides *in silico* and *in vivo* evidence for the relevance of the spatial component in the system during the transition, showing that different meristematic regions can be subjected to distinct dynamic regimes. Detailed image analysis in the *rAP2* genotype, which maintains high AP2 levels while producing floral primordia, revealed persistently lower AP2 expression at the boundary regions of newly formed primordia (Figures S10A–S10C). This suggests a distinct, localised mode of AP2 action across SAM regions, explaining how floral primordia development can proceed despite high inhibitor levels at SAM central regions. Extending the model into the spatial dimension unveiled a rich variety of dynamical behaviours simultaneously occurring at different meristematic positions (Figures 6 and S21). Therefore, the spatial dimension offers an additional route to the dynamical system for being poised at criticality^20^, given that different regions of the meristem undergo different age signal trajectories. Future advances in live-imaging protocols will enable a more comprehensive assessment of the spatial dimension of the vegetative-to-inflorescence switch at the SAM.

While this study offers significant mechanistic insights into the transition, our analysis adopts a parsimonious approach and involves certain assumptions and simplifications. These factors would account for the fine-scale differences observed between experimental and model results. We grouped SOC1 and FUL into a single functional activator entity despite differences in their activation timing^41,43,56,81^. Similarly, to maintain analytical tractability, we omitted any potential secondary influence from *AP2-LIKE* genes^42,55,66^ and an explicit individual variable for miR172. Incorporating these factors or the implementation of a wave-like light signal to represent photoperiodic periodicity could increase the model’s accuracy. However, the primary goal of this study was not to perfectly fit experimental data, but rather to understand the fundamental nature of the floral transition dynamics and explain the range and likelihood of its possible behaviours. In the future, exploring how the time-dependent bistable switch is modulated in other environmental conditions and genetic perturbations would be of interest.

Our work highlights the importance of analysing time-dependent and transient behaviours in developmental contexts^5,64^, which can reveal relevant behaviours in plants and in other biological systems that are missed in traditional steady-state analyses^82^. A time-dependent bistable switch introduces a new timescale due to the emergence of a critical slowing down close to the bifurcation point. This critical behaviour confers dynamical flexibility and can prolong developmental transitions upon genetic and environmental perturbations. Furthermore, this slowing down drives the emergence of an intermediate, transient state, which might be associated with a new developmental state. Simultaneously, time-dependent bistability serves as a dynamical motif that makes the committed new developmental state robust against endogenous and external fluctuations. In the future, it will be important to develop new combined theoretical and experimental approaches to address how the underlying dynamics of developing tissues can be modulated simultaneously in both the temporal and spatial dimensions.

## Methods

### Model equations

We followed a parsimonious modelling strategy and reduced the complexity of the global floral transition GRN^31,35,40^ to a minimal ODE-based model that retains the core functional elements: a mutual inhibitory motif between floral activators and inhibitors modulated by a time-varying age signal. From a dynamical systems theory perspective, the reduction is based on the consideration that the qualitative behaviour of developmental switches is typically governed by a functional core of interactions. Biologically, we reduce the system to major flowering activators and inhibitors in the SAM, where the transition occurs, and importantly, our model is motivated by direct interactions, as AP2 binds to the promoters of *SOC1* and *FUL*, and in turn, SOC1 and FUL bind the promoter of *AP2*. By focusing on these key interactions, we ensure mathematical tractability, enabling the application of bifurcation theory to investigate, for instance, the existence of time-dependent bistable dynamical features, which is often difficult to resolve in high-dimensional discrete models. Thus, the models presented here are designed to provide a mechanistic framework to study the dynamics of floral transition at the SAM while remaining consistent with the available experimental data resolution.

The first two formulated models are non-spatial, single-compartment phenomenological models that describe the rate of change of protein (and RNA in model 3) concentrations of AP2 (*A*) and SOC1/FUL (*F*), using ordinary differential equations. The variable *S* represents the age-related developmental signal itself and serves as a proxy for the physiological age of the plant. Biologically, *S* models the abundance of SPL TFs (e.g., SPL15), and, to some extent, of MIR172. In Arabidopsis, the activation of these components is triggered by the gradual decline in MIR156 levels as plant ages, required for the floral transition. Hence, this microRNA and the SPL levels can be understood both as a timer and biological proxy of the age of the plant. Consequently, *S* accounts for SPL levels and is used as the external control parameter (*S*\*) for the bifurcation analyses to drive the system through its developmental states. This age signal is thus modelled through a simple basal production and a degradation proportional to its concentration. In all models, the rate of protein synthesis is assumed to be proportional to the corresponding activity of the promoter. Only pairwise interactions are considered and all proteins are assumed to be linearly degraded for simplicity. The following subsections describe the specific characteristics of each model. For the three models presented here, we performed a minimal qualitative parameter exploration searching for dynamical behaviours whose temporal dynamics were similar to those quantified in experiments. In the case of the model with AP2-autoinhibition, we required the system to display oscillatory behaviour (sustained oscillations) in the absence of age and activator variables. To achieve that the Hill exponent of AP2-autoinhibition must be greater than eight (see Model 2 for more details). All parameter values can be found in See Suppl. Tables 1 and 2. See Suppl. Tables 3 and 4 for more information about normalisations.

### Model 1: Toggle-switch model

In this case, AP2 and SOC1/FUL proteins mutually repress each other through Hill functions. They are additionally inhibited and activated by the age signal, *S*, respectively. The quasi-steady assumption for mRNA has been implemented (i.e., mRNA dynamics are faster than protein dynamics). The dynamic equations over time (*t*) for AP2 (*A)*, SOC1/FUL (*F*) and the age-signal (*S*) concentrations can be written as:

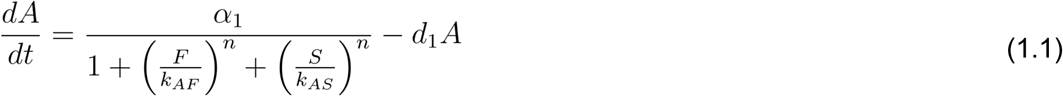

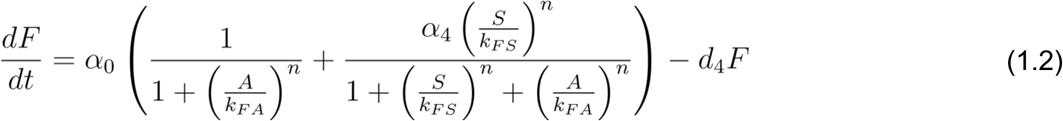

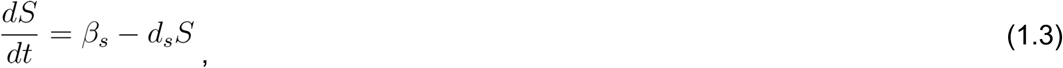

where *⍺*_1,_ *⍺*_0_ and *β*_s_ are the basal production rates of AP2, SOC1/FUL and age; *d_x_* represents the degradation rate constants for each variable *X*; *k_ZY_* are the concentrations of *Y* inhibiting or activating Z at which the production is half the saturated value; and exponent *n* sets the cooperativity (*n* = 2). The *F* production term – i.e., the first term within the parentheses – is formed by two parts to decouple the contributions of the photoperiodic and ageing pathways, whereby *⍺*_4_ is the relative strength between the ageing pathway with respect to the photoperiodic pathway. Unless otherwise stated, we consider *⍺*_4_<1, implying that the photoperiodic pathway has a stronger contribution. In the results shown in Figures S4C–S4D, we chose another parametrisation of Equation 1.2, namely,

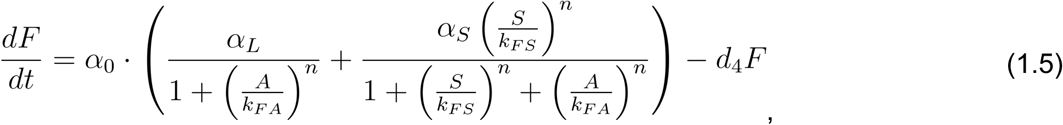

where *⍺*_L_ and *⍺*_S_ represent the strength of the photoperiodic pathway and ageing pathways, respectively. Nullclines in Figures 3E (right), S4A and S4B were obtained by setting equations 1.1 and 1.2 to zero (d*A*/dt = d*F*/dt= 0), and simultaneously treating S as a parameter (S*) as follows:

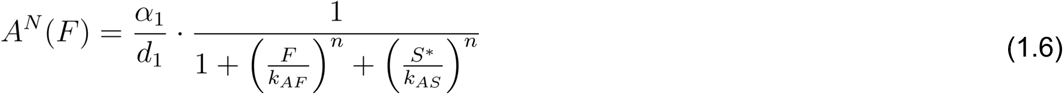

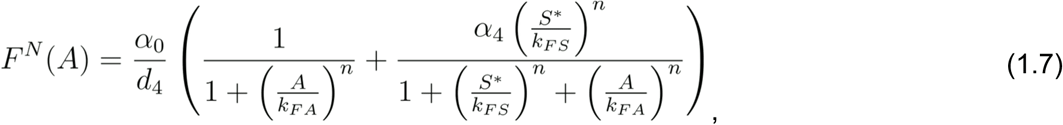

where *A^N^*(*F*) and *F^N^*(*A*) correspond to AP2 and SOC1/FUL nullclines, respectively. For Figure 3E, *A^N^* is plotted as a function of *A*, by first computing 1.7 and then introducing it into 1.6, leading to *A^N^*(*F*) = *A^N^*(*F^N^*(*A*)). A multiplicative factor is used for visualization purposes (*c =* 25).

Floral activator mutants were modelled through a reduced activation expression rate (*⍺*_0_), with late flowering phenotypes paired with lower *⍺*_0_ values (i.e., *⍺*_0_^wt^*>⍺*_0_*^ful^ >⍺*_0_*^soc^*^1^ *>⍺*_0_*^soc^*^1^ *^fu^*^l^), whereas for *rAP2*, the inhibition of AP2 by the age signal was completely removed, mimicking the resistance to miR172 binding, and therefore using the following equation:

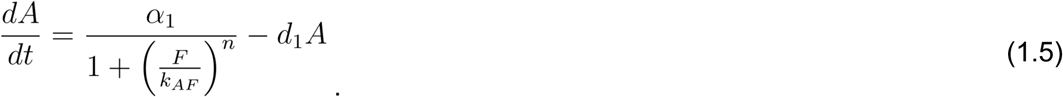

See Suppl. Table 1 for parameter values used in the simulations. For making the comparison between theory and experiments in the figures, note that the *A* variable is represented as [*AP2*], and the *F* variable is represented as [*SOC1*].

### Model 2: Toggle-switch model with a negative feedback loop

This model is based on Model 1, except that the quasi-steady assumption for AP2 dynamics is absent because a time delay is required to generate oscillations in the system, and this is achieved by considering RNA explicitly. The dynamic equations over time (*t*) for *AP2* mRNA (*Am)*, AP2 protein intermediate-I as cytosolic translated protein (*Ac*), AP2 protein intermediate-II as functional nuclear transcription factor (*Ap*), SOC1/FUL protein (*Fp*) and the age-signal (*S*) concentrations can be written as:

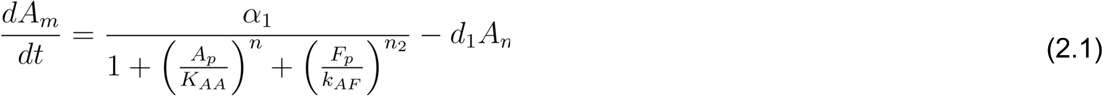

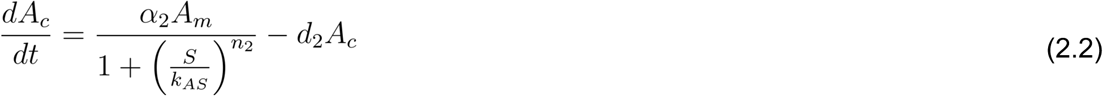

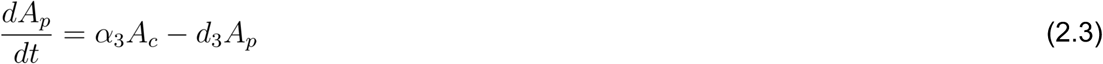

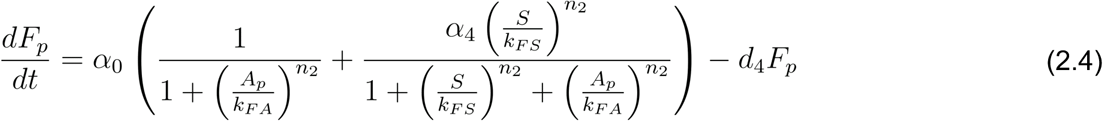

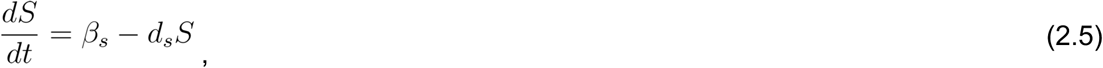

where *⍺*_i_ (i = 0,..,3) and *β*_s_ are the basal production rates of *Am, Ac, Ap*, *Fp* and age; *d_x_* represents the degradation constants; *k_ZY_* are the concentrations of *Y* inhibiting or activating *Z* at which the production half the saturated value (notice that *K_AA_* is the equilibrium constant associated to AP2 autoinhibition); and exponents *n* and *n_2_* set the cooperativity (*n* = 10 and *n_2_* = 2). AP2 is repressed by miR172, which is in turn activated by SPLs. This indirect inhibition is modelled here through the repression of *Ac* variable by the age signal, *S*. Although miR172 can also target AP2-RNA for degradation (cleavage mechanism^65,66^), most of its inhibition on AP2 occurs at the translational level^66^. This justifies why *S* was chosen to repress *Ac* and not *Am* in this model. Delay in AP2 self-repression was implemented, including two additional variables (*Am* and *Ac*). Without an explicit time delay, a negative feedback loop requires a minimum of three components to produce oscillations, provided there is sufficient nonlinearity^62,83^. To obtain limit cycle oscillations in such a simple three-component network, the Hill coefficient must be greater than eight^83^. See Suppl. Table 1 for parameter values. For *rAP2*, the inhibition of AP2 by the age signal was completely removed, mimicking the resistance to miR172 binding, and therefore using the following equation:

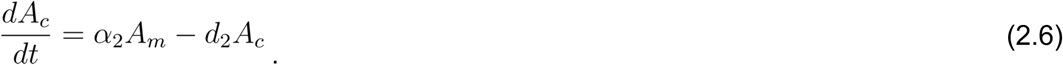

### Spatial model

This model consists of the 1D-spatial extension of Model 2. The dependence on the space variable has been introduced into the system through the age signal, which is now assumed to be able to diffuse in space.

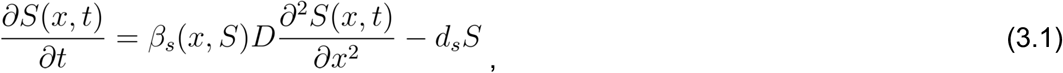

where the age signal production was considered just for a range of values of the spatial domain or when the age signal was above a certain threshold, specifically, *ꞵ_s_* > 0 if *x* ∊ [30,70] or if *S*(x,t) > 0. The diffusion of the age variable can be interpreted as a proxy of miR172 spatial movement through plasmodesmata during floral transition^37^. Zero flux boundary conditions were assumed, so the age variable did not diffuse out of the modelled space. The rest of the equations from Model 3 are identical to Eqns. 2.1-2.4 with the adjustment that if x > 70 µm, *⍺*_i_=1/(*⍺*_i_ᐧ(1+0.1ᐧ|*x*-70|)) for i =1 and i =3. See Suppl. Table 1 for parameter values.

### Model parametrisation

We adopted a parsimonious approach for the selection of model parameters, prioritising biological plausibility and the preservation of known regulatory motifs. In our Toggle-switch model with a negative feedback loop, for simplicity, we assumed identical basal expression rates (*⍺*_1_ = *⍺*_0_ = 1) of the activator and inhibitor. Protein stability was modeled by setting degradation rates below unity (*d*_1,2,3,4_ < 1), reflecting the increased half-life of proteins relative to their mRNA transcripts (*d*_1_^RNA^ > *d*_2,3,4_^Protein^)^84^. To distinguish the relative strength between the ageing versus the photoperiodic pathways, we set *⍺*_4_ < 1, to ensure that under LD conditions, the photoperiodic pathway has a stronger contribution to the progress of floral transition. For the equilibrium constants (*k_ZY_*), we assumed a lower threshold for the age signal to promote the activator than to repress the inhibitor (*k_FS_ < k_AS_*). This ensures the functional dominance of the activator variable (*Fp*) repressing the inhibitor of the floral transition. In the alternative scenario, low initial levels in the activator (*Fp*) increase the likelihood that AP2 is reduced prematurely by the age signal alone without any contribution from the SOC1/FUL variable. This hierarchy allows the reduction in AP2 to be a coordinated result of the contribution of both *S* and *F* variables: initially driven by the age signal (because at initial times low *Fp* levels are strongly repressed by high *Ap*) and subsequently reinforced by the increase in the activator, which ensures the final decline of the inhibitor, as observed experimentally. The equilibrium constants regulating the double negative feedback loop (*k_FA_* and *k_AF_*) and therefore, with a key role on the system’s dynamics, were parametrised under the same logic. Hence, a lower concentration of activator is required to repress the inhibitor (*k_AF_* = 0.2), compared with the opposite interaction (*k_FA_* = 1.5). The age signal parameters (*β*_s_ and *d*_s_) were calibrated to satisfy two global constraints. Specifically, *β*_s_ must be sufficiently high to avoid sustained oscillations in wt and single mutants (i.e., to reach the Hopf bifurcation without oscillatory behaviour in the temporal dynamics), yet the ratio between *β*_s_ and *d*_s_ must remain within a range that permits oscillatory behaviour in *soc1 ful* and *rAP2* genotypes. To obtain limit cycle oscillations in the AP2 autoinhibition three-component network, the Hill coefficient must be greater than eight (*n* = 8)^83^. All other Hill coefficients (*n_2_*) were set to 2 to maintain essential non-linearity without overparametrising the system in the absence of specific cooperative data.

Although the motivation of the analysis was to qualitatively understand the dynamics of the floral transition rather than to fit experimental data, the inhibitor dynamics was plotted alongside with the numerical simulation of the repressor variable to facilitate a simultaneous comparison of the data and model simulation results (Figures 2M, 4D and fourth column of 5C–G). To perform the conversion between the dimensionless model time (*t_num_*) and experimental time (*t_exp_*), we implemented a linear temporal scaling: *t_exp_*= *t_num_*ᐧ*γ*+offset. The scaling factor *γ* was determined by taking as reference the time point where AP2-VENUS first reaches its basal levels (17LDs; *γ*=17/125 for both models). The offset was fixed to align the simulation start (*t*=0) with the first experimental measurement at 7LDs (6.25 and 5.5 for Model 1 and 2, respectively). Consistency in normalisation was maintained across all datasets: AP2-VENUS data curves in wt, *ful*, *soc1*, and *soc1 ful* were normalised to the maximum median value reached in wt (which occurs at 7LDs), and AP2 model simulations (dashed lines) were similarly individually normalised to their maximum initial values. This value is identical for wt, *ful*, *soc1*, and *soc1 ful,* and, as in the experiments, also corresponds to the initial value (AP2 at *t*=0). This approach ensures that the normalised units represent a consistent and comparable relative concentration of the repressor across all genotypes. For *rAP2*, experimental data were individually normalised by its median value at 7LDs because the insertion of the transgene is in a different position, and in the model, normalisation was also performed separately with respect to its initial value.

As a verification step, we performed a systematic parameter exploration of the (*⍺*_0_,*⍺*_1_)-plane to identify optimal parameter sets to fit AP2-VENUS experimental data measured for each genotype (Figure S22). Root mean square error (RMSE) was computed on a 50 × 50 discrete logarithmic grid within the range [10^−3^, 10]). The parameter pairs with the minimum RMSE from the square grid and those derived from the Nelder-minimisation of the RMSE (via *minimize* function in Python) were compared with the Model 2 parameter selection used in the main text. We also investigated whether the quality of the fits could be improved by changing the temporal rescaling of the simulation data (*t_exp_*= *t_num_*ᐧ*γ*+offset).

This analysis confirmed that the parameters used to simulate the different genotypes are found among the optimal parameter sets (Figure S22). The RMSE associated to the wt parameter set was found to be located in the (*⍺*_0_,*⍺*_1_)-region with the minimum RMSE (Figures S22A–S22D). Optimal positions for the activator mutants were then computed by fixing *⍺*_1_ and allowing *⍺*_0_ to be optimised according to *⍺*_0_^wt^ > *⍺*_0_*^ful^* > *⍺*_0_*^soc^*^1^ > *⍺*_0_*^soc^*^1^*^ful^* (Figures S22B–S22E). For *rAP2*, an equivalent systematic parameter exploration of the (*d*_3_,*k_FS_*)-plane (*⍺*_0_ and *⍺*_1_ are identical to those in wt) confirmed an optimal selection of this parameter pair (Figure S22F). Although we identified alternative parameter configurations with lower RMSE, we considered these “mathematically optimal” sets not to be biologically plausible because they showed the lack of important dynamical features related to certain genotypes. For instance, for *soc1* and *soc1 ful* genotypes, these low-RMSE configurations failed to reproduce important dynamical features observed in experimental data such as the characteristic lengthening of the plateau in AP2 reduction (Figure S22B) or the smaller activator strength linked to the delayed flowering time in these mutants compared with wt and *ful* (Figure S22B). This is also reflected in the ratios of the steady-state activator levels (SOC1p^ss^) of each mutant versus wt (Figure S22D). The search for the best-fitting parameter configuration to recapitulate the studied genotypes is not a simple minimisation problem of the distances between model simulations and experimental results, if such minimisation erases key dynamical features. Instead, it is a search for parameter configurations able to satisfy the qualitative dynamics of the transition.

Alternative metrics such as Dynamic Time Warping (DTW) can better account for the shape of the curve that the model aims to recapitulate^85,86^, but involve a different temporal rescaling/stretching for each particular curve, which makes this metric unsuitable for a system where an absolute time is shared across all genotypes. Furthermore, although adding extra constraints or timepoint weights could lower the error and improve the optimisation, it carries a significant risk of overfitting. Thus, our selected parameters represent an optimal balance between mathematical fit and preservation of the essential dynamical features that define the floral transition. See Suppl. Tables 1–4 and Figure S22 for further information about normalisations and parameters.

### Bifurcation and stability diagrams

The analytical stability diagrams showing the stability regimes of the system (Figures 5A, S9A and S9B) have been computed through continuation algorithms of the saddle-node and Hopf bifurcations in Python using PyDSTool^87^. A bifurcation diagram for a given (*⍺*_0_,*⍺*_1_) as a function of *S\** is computed and then, from the saddle-node and/or Hopf bifurcation points, a continuation algorithm is run varying both parameters *⍺*_0_ and *⍺*_1_. Such continuation of the bifurcation point yields a 2D-stability diagram as shown in Figure S9A, which shows the different stability regions (monostable or bistable in this case) for the system for a particular *S*\*. In such a diagram, the boundaries between regions correspond to bifurcations. Because *S* is a system variable that depends on time, *S*\*-bifurcation diagrams can also be plotted as a function of time. This temporary dependency enables the study of the system’s stability at each time. Similarly, as S changes along the development, different 2D-stability diagrams are computed for different *S*\*-values (Figure S9A). All these diagrams can be collapsed into a single one that distinguishes among regions in the space with different stability regimes as *S\** changes. These three-parameter stability diagrams are shown in Figures 5A and S9B. The initial conditions of the numerical simulations are always the steady-state concentrations when *S* = 0.

### Numerical integration analysis

Numerical simulations were performed in Python using also PyDSTool^87^ toolkit with the VODE integrator, which can also be used for stiff systems. Although the output resembles that of a fixed time-step integrator, an adaptive time-step is used internally by this integrator. The creation of a Trajectory object is needed to later extract values at a desired resolution with the sample() method. Numerical simulations were initialised by setting the inhibitor variables (*A*, for Model 1; and *Am*, *Ac* and *Ap*, for Model 2) at its corresponding steady state for *S* = 0 (i.e., *ꞵ_s_*=0). If a limit cycle existed at *S* = 0, the center coordinate of the stable limit cycle of the corresponding inhibitor variable was used as the initial condition. Initial conditions for SOC1/FUL (*F*) and age (*S*) variables were set to zero for both models. Dynamics of *rAP2* in Model 2 were initialised with [*Am*, *Ac*, *Ap*, *Fp*, *S*] = [0, 2, 15, 0, 0]. Model 3 initial conditions were set to zero for *Am*, *Ac* and *Fp* and *Ap*(x,0) =15*e*(“#·%&!“’#), fitting the spatial profile of AP2-VENUS fluorescence protein along the SAM at 7LDs. For the age variable, *S*(0,x≥50) = -0.25x^2^ + 35x - 1125 and *S*(0,x≥70) = 100 to form an initial *S* profile as shown in Figure 6B. See Suppl. Tables 1 and 2 for parameters. To better understand how the bifurcation diagrams related to the actual numerical results, we visualised the bifurcation diagrams as a function of time, by assuming that *S\** is a variable *S(t)* governed by equation 1.3 (or 2.5) with initial conditions at 0, whose solution is *S*(*t*) = (*β*_)_/*d*_)_)01 − *e*^“*$+^ 2. The simultaneous plot of bifurcation diagrams and numerical simulations highlights a strong agreement between simulations and theory (Figures S3, S13D–S13G and S14C–S14E). This bifurcation over time provides a notion of instantaneous stability of the trajectory of the system, whose accuracy will depend on the relative timescales of the age signal progression with respect to the timescales of the activator and inhibitor variables. Henceforth, the computed steady states should be interpreted as instantaneous stability of the system at a certain time point.

The first and second derivatives in Figures S5E and S5F were also computed numerically on the normalised numerical simulations of AP2 dynamics that had been previously smoothed to avoid numerical errors in the derivative with a Gaussian 1D-filter (*gaussian_filter1d* in Python). The derivative was performed by second order central accurate differences in the interior points of the array and first order accurate one-side differences at the boundaries (*gradient* function). The second derivative was computed using a second time the previous function. For visualization purposes, the first derivative was normalised by itself. See code for more details.

Stochastic dynamics of the Toggle-switch model with autoinhibition were modelled by extending the deterministic Toggle-switch model (Model 1) with multiplicative noise, being the change in the modelled variables during the time d*t* described as d*X_t_* = f(*X_t_*)d*t* + σ*X_t_*d*W_t_*, where *X_t_* is the vector representing the studied variables, f(*X_t_*) is the deterministic vector field, σ is the noise intensity, and dW_t_ denotes a Wiener process^88^. Numerical integration was performed using the Euler-Maruyama method in Python with a constant time step (Δ*t* = 0.01). Two types of noise were implemented to account for biological variability: extrinsic noise, where a single Gaussian random number was applied globally to all variables at each time step; and intrinsic noise, where independent random Gaussian numbers were applied to each variable individually at each time step. We tested three noise intensities (σ = 0.005, 0.01, and 0.05). The generated random Gaussian numbers had mean 0 and standard deviation 1. Reflective boundary conditions were used to ensure all concentrations remained within the positive domain. For each genotype, *N* = 100 independent trajectories were generated. Median, first (Q_1_) and third quartile (Q_3_) were computed and compared against the deterministic simulation. To quantify the relative dispersion of trajectories over time, we also computed the Quartile Coefficient of Variation (QCV) at each timestep, QCV = (Q_3_-Q_1_)/(Q_3_+Q_1_).

### Plant material and growth conditions

All plants in this study were in the *Arabidopsis thaliana* Columbia-0 (Col-0) background. Mutant alleles were previously described: *ap2-12*^44^*, soc1-2*^89^ and *ful-2*^59^. The following transgenic lines were used: AP2::AP2-VENUS *ap2-12*^43^, AP2::AP2-VENUS *ap2-12 soc1-2*^43^ and AP2::rAP2-VENUS *ap2-12*^57^. AP2::AP2-VENUS *ap2-12 ful-2* and AP2::AP2-VENUS *ap2-12 soc1-2 ful-2* genotypes were generated in this study by crossing. An image dataset of AP2::AP2-VENUS *ap2-12*, and both AP2::AP2-VENUS *ap2-12 soc1-2* and *SOC1::SOC1-GFP soc1-2* images were obtained from a previous publication^43^. Plants were grown on soil under controlled conditions of LDs (16 h light/8 h dark).

### Confocal imaging

Shoot apices of plants at different developmental stages were dissected under a stereo microscope and fixed with 4% (w/v) paraformaldehyde (PFA; Electron Microscopy Sciences) by vacuum infiltration. Samples were incubated overnight in the dark and washed twice for 1 min in phosphate-buffered saline (PBS), to be transferred to ClearSee for 5 – 6 days^90^. Cell walls were stained with Renaissance 2200 [0.1% (v/v) in ClearSee]^91^ at least 1 day before imaging. Confocal microscopy was performed with a Stellaris 5 confocal microscope (Leica). Renaissance was excited at 405 nm, and image collection was performed at 445–475 nm (Figures 2A, 3A, 3B, 3E, 3F, S6J, and S10A). VENUS was excited at 514 nm, and the signal was detected at 520–600 nm. Images of SOC1-GFP were obtained from published data^43^.

### Fluorescence quantification

In this work, to analyse AP2 and SOC1 protein dynamics at the SAM during floral transition under LDs, we quantified previously published data^43^ (AP2::AP2-VENUS ap2-12 and *SOC1::SOC1-GFP soc1-2)* and added further experimental repeats and genotypes (AP2::AP2-VENUS *ap2-12,* AP2::AP2-VENUS *ap2-12 ful-2,* AP2::AP2-VENUS *ap2-12 soc1-2 ful-2,* and AP2::rAP2-VENUS *ap2-12).* Previously, at the tissue level, we quantified normalised levels of translational reporters, referred to as effective concentrations, exclusively in fixed-volume regions in the SAM^43^. Here, to increase the accuracy of the analysis at the tissue level and determine how SAM growth influences such measurements, we reanalysed the data using a new computational pipeline that accounts for meristem size by defining characteristic expression domains from each reporter. Furthermore, we performed quantification at the single-cell level for some of the data to corroborate that tissue and single-cell level quantifications were showing equivalent results.

### Tissue quantification

All confocal fluorescence z-stacks were processed and analysed using our custom-made MATLAB code^41,43^ (https://gitlab.com/slcu/teamHJ/pau/RegionsAnalysis), which we extended it from previous versions as explained below. The main objective of this analysis was to obtain reproducible measures of fluorescence intensity within the SAM. A normalised fluorescence intensity measure, together with spatial SAM-longitudinal-axis expression profiles, was computed as a proxy for the concentration of protein at the SAM. A semi-automated pipeline was developed and refined for this purpose (Figure S1). To achieve homogenous volumetric resolution, z-stacks were resized via bicubic interpolation, increasing the number of slices in the z-direction to account for differences in the xz-plane and z-direction.

To exclude fluorescence signals outside the region of interest and to quantify only the fluorescence intensity within the meristematic region, a pre-processing step was performed: a 3D paraboloid mask was automatically generated using the curvature of the meristem (Figure S1). First, a stack-slice interval containing the meristem tip was selected, and the cell wall signal within this interval was projected onto the orthogonal *xy* and *yz* planes (Figure S1B). Then, two curved lines following the parabolic outline of the SAM were drawn from the sum-of-slice projection of each plane. Later, a parabolic fitting was performed on these two drawn lines (Figure S1B), accounting for potential SAM tilting relative to the vertical axis.

Specifically, the code recursively fits parabolas in different orientations of the drawn outline and selects the one that minimises the *R*^2^ value. The coordinates of the apex, necessary to derive the 3D paraboloid equation, were computed from the two orthogonal parabolas fitted for each z-stack. The *zo* coordinate of the paraboloid was determined by averaging the apices of the orthogonal parabolas (Figure S1B). The parameter *a* in the parabola equation (i.e the curvature) was used to replace the denominator terms in the paraboloid equation (*c*1^2^ and *c*2^2^; Figure S1 legend) such that the paraboloid equation matched the linear and quadratic terms of their respective equations at *y* = *y*o and *x* = *x*o. Since z-stacks did not always encompass the entire meristem in the *yz* plane (lateral view), only the *xy* curvature was used (*c* ^2^ = *c* ^2^). Finally, to extract the fluorescence signal within the SAM using the 3D paraboloid, a 2D parabolic mask was created for each z-stack slice, and, for each slice, all intensity values of pixels outside the paraboloid were set to 0 (Figure S1C).

To exclude fluorescence signal at the boundaries of the SAM and primordia, the paraboloid’s curvature was increased relative to the original (Figure S1C), such that *a*’ = a/α, where *a*’ is the curvature of the new paraboloid and α is the image resolution (α < 1 μm) (Figure S1 legend). The 3D paraboloid was then divided into consecutive 3D subsections of 10 μm-height along the longitudinal axis of the SAM starting from the apex (Figure S1D). A fluorescence intensity concentration profile along the SAM longitudinal axis was extracted by computing the concentration of fluorescence signal within each section. Concentration was defined as the ratio of total intensity (sum of voxel intensities) to the total volume (sum of voxel volumes).

To determine the effective fluorescence concentrations in each meristem, a characteristic domain was defined based on the median concentration profile for each time point in each experiment (Figure S1E). The boundaries of these characteristic domains were defined by the points at which the normalised median concentration profile crossed a predefined threshold. Two or three thresholds were used to demonstrate the robustness of this quantification: 0.25, 0.50, and 0.75, always including the last two. The effective fluorescence concentrations per meristem were then calculated by dividing the total intensity within all sections included in the defined characteristic domain by the sum of the volume of all the voxels in these sections (Figure S1E). The discrete nature of the SAM-paraboloid partitioned sections was handled by shifting the boundaries of the characteristic domains to those of the nearest paraboloid subsection.

To quantify SOC1::SOC1:GFP *soc1-2 ap2-12* fluorescence, a Gaussian filter (*σ* = 2.5) was applied in the region within the paraboloid to homogenise a weak reporter signal. Two independent experiments for AP2::AP2:VENUS *ap2-12* were combined (Figures S2A–S2C) for all non-spatial AP2 data shown. The 7LDs data points from the *wt-3* experiment (Figures S2A–S2C) were not included in the analysis due to poor signal quality in the samples. An additional tissue quantification analysis was also performed for AP2::rAP2:VENUS *ap2-12* (Figures S10A–S10C). In Fiji, the fluorescence signal from 19, 21, and 24LDs samples was projected and summed within 10 z-stack slices (40 μm) containing the central region of the SAM, and then homogenised with a 3D-Gaussian filter (radius 2.5 μm). A curved line was drawn between the two uppermost primordia at opposite sides of the main shoot (Figure S10C). Normalised intensity profiles by time point were then extracted to highlight the lower expression of rAP2 at regions adjacent to the SAM primordia, compared with that in the central part of the SAM (Figure S10C).

All tissue quantification plots of AP2-VENUS included in this publication (i.e., effective concentrations in characteristic domains of expression and spatial concentration profiles along the SAM longitudinal axes) were normalised by the maximum median value reached in wt. Individual normalisation was performed for AP2:rAP2::VENUS fluorescence because the transgene is inserted in a different genomic position to that of AP2-VENUS, and the fluorescence intensity signal is therefore not directly comparable with AP2:AP2::VENUS of the other genotypes. SOC1-GFP tissue quantification was identically normalised (i.e., by the maximum median value). For further details, see Suppl. Table 3.

To compute the continuous median and IQR curves of the AP2-VENUS and SOC1-GFP data, fluorescence quantification median data points, as well as first and third quantiles, were interpolated through a Piecewise Cubic Hermite Interpolating Polynomial (PCHIP) in MATLAB (*pchip* function).

### Single-cell quantification

All confocal fluorescence z-stacks with AP2-VENUS or rAP2-VENUS reporters were also analysed at the single-cell level. Confocal fluorescence z-stack .lif files were converted to .tif files, and prior to segmentation, a 3D-Gaussian filter (r = 2 μm) was applied. Nuclei expressing the reporters were segmented using Stardist^92^ with StarDist Plant Nuclei 3D ResNet model^93,94^ (Figures S10D and S10E). In MATLAB, the segmented images were then re-labelled and single-nuclear features (nuclear concentration, volume and position) were computed (Figures S10F, S11, and S12) using the original and labelled images, together with the 3D-paraboloid generated for each SAM by the tissue quantification pipeline (Figure S1). In Figure 3I, single-cell nuclear quantification was normalised by the maximum median value reached, separately in *soc1 ful* and *rAP2*. Single-cell nuclear quantifications included in the Supplementary Information show raw concentration values without normalisation.

### SAM morphology quantification

To quantify the morphology of the meristem, its height and width were measured. The parabolic fits, obtained from the parabolic outline of the SAM drawn in the *xy* and *yz* planes between the uppermost primordia on opposite sides of the SAM (Figure S1), were used to obtain the coordinate of the apex and calculate the horizontal (width) and vertical (height) distance from the apex to the nearest primordium (Figures S2B, S2C, S6A, S6B, S6F, and S6G).

### Characterisation of critical slowing down in experimental data and simulated genotypes

To quantitatively assess the presence of the critical slowing down in the experimental data, we used two complementary analytical approaches. For wt and *soc1* genotypes, we first used a curve-fitting framework to compare the performance of three different functions on experimental data: a single exponential, a sum of two exponentials, and a sum of two arctangent functions (Figure S7, Suppl. Tables 5 and 6). Statistical comparisons of residuals and goodness-of-fit metrics between the experimental data and the fits determined that the sum-of-arctangents fit was optimal. This parametrisation enabled the identification of the slowing down timescale in the system, which is accentuated in floral activator mutants such as *soc1* (Figures S7E–S7G).

Secondly, we compared the critical slowing down dynamics within the theoretical model (Model 1) to the experimental data of AP2-VENUS fluorescence quantification (Figure S8). Specifically, we compared the velocity of AP2 reduction numerical simulations (Figure S8G) against the experimental velocities derived from both the numerical derivatives of interpolated AP2-VENUS curves (Figure S8H) and the discrete rates of change (Δ[AP2]/Δt) from normalised raw measurements (Figure S8I) across genotypes. We then extracted the different dynamical phases in AP2 decline from both theoretical and experimental datasets (Figures S8J–S8L). In particular, the transition times were partitioned into three characteristic times *τ*_0_, *τ*_g_, and *τ*_e_ defined as follows. *τ*_0_ (red region) extends from the start of the measurement (7LDs) until the derivative reaches its first local minimum (wt and *soc1*, Figure S8J). For *soc1 ful*, the *soc1* minimum was used as the temporal reference. *τ*_g_ (blue region) terminates when the derivative peaks again and reaches its maximum, marking the initiation of the final effective decline of AP2 (Figure S8J). *τ*_e_ (yellow region) expands until the termination of the floral transition, defined as the point of visible floral primordia (17, 24, and 26LDs for wt, *soc1*, and *soc1 ful*, respectively in Figures 3C and 3H). To compare the characteristic times among genotypes, characteristic times were normalised to the total sum of times for each genotype (Figure S8K). The same procedure was used to extract the characteristic times based on the discrete rate changes in AP2-VENUS concentrations (Figure S8L). The second double mutant measurement (*soc1 ful*\*) excludes the initial oscillatory transient (θ) to computate the characteristic times, with the reference initial time (*t*_ini_) taken as the first maximum after the oscillatory transient (15.44LDs and 14LDs for continuous and discrete derivatives, respectively).

Continuous derivatives (Figures S8H and S8J) were computed from the interpolated median fluorescence curves using second-order central accurate differences in the interior points of the array and first-order accurate one-side differences at the boundaries (*gradient* function in Python). The results were smoothed with a 1D Gaussian filter (*gaussian_filter1d*, n = 4).

### Statistics and reproducibility

Plants from a single experiment were grown together in the same chamber, and their positions in the growth chamber were randomised to avoid bias due to non-homogeneous growth conditions. To minimise potential effects of the clearing treatment on fluorescence signal quantification, samples of each genotype were included in each imaging session when time point acquisition extended beyond one day. Meristems exhibiting severe defects (i.e. tissue breakage, poor cell wall dye penetration, or blocking of the fluorescence signal by leaf primordia) and/or strong developmental differences compared to the other samples from the same time point were not considered for fluorescence quantification. A detailed list of the meristems analysed in each experiment is provided in the code files. Investigators were not blinded in allocation during experiments and outcome assessment, because the material was rigorously labelled, thus making blinding impossible. See Suppl. Tables 7-13 for statistics.

## Data availability

The original confocal microscopy images for one experiment of AP2-VENUS expression in *ap2* and *ap2 soc1* and SOC1-GFP in *soc1* and in *soc1 ap2* have been obtained from this study^43^. The new acquired confocal microscopy images are available on Edmond database (https://doi.org/10.17617/3.X6UHYT). Source data used for the analysis is available in the OSF repository https://osf.io/jyq58/.

## Code availability

Source code for the the image analysis at both tissue and single cell levels can be found in the OSF repository https://osf.io/jyq58/ and also in the following GitLab repositories https://gitlab.gwdg.de/devplantpatterning/Publications/2025-rodriguez-maroto_wang_et_al and https://gitlab.com/slcu/teamHJ/pau/RegionsAnalysis/-/tags/Rodriguez_Maroto_et_al_2025_Region_Analysis_code.

## Acknowledgments

We thank Josep Mercadal, John Chandler, and Ruben Perez-Carrasco for critical comments on the manuscript, and Tobias Bollenbach, Marta Ibañes Miguez, and James Locke for fruitful discussions. G.R.-M. acknowledges funding from the International Max Planck Research School on “Understanding Complex Plant Traits Using Computational and Evolutionary Approaches”, and K.W. received a studentship from the China Scholarship Council. P.C. and M.C. received a post-doctoral fellowship from the Alexander von Humboldt foundation (ESP - 1238682 - HFST-P and ITA 1216206 HFST-P, respectively), P.C. also received a post-doctoral Marie Curie fellowship (101150752 from HORIZON-MSCA-2023-PF-01). G.C. received funding from the Deutsche Forschungsgemeinschaft (DFG) through grant CO 318/14-1. G.C. and P.F.-J. also received funding from the DFG through the Cluster of Excellence CEPLAS (EXC 2048/1 Project ID: 390686111) and from a Core Grant from the Max Planck Society. P.F.-J. also acknowledges funding from the National Science Foundation (US NSF, https://www.nsf.gov), Division of Biological Infrastructure DBI-232051.

## Author contributions

G.R.-M., K.W., G.C., and P.F.-J. conceived and designed the study. K.W. generated the AP2::AP2-VENUS *ap2-12 ful-2* and AP2::AP2-VENUS *ap2-12 soc1-2 ful-2* transgenic lines. G.R.-M., K.W. and M.C. performed confocal imaging. G.R.-M., P.F.-J. and P.C.-F. designed the imaging quantification pipelines. G.R.-M. built and performed the quantification analyses. P.C.-F. assessed primordia identity. G.R.-M. and P.F.-J. formulated and analysed the mathematical model. G.R.-M., G.C. and P.F.-J. wrote the manuscript. P.F.-J., G.C., M.C. and P.C.-F. provided scientific guidance throughout the project. All the authors discussed the results and commented on the manuscript.

