## Supplemental information for "Time-dependent bistability leads to critical slowing down during floral transition in Arabidopsis"

#### Supplementary Figures

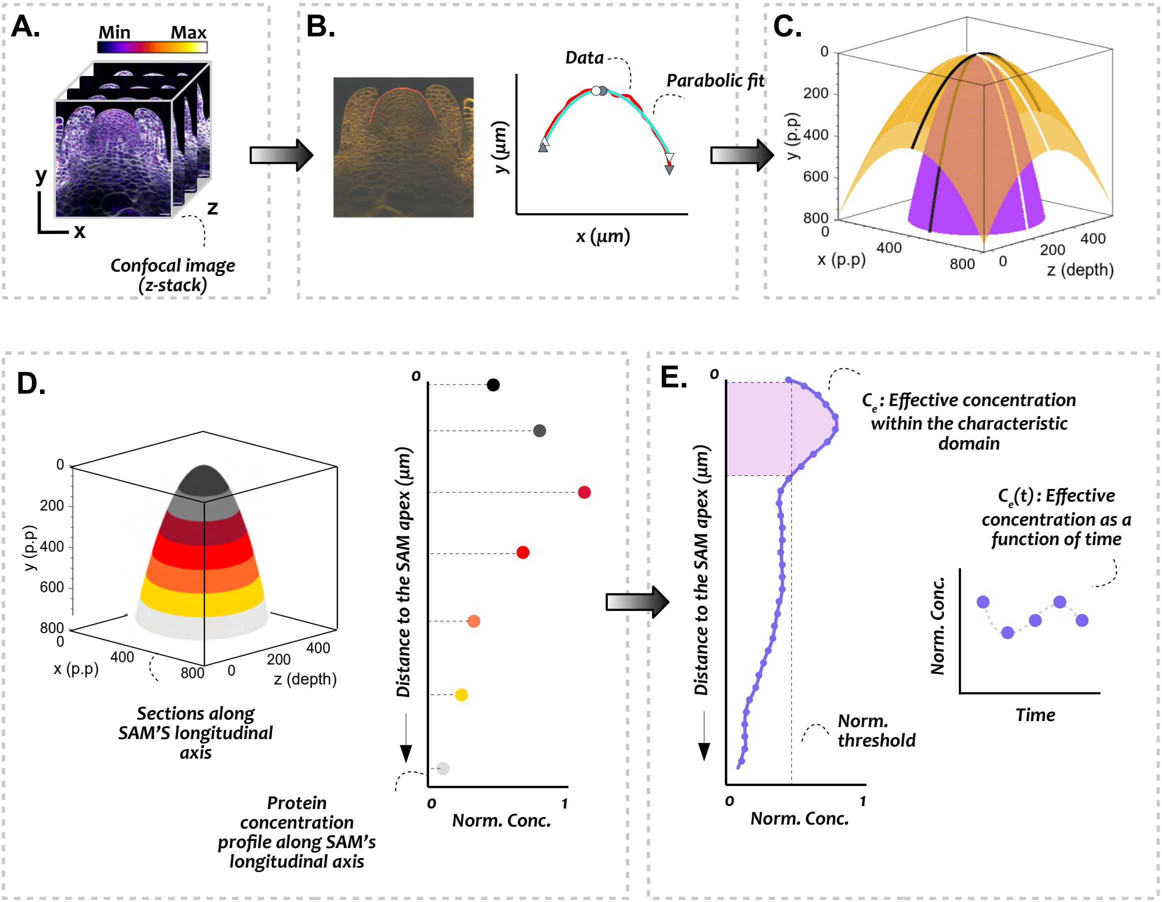

**Figure S1. Tissue quantification pipeline for temporal and spatial protein dynamics of AP2-VENUS and SOC1-GFP during floral transition at the SAM.** A) Fluorescence confocal microscopy images were obtained from the lateral side. The panel shows z-stacks of AP2::AP2:VENUS meristematic expression in *ap2-12* of a plant grown for 12LDs. The membrane marker channel is displayed in white, and the AP2-VENUS fluorescence signal is coloured according to the Fire colour look-up table in Fiji. Scale bar = 20 µm. Acquisition parameters are described in Materials and Methods. B) A red curved line marks the meristem outline to the boundaries of the primordium closest to the SAM tip, on a sum-projection of a specific z-stack interval. The parabolic fit of the red curve is shown in light green. Triangles mark the beginning and end points of the drawn curve line (white) and fitted parabola (black). Circles indicate the apex position for both the drawn curve line (white) and fitted parabola (black). C) 3D paraboloids constructed from previously extracted orthogonal parabolas. In orange, a paraboloid using the original curvature values, with extracted orthogonal parabolas superimposed on top. In purple, a paraboloid with higher curvature focuses on the central area of the meristem along its longitudinal axis. The purple paraboloid was used to generate a mask. Intensity values of pixels whose positions lie outside the paraboloid were set to zero (p.p. = pixel position). D) The 3D-paraboloid was divided into 10 µm height sections along the SAM longitudinal axis. Fluorescence signal concentration was computed in each section as the ratio of total intensity (sum of pixel intensity) to the total voxel volume. Each 3D-paraboloid section thus yields a concentration value that can be plotted as a profile of concentration versus distance to the SAM apex and normalised by the maximum within that profile. E) To account for SAM growth during floral transition, effective concentration measurements were computed within characteristic domains. The beginning and end points of these domains are defined by the coordinates where the normalised concentration profile crosses a given threshold (0.25, 0.50 or 0.75). Finally, the sum of pixel intensity from all 3D-paraboloid sections within the characteristic domain was divided by the total volume, resulting in an effective concentration. Characteristic domains were computed along the normalised median signal concentration profile for each time point.

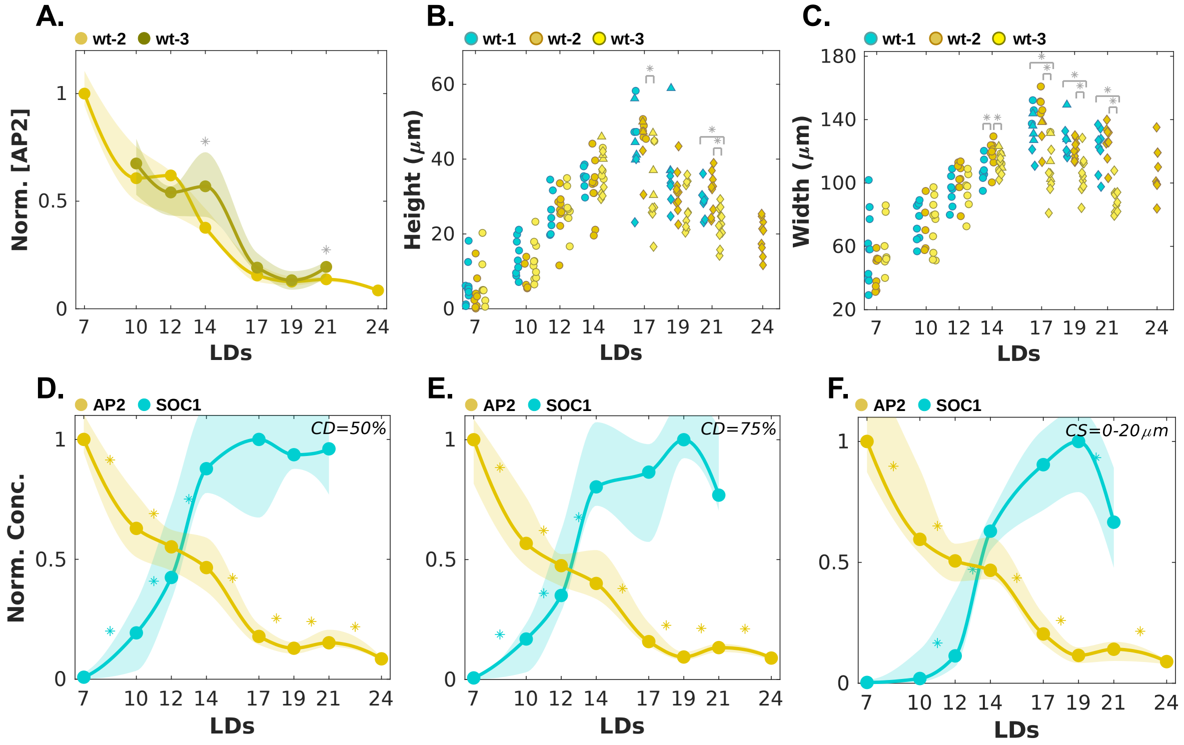

**Figure S2. Tissue fluorescence signal of SOC1 and AP2 and morphological quantification analyses of the SAM during floral transition.** A) Normalised wt AP2 concentration computed in defined characteristic domains as a function of time for two different experiments. Filled monochrome dots represent normalised median protein concentration measurements, and the shaded regions represent the inter-quartile range (IQR). Normalisation was performed with respect to the maximum median value reached in the wt-2 experiment. Asterisks indicate significant differences between the two experiments at each time ($\alpha$ < 0.05 Mann-Whitney-Wilcoxon test). B and C) Quantification of SAM height (B) and width (C) from the wt-SAMs whose reporter expression was quantified in Figures 2C and 2D. Data in cyan are from the SOC1-GFP experiment, and in yellow/brown from the two different AP2-Venus experiments. The quantification pipeline described in Figure S1 also provides these two SAM morphological observables from the drawn parabolas. SAM height is defined as the vertical distance from the apex to the adaxial side of the youngest primordium, and SAM width is the horizontal distance between the youngest primordia. Dots, triangles, and diamonds indicate SAMs with either only leaf primordia (vegetative state), cauline leaves, and floral primordia (inflorescence SAM), respectively. Statistics as in (A). D–F) Examples of AP2 and SOC1 concentrations computed in a defined characteristic domain (CD) when using a CD=50% (D) and CD=75% (E) threshold (Methods) and within the first top 20 μm of the SAM (F) as a function of time; CS constant size region. Characteristic domains are defined on the median normalised profile at each time point by taking its beginning and end points as the spatial points along SAM longitudinal axes at which the normalised concentration profile intersects 0.5 and 0.75 thresholds, respectively. Asterisks indicate significant differences between consecutive time points for each protein (Mann-Whitney-Wilcoxon test, $\alpha$ < 0.05).

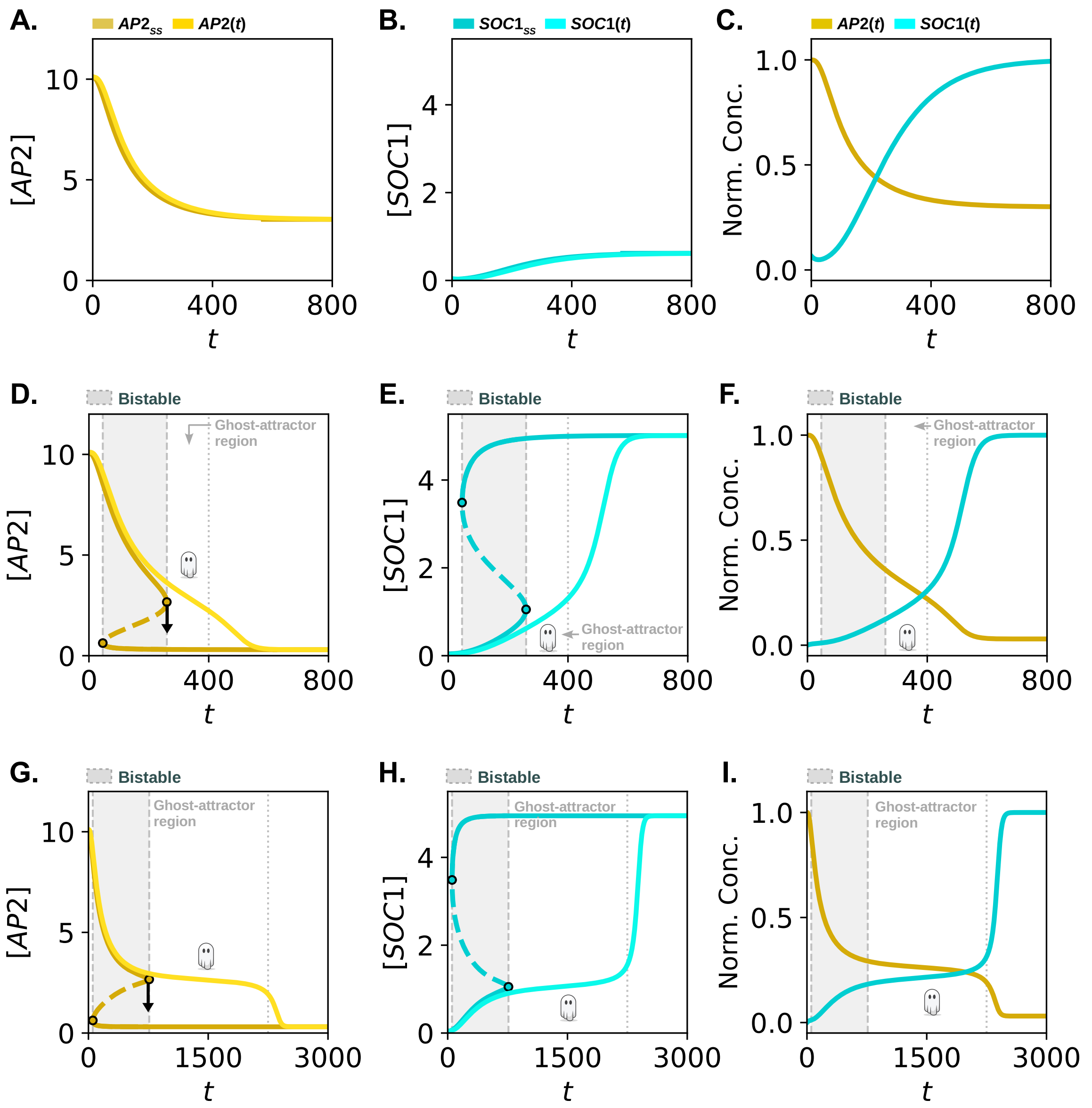

**Figure S3. Model comparison between monostable and bistable transition scenarios.** A) AP2 model numerical simulation (yellow) and instantaneous stable state $AP2_{ss}$ (dark yellow) concentration as a function of time in a monostable scenario (Methods). B) SOC1 model numerical simulation (cyan) and instantaneous $SOC1_{ss}$ (dark blue) concentration as a function of time in a monostable scenario. C) Normalised model AP2-SOC1 temporal dynamics in a monostable scenario. D–F) Equivalent figures to A–C illustrating the model's dynamics in a bistable scenario. The grey region indicates the bistable region, and a "ghost" is used to highlight the time window in which the ghost attractor exerts the strongest influence on the system dynamics. G–I) Equivalent figures to D–F in a bistable scenario with a very long plateau. Coloured-filled dots indicate the saddle-node bifurcation points. See Suppl. Tables 1 and 2 for parameter values.

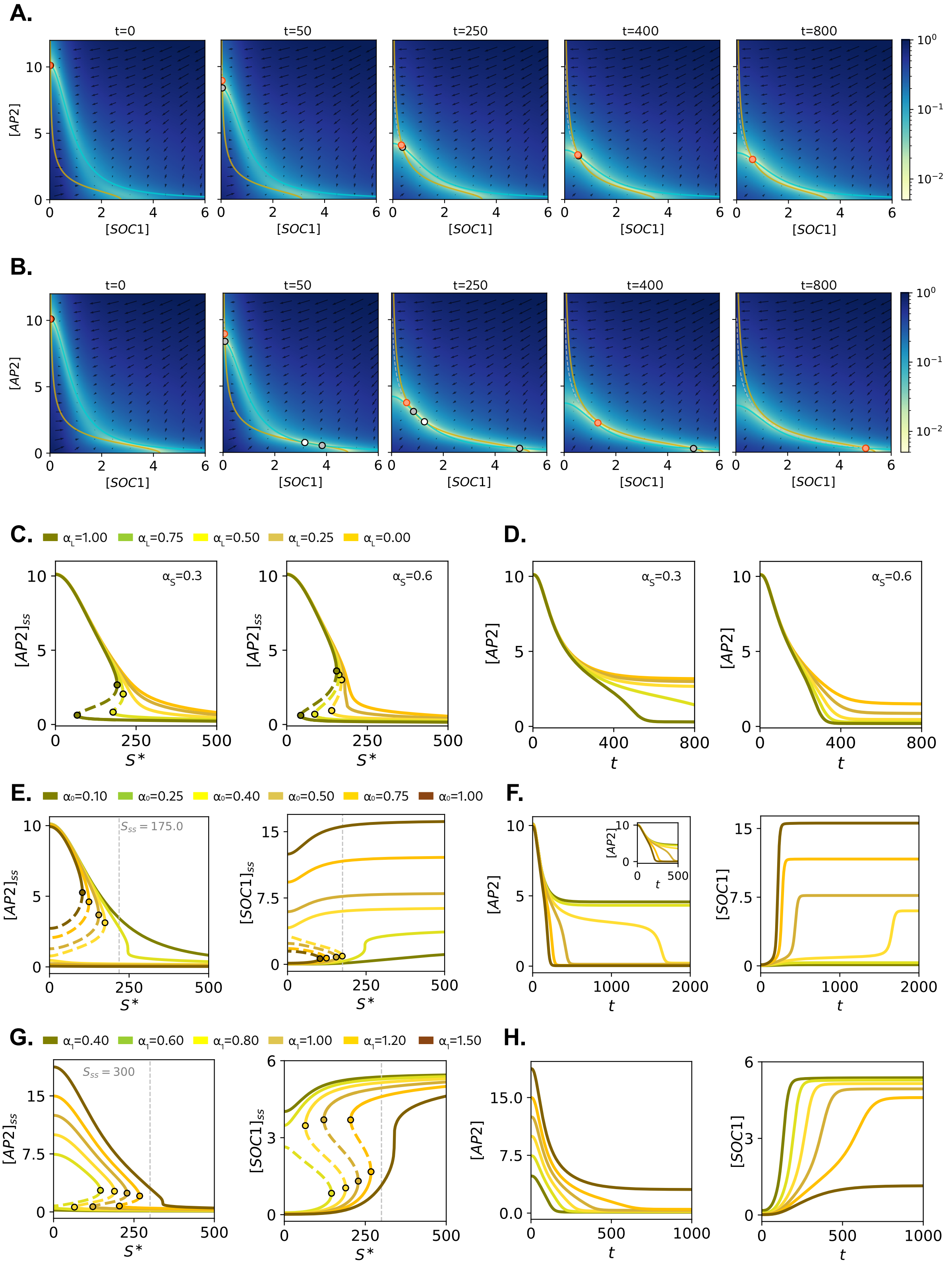

**Figure S4. Understanding the effects of bistability in the system dynamics.** A and B) AP2–SOC1 model phase portrait diagrams showing trajectory evolution over time for a monostable (A) and a bistable (B) scenario. The colour of the phase portrait represents the speed of the system at each time point (Methods). See colour bar. Continuous yellow and blue lines represent SOC1 and AP2 nullclines, respectively. The orange dot represents the current position of the system. Stable and unstable fixed points are illustrated as grey and white-filled dots, respectively. The initial conditions of the system were set to zero for SOC1 and the age variable, and to its steady-state concentration for $\beta_s = 0$ for AP2. C and D) AP2 plots (C) and corresponding numerical simulations (D) for different $\alpha_L$ and $\alpha_S$ values (see Methods). E and F) AP2 and SOC1 bifurcation plots (E) and corresponding numerical simulations (F) for different values of SOC1/FUL production rates ($\alpha_0$). G and H) Same as (G) and (H) for different values of AP2 production rates ($\alpha_1$). Coloured dots represent saddle-node bifurcation points. Vertical dashed lines in (E) and (G) indicate the $S_{ss}$ value reached in all numerical simulations. See Suppl. Tables 1 and 2 for parameter values.

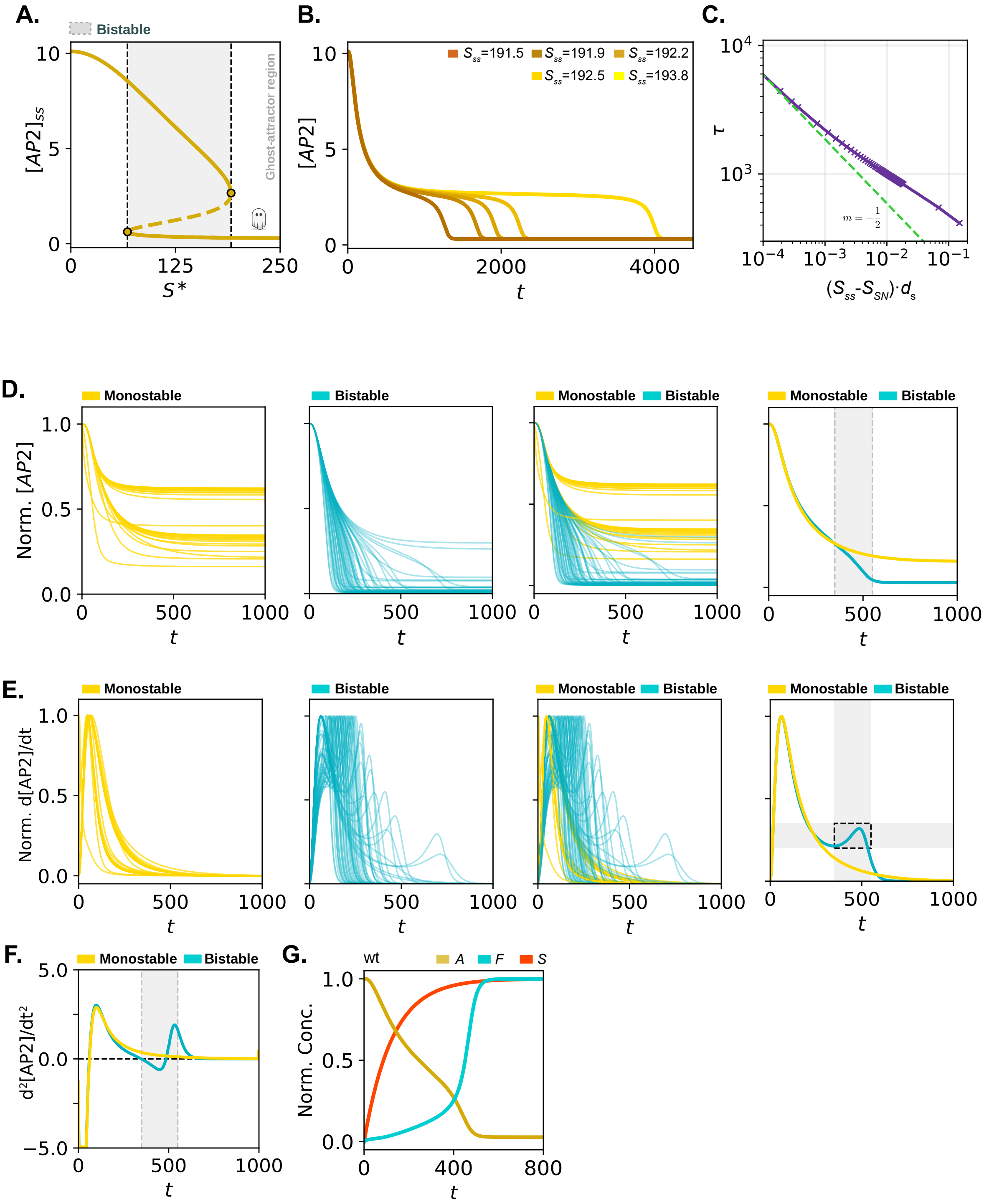

**Figure S5. Influence of the ghost attractor in the system through the critical slowing down.** A) AP2 bifurcation diagram for a bistable scenario. The grey background indicates the region of bistability, and the ‘ghost’ after the saddle-node bifurcation indicates the presence of a ghost attractor. Coloured dots indicate saddle-node bifurcation points. Solid and dashed lines represent stable and unstable states, respectively. B) AP2 numerical simulations show progressively longer plateaus for different $S_{ss}$ values. C) Simulations showing time $\tau$ that AP2 takes to decay with respect to the distance to the saddle-node bifurcation $\Delta S \bullet d_s = (S_{ss} - S_{SN}) \bullet d_s$. $\tau$ is computed as the time that AP2 requires to reach the lower branch of the bifurcation diagram in (A). For smaller $\Delta S \bullet d_s$, the computed $\tau$ is dominated by the slowing-down phase and follows a power law with an exponent of -½, in agreement with bifurcation theory. The dashed green line represents the theoretical power law relation $\tau \propto (\Delta S \bullet d_s)^m$ with m=-½. D–F) Numerical simulations of AP2 dynamics (D), its first derivative (E), and its second derivative (F) in monostable (yellow) and bistable (blue) scenarios. In the rightmost graphs in D and E, a representative individual scenario was selected to highlight the differences. G) Numerical simulations showing temporal dynamics of AP2 (cyan), SOC1 (yellow), and S (red) variables in wt. In (D) and (E), each curve is individually normalised by its maximum (D) and by its maximum absolute value (E). In (G), each variable is normalised by its maximum value reached in the simulation. See Suppl. Tables 1–3 for parameter values and normalisations.

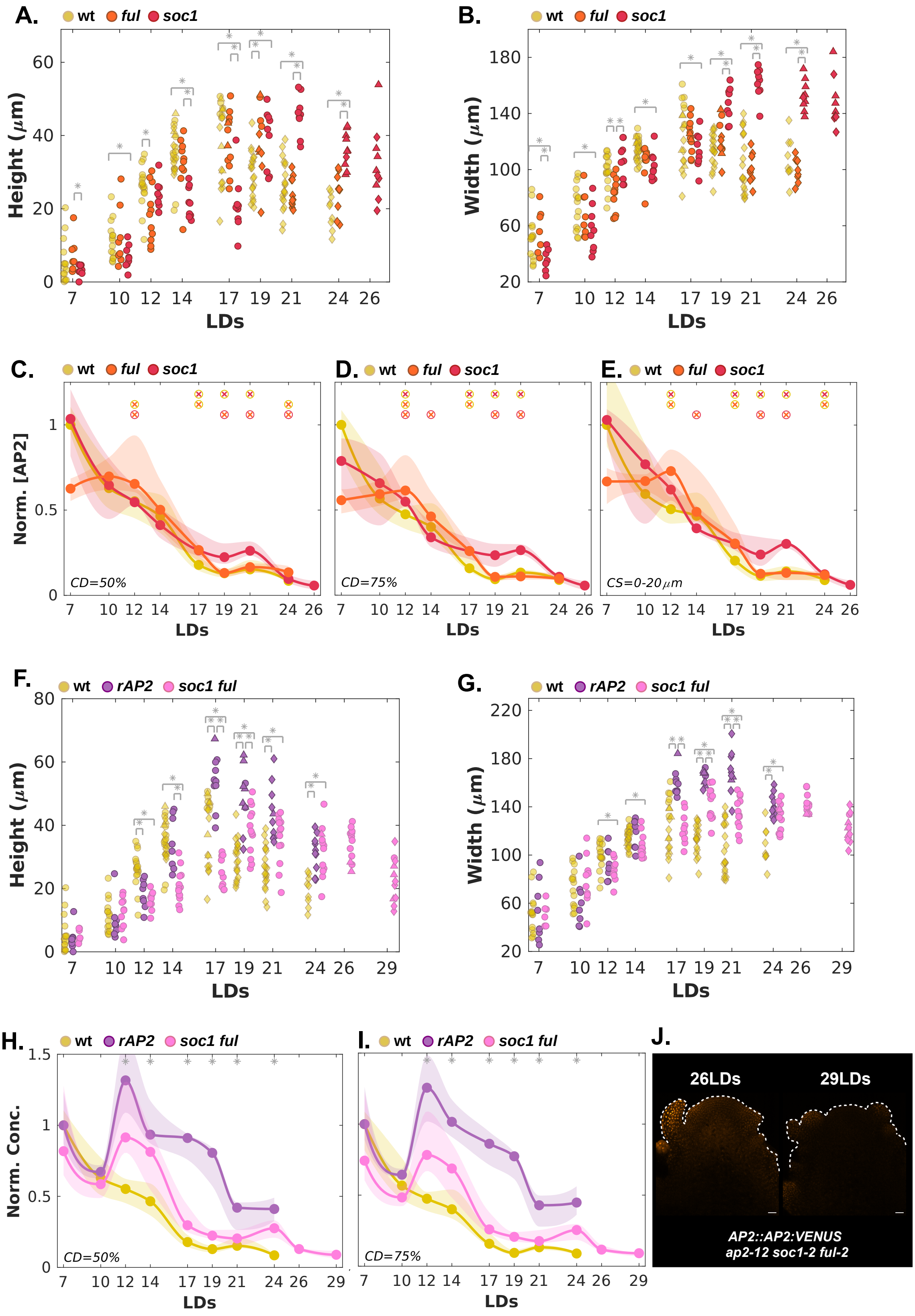

**Figure S6. Morphological observables and temporal protein dynamics of AP2-VENUS in wt, *ful*, *soc1*, *soc1 ful*, and rAP2-VENUS during floral transition at the SAM.** A and B) Quantification of SAM height (A) and width (B) of wt, *ful,* and *soc1* SAMs whose reporter expression was quantified in Figure 3C. C–E) Examples of AP2 normalised concentrations computed in wt, *ful,* and *soc1* in characteristic domains (C and D) and within the first top 20 µm of the SAM (E) at different time points. The colour of the asterisk and its surrounding circle indicates significant differences between the genotypes with the corresponding colour (Mann-Whitney-Wilcoxon test, $\alpha$ < 0.05). F and G) Quantification of SAM height (F) and width (G) in wt, *soc1 ful* and rAP2 plants whose reporter expression was quantified in Figure 3F and 3G. H and I) AP2 normalised concentrations computed in wt, *soc1 ful* and rAP2 in two characteristic domains at different time points (threshold values = 0.5 and 0.75 in (F) and (G), respectively). Asterisks indicate significant differences between wt and *soc1 ful* (Mann-Whitney-Wilcoxon test, $\alpha$ < 0.05). J) Expression pattern of AP2:AP2::VENUS in *ap2*-12 *soc1 ful* at the SAM at 26 and 29LDs. Scale bars = 20 µm. In A, B, F, and G, circles, triangles, and diamonds indicate SAMs with either only leaf primordia, cauline leaves, or floral primordia, respectively. Asterisks mark significant differences among genotypes at each time (Mann-Whitney-Wilcoxon test, $\alpha$ < 0.05). See Suppl. Tables 7–13 for *p*-values. Normalisation of fluorescence concentration measurements was performed with respect to the maximum median value in wt (which is also the initial wt concentration at 7LDs). For rAP2, because the position of the transgene insertion differs from that of AP2-VENUS, fluorescence measurements were normalised with respect to its initial concentration (at 7LDs).

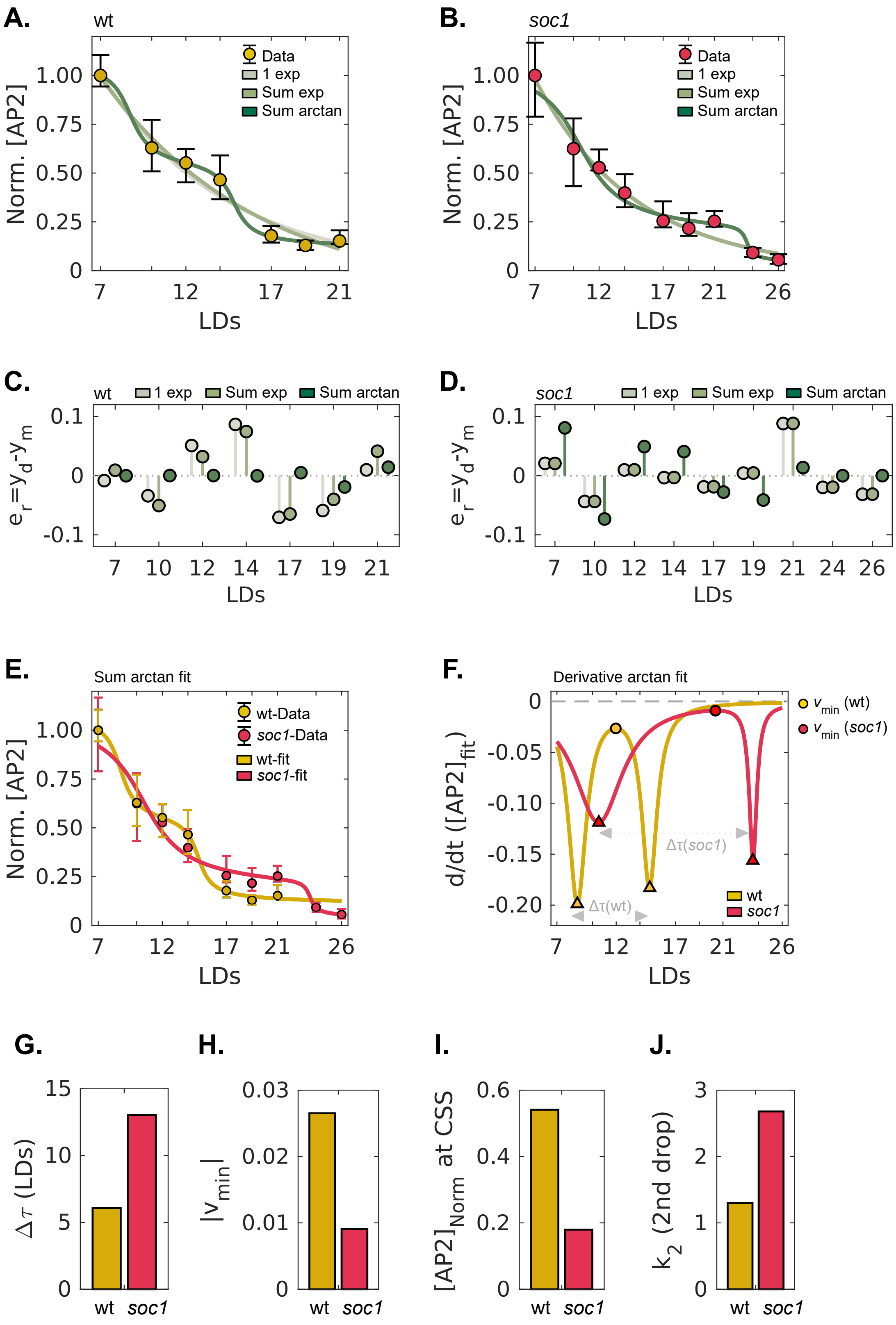

**Figure S7. Mathematical fits of AP2-VENUS dynamics quantify critical slowing down in wt and its accentuation in *soc1* mutant.** A and B) Comparison of a single exponential, a sum of two exponentials, and a sum of two arctangent functions fitted to AP2-VENUS fluorescence quantification data in wt (A) and *soc1* (B). Coloured dots and error bars indicate the median and IQR. Data are normalised to the maximum median wt value (i.e., values at 7LDs). C and D) Residual analysis ($e_r = y_d - y_m$) for the three models shown in A and B. The sum of the arctangent function minimises the error at the transition points. E) Direct comparison of the sum-of-arctangent fits for wt and *soc1*. F) Analytical derivatives (*d*/*dt*) of the sum-of-arctangent fits. The local decline velocity minimum (circles) identifies the "bottleneck" of the ghost attractor. We define the duration of the critical slowing down ($\Delta\tau$) as the temporal distance between the two points of maximum decline (triangles). G–J) Characterisation of dynamical features derived from the arctangent fits in wt and *soc1*. Bar plots show the bottleneck duration, $\Delta\tau$ (G); the minimum absolute speed of decline, $|v_{min}|$ (H); the normalised concentration of AP2-VENUS at the transition plateau (I); and the slope of the second decline, $k_2$, after the critical slowing down. These metrics confirm that whereas wt exhibits the effect of the ghost attractor (bottleneck), this is significantly accentuated in *soc1* dynamics by increasing $\Delta\tau$ and decreasing $|v_{min}|$.

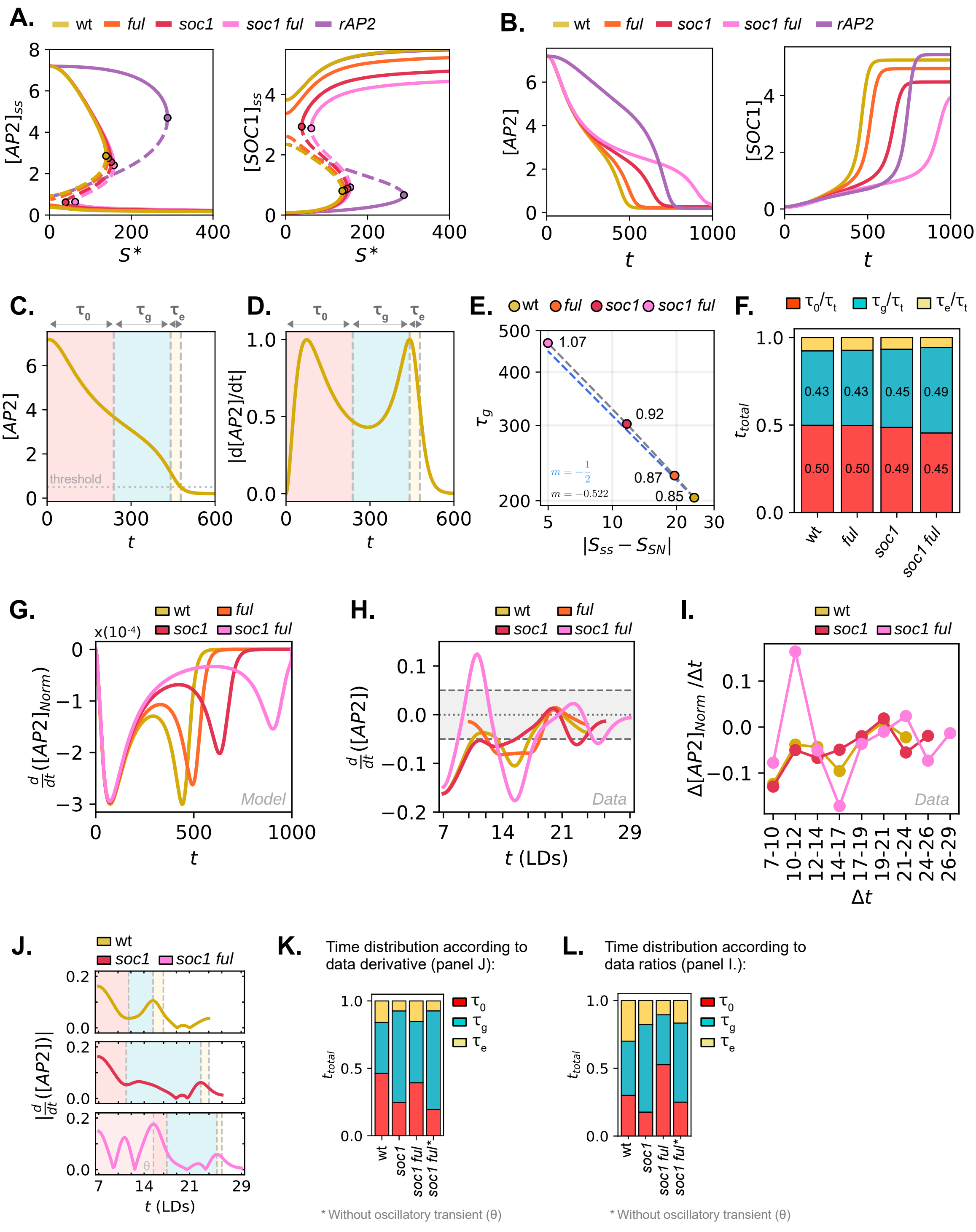

**Figure S8. Model and data analyses of the critical slowing down dynamics.** A) AP2 (left) and SOC1 (right) bifurcation diagrams for wt, *ful*, *soc1*, *soc1 ful,* and *rAP2* genotypes. Coloured dots indicate saddle-node bifurcation points. Solid and dashed lines represent stable and unstable states, respectively. B) Numerical simulations of AP2 (left) and SOC1 (right) temporal dynamics across genotypes. C and D) Example of time distribution computation for wt. The red region extends until the system reaches the age-value associated with the saddle-node bifurcation in the upper branch ($\tau_0 = t(S_{sim}{=}S_{SN})$) in the numerical simulation. The blue region, representing the time the system is influenced by the ghost ($\tau_g$), terminates when the normalised derivative of the AP2 variable (see D) in the numerical simulation peaks for the second time. The yellow region ($\tau_e$) expands until the AP2 variable reaches a basal threshold (th. = 0.5). E) Time under the influence of the ghost ($\tau_g$) against the distance from *S* steady state value ($S_{ss}$) and *S*-coordinate of the saddle-node bifurcation ($S_{SN}$) for wt, *ful*, *soc1,* and *soc1 ful* numerically simulated genotypes. The blue-dashed line is the theoretical power-law with exponent $m$=-½. F) Distribution of times ($\tau_0$, $\tau_g$, and $\tau_e$) for wt, *ful*, *soc1,* and *soc1 ful* genotypes. Normalisation was performed with respect to the total sum of times for each genotype. G-I) Velocity of AP2 reduction across genotypes as (G) derivatives of the normalised AP2 model variable as a function of time; (H) derivatives of the normalised AP2-VENUS experimental interpolated median data curves; and discrete experimental rate changes of normalised AP2-VENUS fluorescence ($\Delta[AP2]_{Norm}/\Delta t$). J) Allocation of experimental time phases ($\tau_0$, $\tau_g$, and $\tau_e$) as in C mapped onto the absolute derivative of normalised AP2-VENUS median curve data, $|d[AP2]/dt|$. Data are the same as in H. K) Distribution of times ($\tau_0$, $\tau_g$, and $\tau_e$) for wt, *soc1,* and *soc1 ful* genotypes associated with J. L) Time distribution allocation based on discrete rate changes from I. Same procedure as in K. In K and L, the second double mutant measurement (*soc1 ful**) excludes the oscillatory transient (θ) observed in this double mutant from the computation of the times. See Methods for further details.

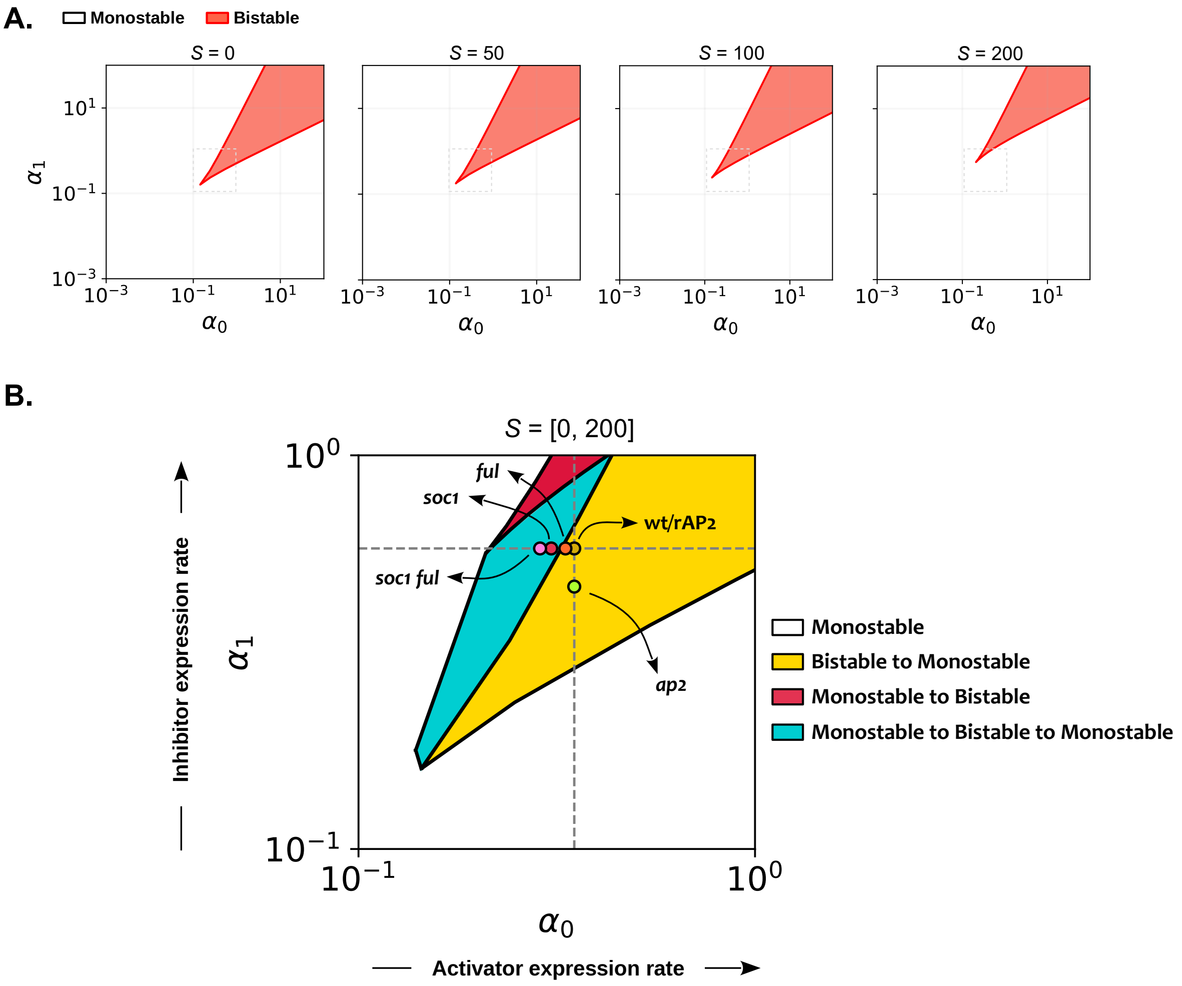

**Figure S9. Stability analysis across genotypes.** A) ($\alpha_0$,$\alpha_1$) 2D-stability diagram for different *S* values. For each ($\alpha_0$,$\alpha_1$) pair, steady states were computed. If only one solution (indicating monostability) was found, that coordinate is depicted in white. If three solutions (one unstable and two stable, indicating bistability) were found, the point is shown in red. The red region thus encompasses all parameter pairs that lead to bistability in the system. B) ($\alpha_0$,$\alpha_1$) 3D-stability diagram. This diagram summarises the evolution of the ($\alpha_0$,$\alpha_1$) 2D-stability diagram from (A) as *S* changes. Different regions are distinguished based on the stability regimes each parameter pair ($\alpha_0$,$\alpha_1$) can generate. The range of ($\alpha_0$,$\alpha_1$) values displayed corresponds to the grey-dashed square shown in (A). White: ($\alpha_0$, $\alpha_1$) pairs that lead to a unique solution (monostable) when $S \in [0,200]$. Yellow: ($\alpha_0$,$\alpha_1$) pairs for which the system always shows bistability when $S \in [0,200]$. Red: ($\alpha_0$, $\alpha_1$) pairs for which the system is initially monostable, but eventually becomes bistable as *S* increases. Blue: ($\alpha_0$,$\alpha_1$) pairs for which the system is initially monostable, then shows bistability for a certain range of *S* values, and later returns to a monostable regime. Coloured circles indicate the position of wt, *ful*, *soc1*, *soc1 ful*, *rAP2*, and *ap2* genotypes. Dashed grey lines indicate ($\alpha_0$,$\alpha_1$) used for wt. See Suppl. Tables 1 and 2 for parameter values.

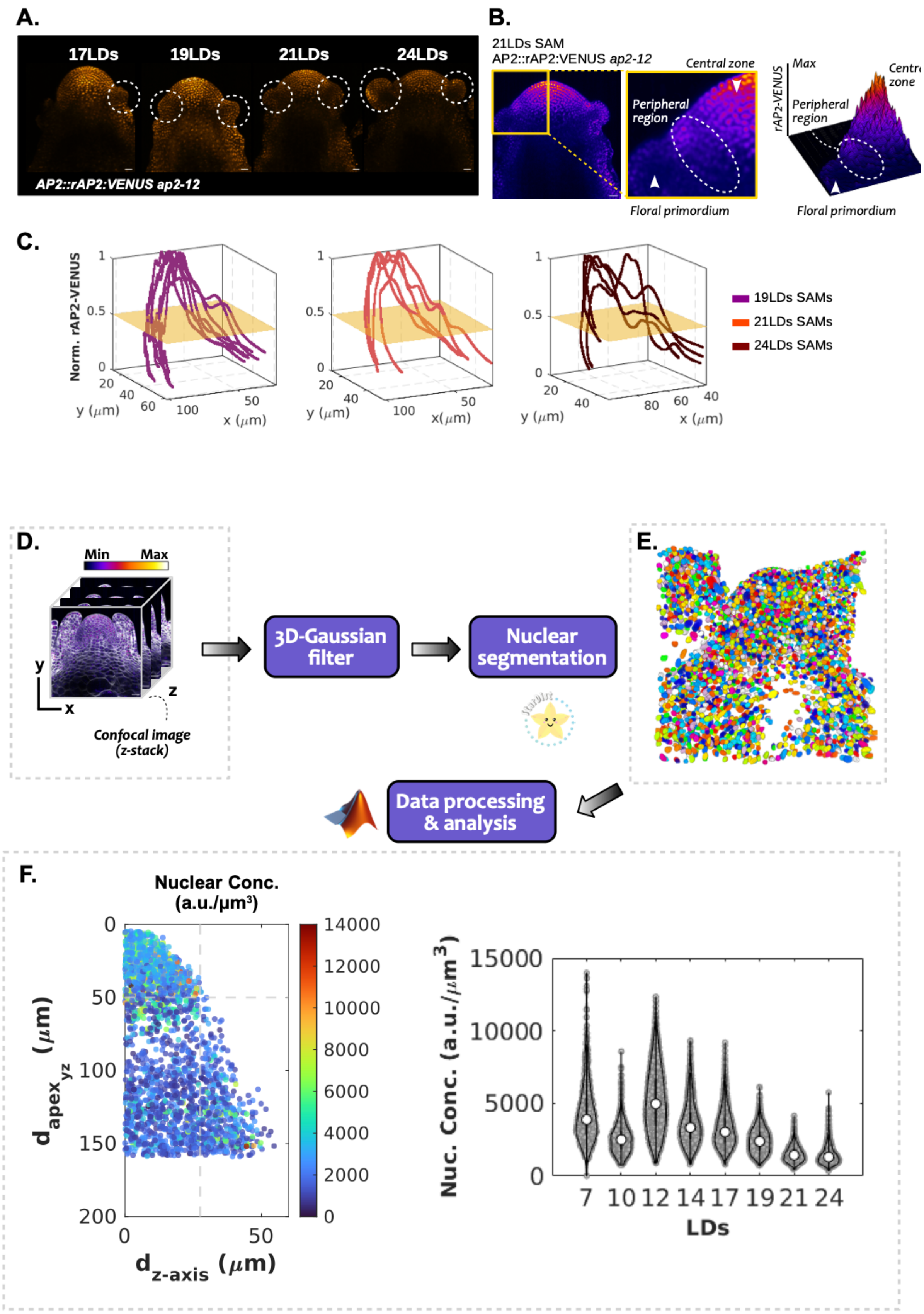
A.
17LDs
19LDs
21LDs
24LDs
AP2::rAP2:VENUS ap2-12
B.
21LDs SAM
AP2::rAP2:VENUS ap2-12
Central zone
Peripheral region
Floral primordium
rAP2-VENUS
Max
Peripheral region
Floral primordium
C.
Norm. rAP2-VENUS
y (µm)
x (µm)
19LDs SAMs
21LDs SAMs
24LDs SAMs
D.
Min
Max
Confocal image (z-stack)
3D-Gaussian filter
Nuclear segmentation
E.
Data processing & analysis
F.
Nuclear Conc. (a.u./µm³)
d_apex_yz (µm)
d_z-axis (µm)
Nuc. Conc. (a.u./µm³)
LDs

**Figure S10. Complementary tissue and single-cell nuclear quantification pipelines for temporal and spatial dynamics of AP2-VENUS and rAP2-VENUS expression at the SAM.** A) Expression of AP2::rAP2:VENUS at the SAM in *ap2*-12 under LDs. Scale bars = 20 µm. B) Example of AP2::rAP2:VENUS *ap2*-12 signal distribution in a 21LDs SAM, visualised in Fiji with Fire look-up table. An adjacent close-up region and its associated quantification in Fiji highlight the differences in signal density between the central zone and SAM periphery. C) Quantification of rAP2-VENUS intensity in different SAMs along an arch connecting opposite primordia (see Methods) at 19, 21, and 24LDs. Each intensity profile corresponds to a different SAM and was individually normalised. The yellow horizontal rectangles represent the 0.5 normalised value. D) Single-cell quantification pipeline description: fluorescence confocal microscopy images were obtained from the lateral side. In the image: z-stacks from a 12LD SAM containing the AP2::AP2:VENUS reporter. The membrane marker channel is displayed in white, and the AP2-VENUS fluorescence signal is coloured according to the Fire colour look-up table in Fiji. Scale bar = 20 µm. Acquisition parameters are described in Methods. A 3D-Gaussian filter (r = 2) in Fiji was applied to each z-stack to homogenise the signal and reduce over-segmentation problems. B) Nuclear segmentations were then performed using Stardist software and StarDist Plant Nuclei 3D ResNet model (see Methods). Single-nuclear segmentation data were analysed with a custom-made code in MATLAB. E) The 3D-paraboloid and the SAM-apex positional information, available from the tissue quantification pipeline (Figure S1), were integrated into the analysis to exclude nuclei from primordia and to provide a reference system. F) Single-cell nuclear AP2 concentration was then plotted as a function of both the vertical distance to the SAM apex and the horizontal distance from the vertical SAM longitudinal axis. Alternatively, all single-cell nuclear concentrations from all SAMs at the same time point can be plotted simultaneously and show the global trend in the temporal evolution of the signal.

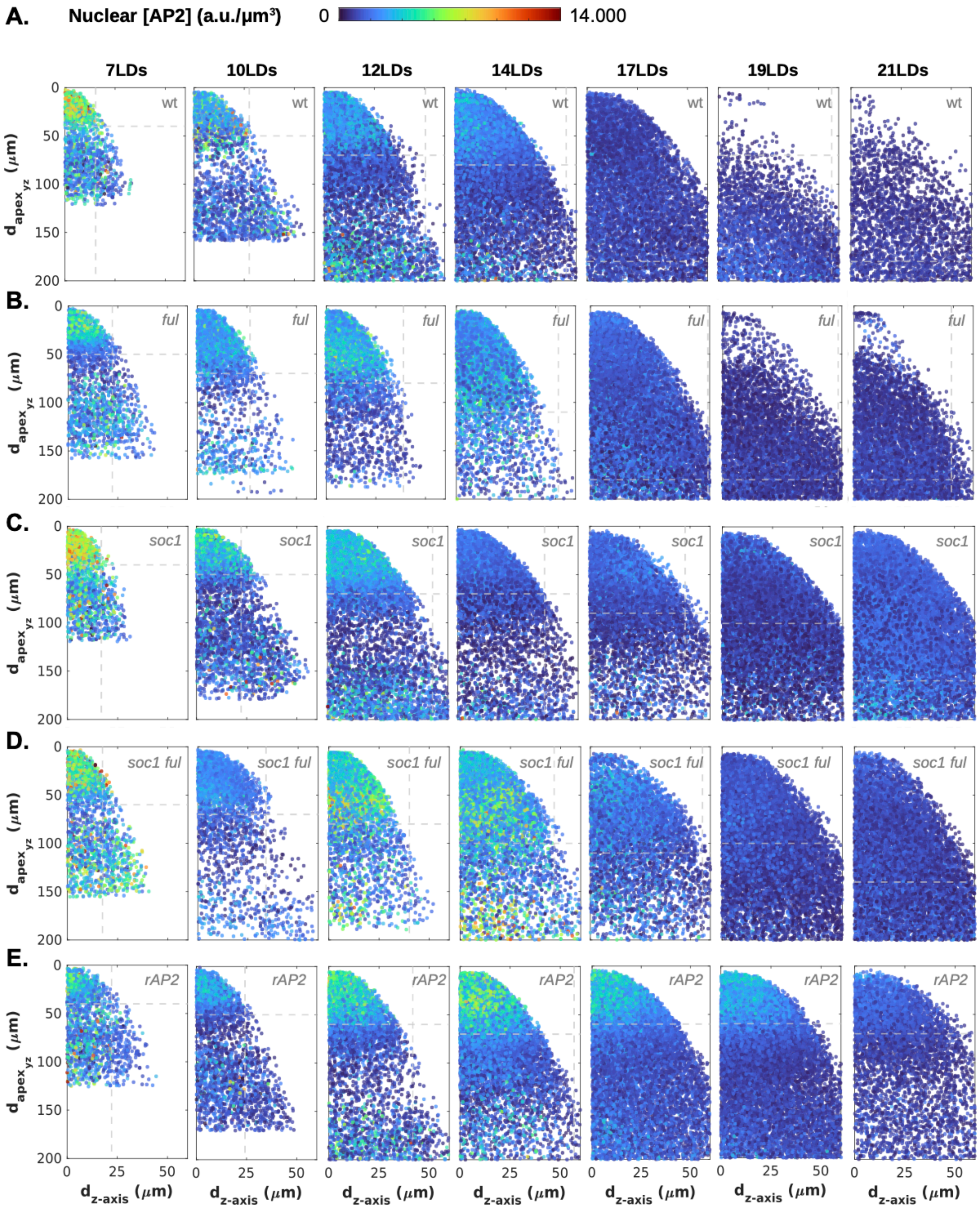

**Figure S11**. **Temporal and spatial quantification of single-cell nuclear protein concentration of AP2-VENUS and rAP2-VENUS during floral transition at the SAM in different genotypes.** A–E) Single-cell nuclear concentration from all SAMs as a function of the vertical distance to the apex ($d_{apex}$) and the horizontal distance to the SAM longitudinal axis ($d_{z\text{-}axis}$) at different time points (7-21LDs) during floral transition in wt (A), *ful* (B), *soc1* (C), *soc1 ful* (D), and rAP2 (E). Horizontal and vertical grey lines indicate the median height and width at each time point. For wt data, only one of the two available experiments was analysed.

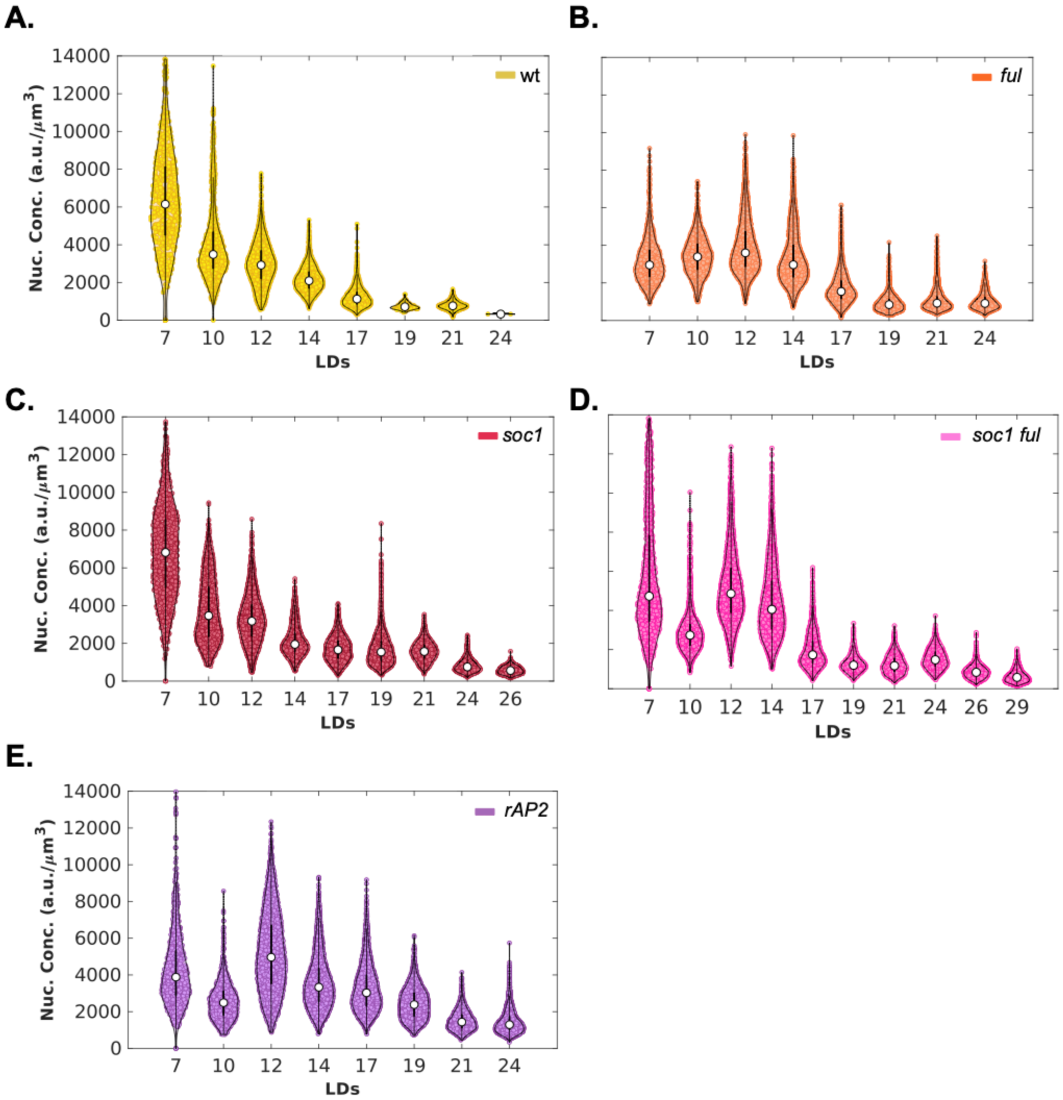

**Figure S12. Quantification of single-cell nuclear AP2-VENUS and rAP2-VENUS protein concentration temporal dynamics at the SAM during floral transition.** A–E) Collapsed single-cell nuclear concentration from all cells within the 50% characteristic domains in all SAMs at the same time point during floral transition in wt (A), *ful* (B), *soc1* (C), *soc1 ful* (D), and rAP2 (E).

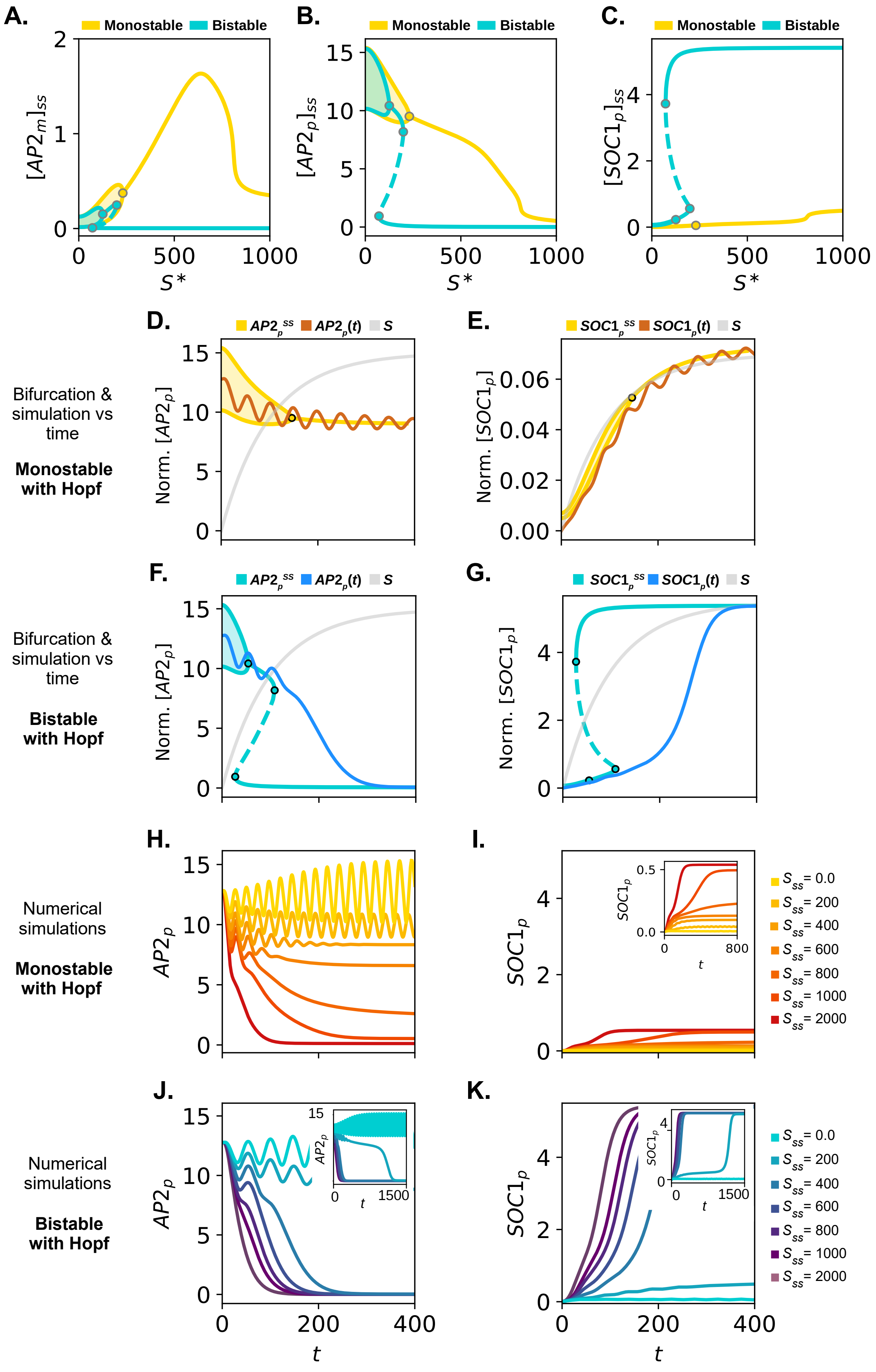
A.
Monostable Bistable
$[AP2_m]_{ss}$
$S*$
B.
Monostable Bistable
$[AP2_p]_{ss}$
$S*$
C.
Monostable Bistable
$[SOC1_p]_{ss}$
$S*$
D.
$AP2_p^{ss}$ $AP2_p(t)$ S
Norm. $[AP2_p]$
E.
$SOC1_p^{ss}$ $SOC1_p(t)$ S
Norm. $[SOC1_p]$
Bifurcation & simulation vs time
Monostable with Hopf
F.
$AP2_p^{ss}$ $AP2_p(t)$ S
Norm. $[AP2_p]$
G.
$SOC1_p^{ss}$ $SOC1_p(t)$ S
Norm. $[SOC1_p]$
Bifurcation & simulation vs time
Bistable with Hopf
H.
$AP2_p$
I.
$SOC1_p$
$S_{ss}$= 0.0
$S_{ss}$= 200
$S_{ss}$= 400
$S_{ss}$= 600
$S_{ss}$= 800
$S_{ss}$= 1000
$S_{ss}$= 2000
Numerical simulations
Monostable with Hopf
J.
$AP2_p$
$t$
K.
$SOC1_p$
$t$
$S_{ss}$= 0.0
$S_{ss}$= 200
$S_{ss}$= 400
$S_{ss}$= 600
$S_{ss}$= 800
$S_{ss}$= 1000
$S_{ss}$= 2000
Numerical simulations
Bistable with Hopf

**Figure S13. Analysis of monostable and bistable scenarios in the model with autoinhibition.** A–C) Bifurcation diagrams for the *AP2m* (A), *AP2p* (B), and *SOC1p* (C) model variables showing monostable (blue) and bistable scenarios (yellow). Coloured dots indicate Hopf (HB) and saddle-node (SN) bifurcations. Solid and dashed lines represent stable and unstable states, respectively; shaded regions enclosed by two solid lines represent stable limit cycles. D and E) *AP2p* (D) and *SOC1p* (E) model numerical simulation (dark orange) and $AP2p^{ss}$ (D) and $SOC1p^{ss}$ (E) (in yellow) steady state concentrations as a function of time in a monostable scenario. F and G) *AP2p* (F) and *SOC1p* (G) model numerical simulation (dark blue) and $AP2p^{ss}$ (F) and $SOC1p^{ss}$ (G) (in cyan) steady state concentrations as a function of time in a bistable scenario. In D–G, the normalised age signal (*S*) is indicated as a grey line. Coloured dots indicate Hopf and saddle-node bifurcation points. H and I) Numerical simulations showing the dynamics of *AP2p* (H) and *SOC1p* (I) for different $S_{ss}$ that lead to monostable scenarios. F and G) Numerical simulations showing the dynamics of *AP2p* (F) and *SOC1p* (G) for different $S_{ss}$ that lead to bistable scenarios. The insets show the dynamics in a longer time window, whereby a simulation case undergoing a long critical slowing down phase can be seen. See Suppl. Tables 1 and 2 for parameter values.

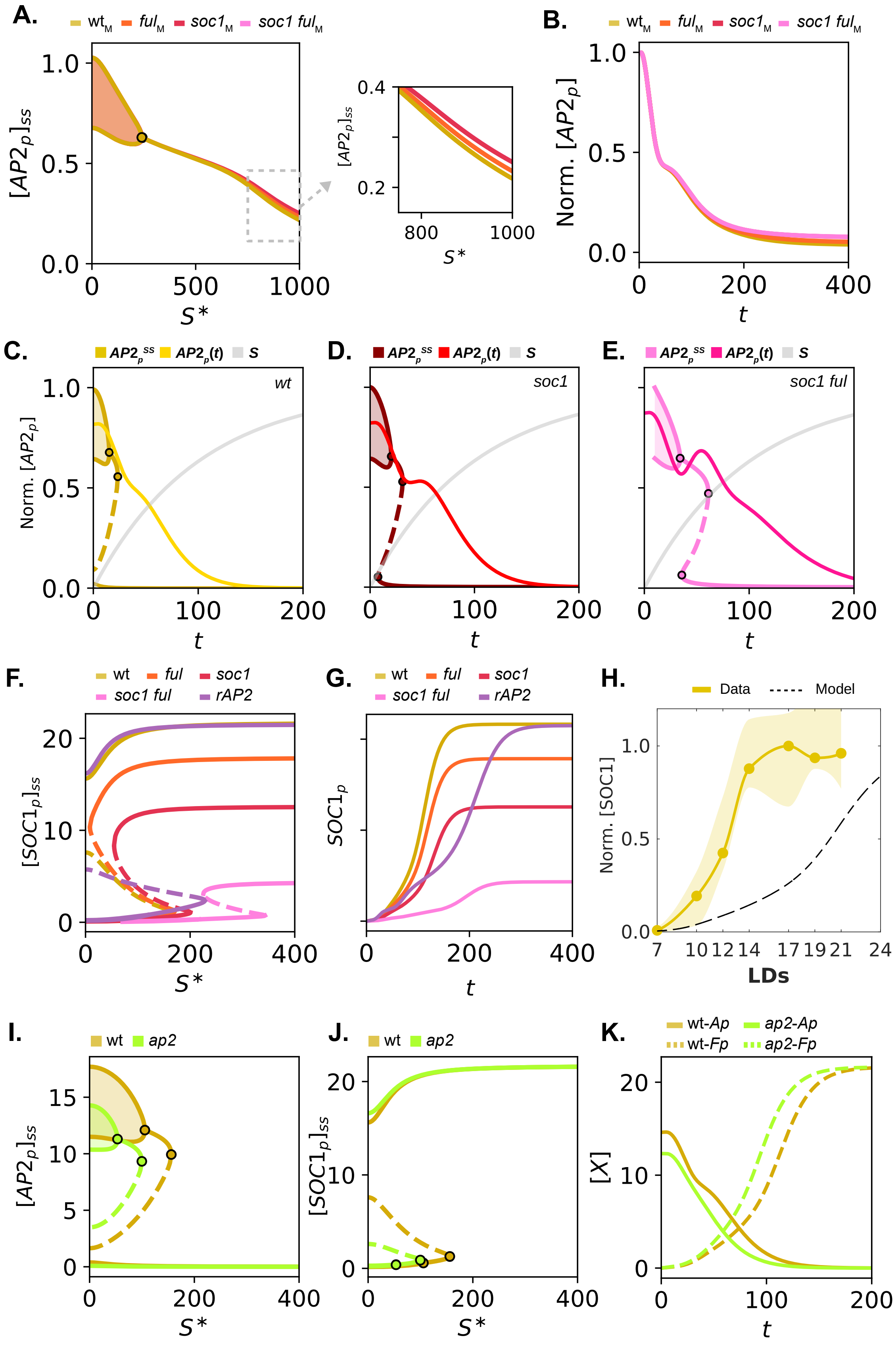
A.
wt_M ful_M soc1_M soc1 ful_M
[AP2_p]_ss
S*
B.
Norm. [AP2_p]
t
C.
AP2_p^ss AP2_p(t) S
wt
Norm. [AP2_p]
D.
soc1
E.
soc1 ful
F.
wt ful soc1 soc1 ful rAP2
[SOC1_p]_ss
G.
SOC1_p
H.
Data Model
Norm. [SOC1]
LDs
I.
wt ap2
[AP2_p]_ss
J.
[SOC1_p]_ss
K.
wt-Ap ap2-Ap wt-Fp ap2-Fp
[X]

**Figure S14. Extended analysis of genotype dynamics in the model with autoinhibition.** A) Normalised *AP2p* bifurcation diagrams for wt, *ful*, *soc1,* and *soc1 ful* genotypes in a monostable region. Coloured dots indicate Hopf (HB) bifurcations. Solid lines represent stable states; shaded regions enclosed by two solid lines represent stable limit cycles. B) Numerical simulations of genotypes in A) in a monostable region. C–E) Normalised *AP2p* model numerical simulation (non-grey continuous line) and $AP2p^{ss}$ (line with dashed region) concentrations as a function of time for wt (C), *soc1* (D), and *soc1 ful* (E) genotypes. The normalised age signal (*S*) is indicated as a grey line. Coloured-filled dots indicate Hopf and saddle-node bifurcation points. Normalisation was performed for each variable individually. F and G) *SOC1p S*-bifurcation diagram (F) and numerical simulation (G) for wt, *ful*, *soc1*, *soc1 ful,* and rAP2, associated with the results shown in Figure 5. In (F), solid and dashed lines represent stable and unstable states, respectively. H) SOC1 wt-temporal dynamics comparison between experimental (continuous yellow line) and model (dashed black line). I-K) *AP2p* (I) and *SOC1p* (J) bifurcation diagrams and numerical simulations (K) as a function of time for wt and *ap2*. See Suppl. Tables 1 and 2 for parameter values.

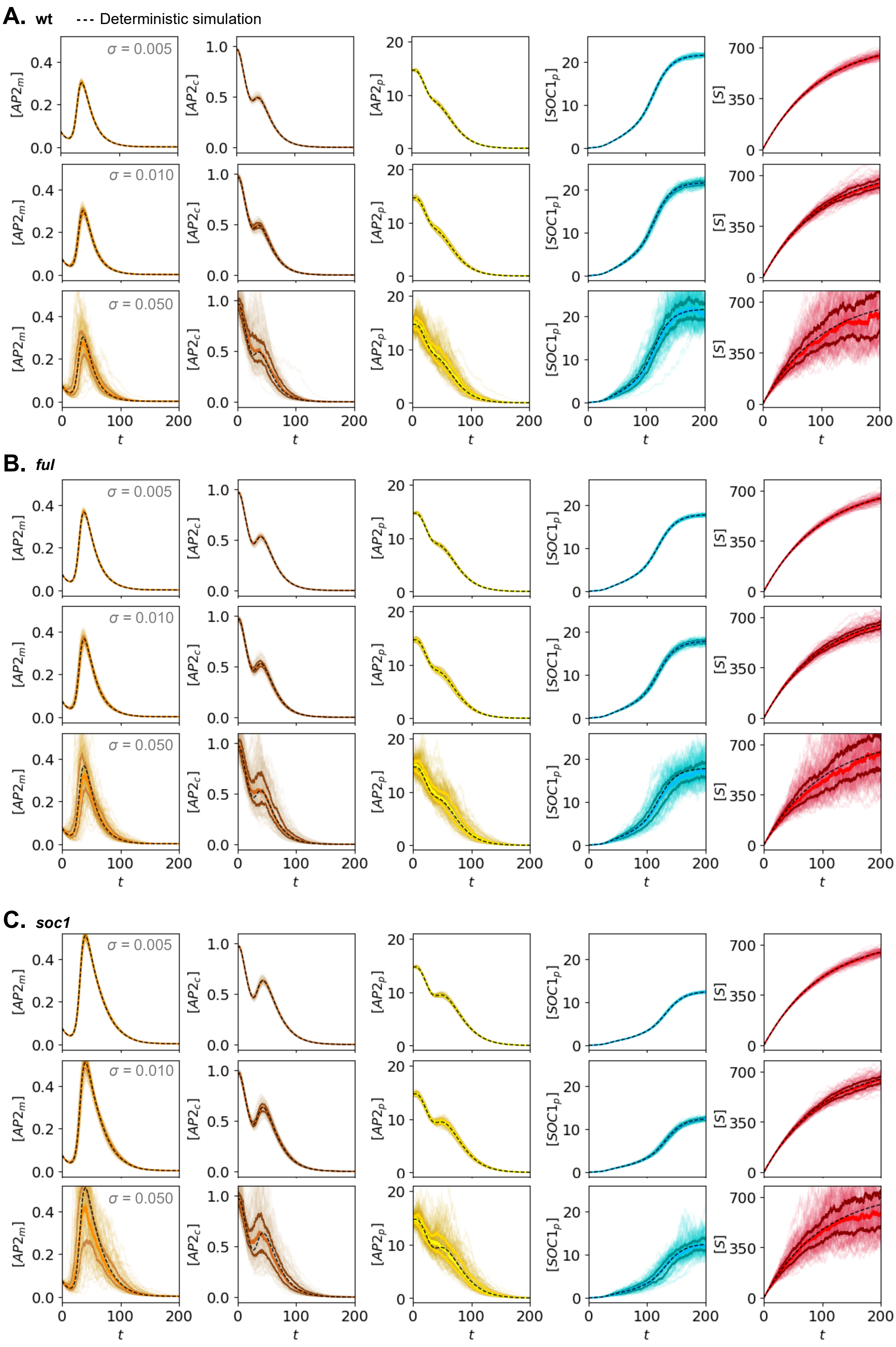

A. wt
--- Deterministic simulation
B. ful
C. soc1
σ = 0.005
σ = 0.010
σ = 0.050
[AP2m]
[AP2c]
[AP2p]
[SOC1p]
[S]
t

**Figure S15. Stochastic numerical simulations under intrinsic noise in the extended toggle-switch model with AP2 autoinhibition for wt, *ful* and *soc1* genotypes.** A–C) Temporal dynamics of model variables *Am* (column 1), *Ac* (column 2), *Ap* (column 3), *Fp* (column 4), and *S* (column 5) for wt (A), *ful* (B), and *soc1* (C) genotypes. Simulations (N = 100) were performed across three noise intensities (σ = 0.005, 0.01, and 0.05). Thick coloured lines represent the median trajectory flanked by first and third quartiles. Black dashed lines denote the corresponding deterministic simulations. Saddle-node bifurcation points occur at $S_{SN}$ = [156, 171, 200], for wt, *ful*, and *soc1* genotypes, respectively. The steady state of *S* reached in the simulations is $S^*$=750. See Methods and Suppl. Table 2 for further details.

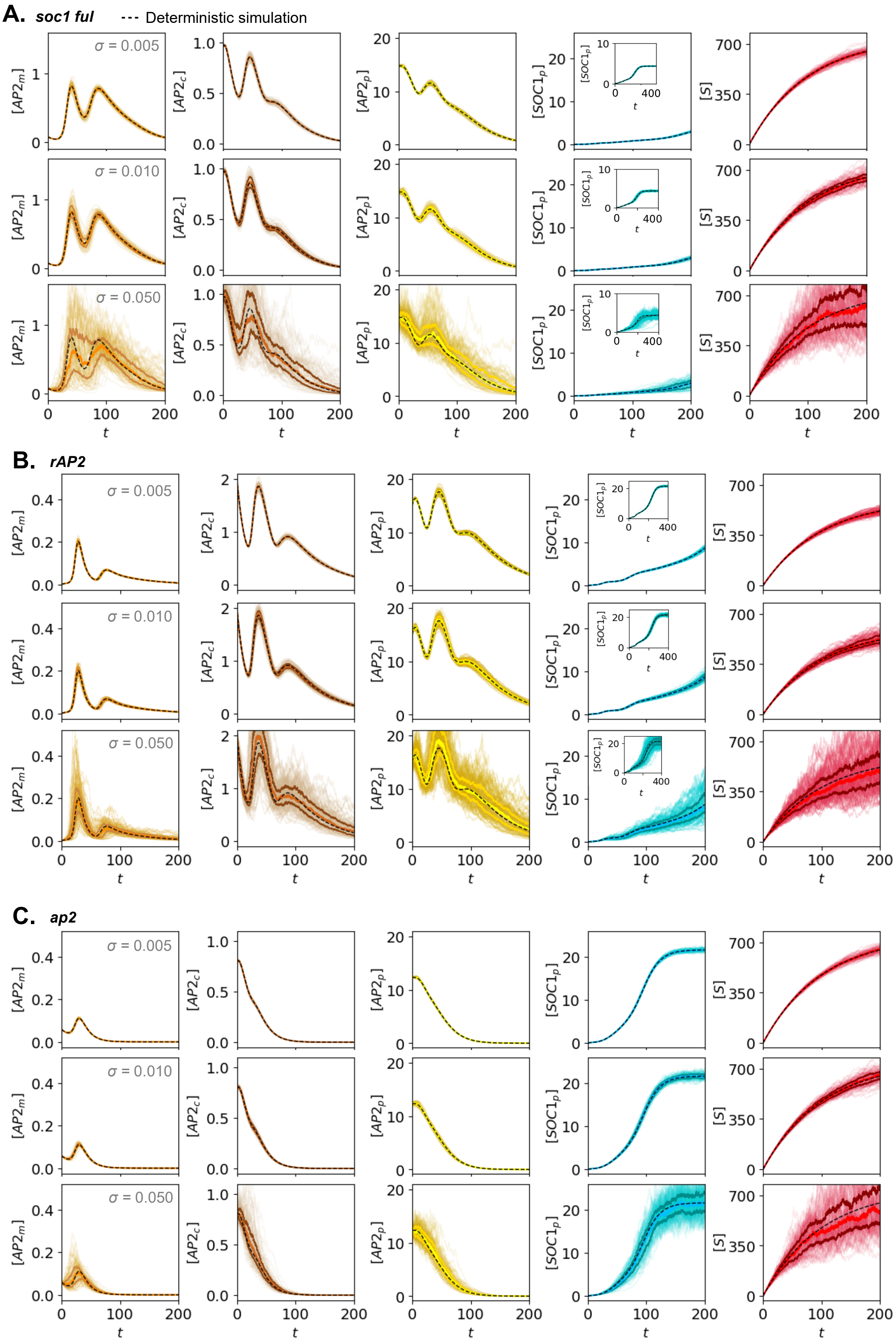
A. soc1 ful
--- Deterministic simulation
σ = 0.005
σ = 0.010
σ = 0.050
[AP2m]
[AP2c]
[AP2p]
[SOC1p]
[S]
t
B. rAP2
σ = 0.005
σ = 0.010
σ = 0.050
C. ap2
σ = 0.005
σ = 0.010
σ = 0.050

**Figure S16. Stochastic numerical simulations under intrinsic noise in the extended toggle-switch model with AP2 autoinhibition for *soc1 ful*, *rAP2* and *ap2* genotypes.**
A–C) Temporal dynamics of model variables *Am* (column 1), *Ac* (column 2), *Ap* (column 3), *Fp* (column 4), and *S* (column 5) for *soc1 ful* (A), *rAP2* (B), and *ap2* (C) genotypes. Simulations (N = 100) were performed across three noise intensities (σ = 0.005, 0.01, and 0.05). Thick coloured lines represent the median trajectory flanked by first and third quartiles. Black dashed lines denote the corresponding deterministic simulations. Saddle-node bifurcation points occur $S_{SN}$ = [343, 228, 99.8], for *soc1 ful*, *rAP2*, and *ap2* genotypes, respectively. The steady state of *S* reached in the simulations is *S**=750 for *soc1 ful* and *ap2* and *S**=600 for *rAP2*. See Methods and Suppl. Table 2 for further details.

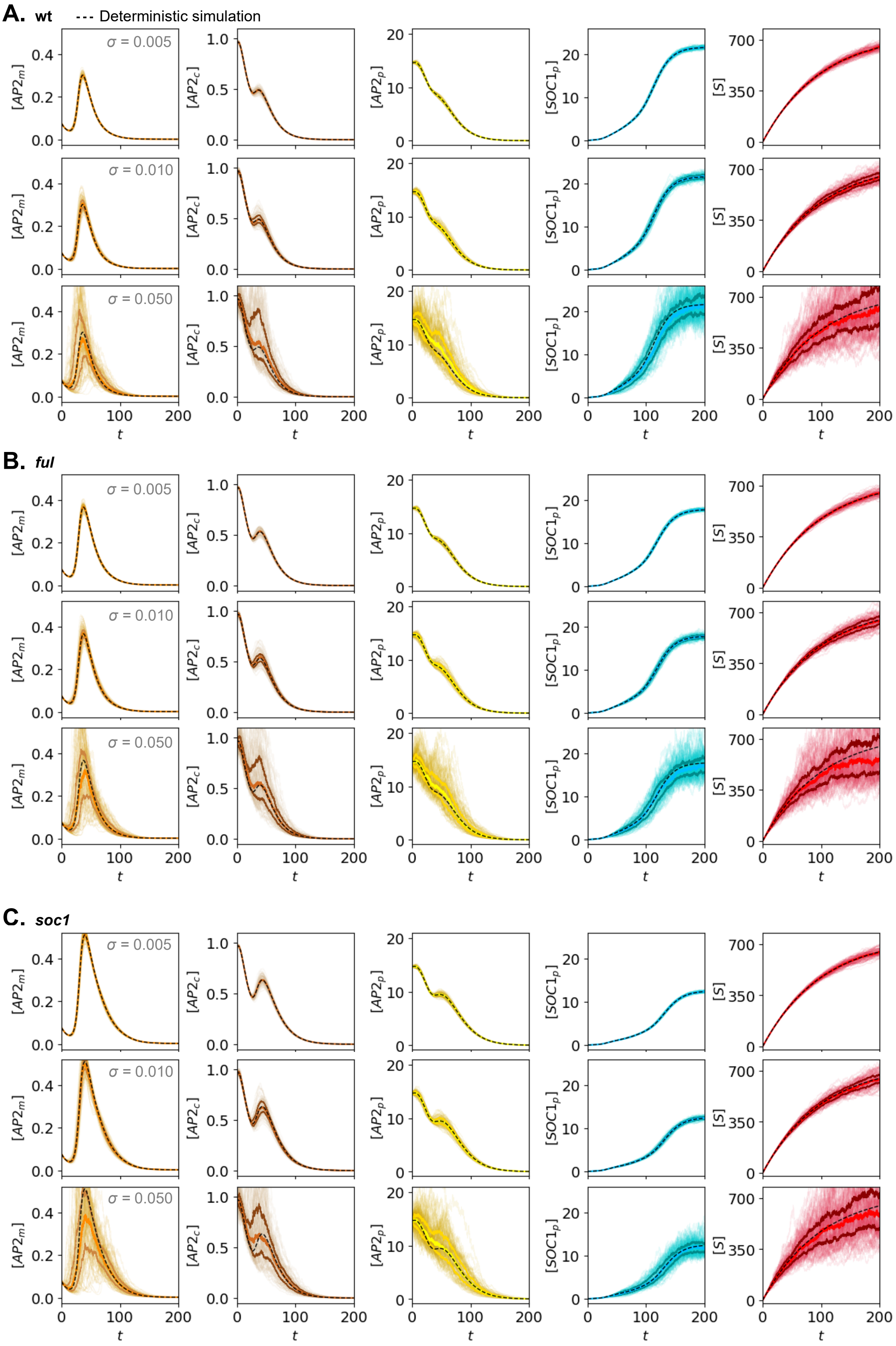
A. wt
Deterministic simulation
B. ful
C. soc1
σ = 0.005
σ = 0.010
σ = 0.050
[AP2m]
[AP2c]
[AP2p]
[SOC1p]
[S]
t

**Figure S17. Stochastic numerical simulations under extrinsic noise in the extended toggle-switch model with AP2 autoinhibition for wt, *ful* and *soc1* genotypes.** A–C) Temporal dynamics of model variables *Am* (column 1), *Ac* (column 2), *Ap* (column 3), *Fp* (column 4), and *S* (column 5) for wt (A), *ful* (B), and *soc1* (C) genotypes. Simulations (N = 100) were performed across three noise intensities (σ = 0.005, 0.01, and 0.05). Thick coloured lines represent the median trajectory flanked by first and third quartiles. Black dashed lines denote the corresponding deterministic simulations. Saddle-node bifurcation points and steady state of *S* as in Figure S15. See Methods and Suppl. Table 2 for further details.

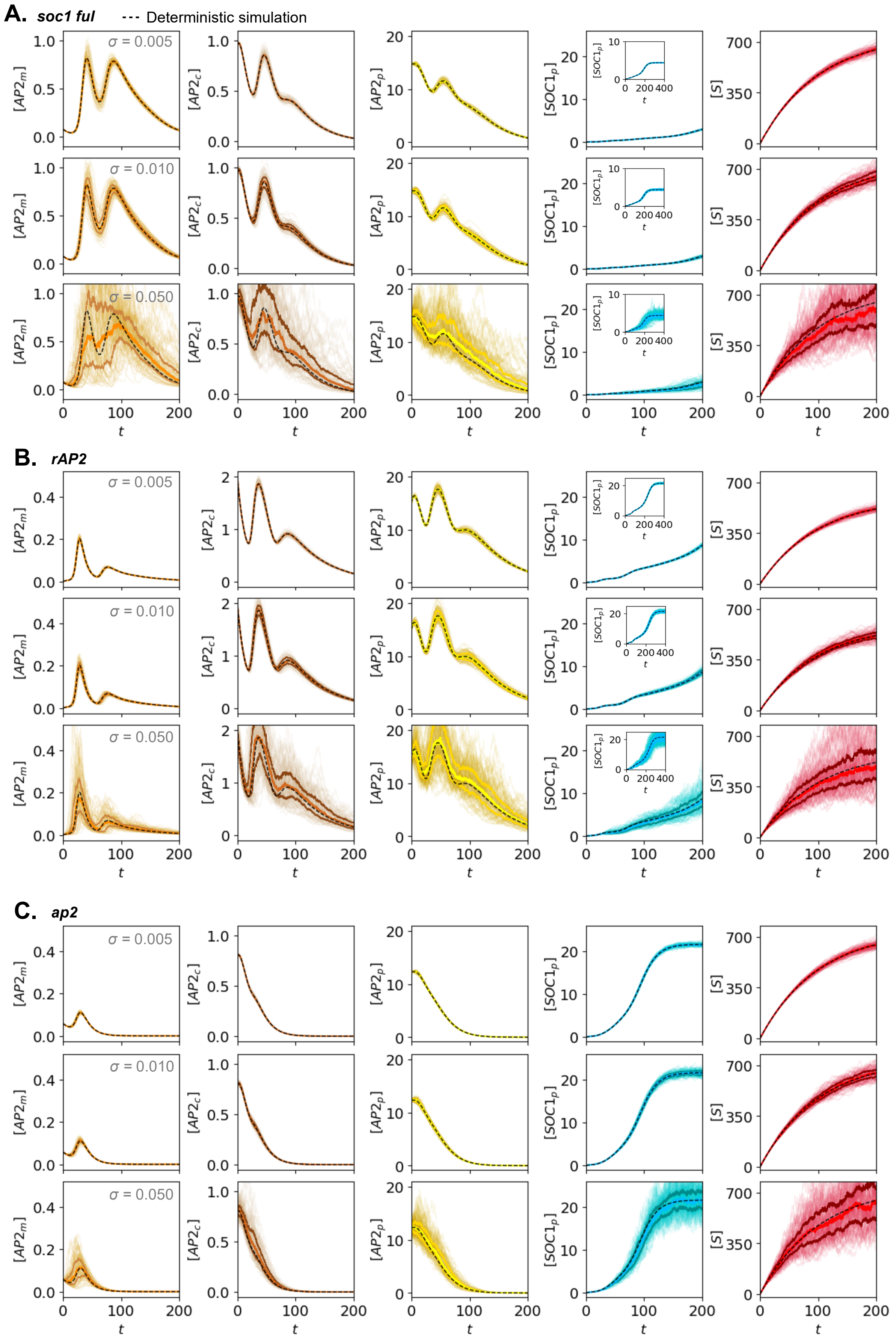
A. soc1 ful
--- Deterministic simulation
B. rAP2
C. ap2
σ = 0.005
σ = 0.010
σ = 0.050
[AP2m]
[AP2c]
[AP2p]
[SOC1p]
[S]
t

**Figure S18. Stochastic numerical simulations under extrinsic noise in the extended toggle-switch model with AP2 autoinhibition for *soc1 ful*, *rAP2* and *ap2* genotypes.**
A–C) Temporal dynamics of model variables *Am* (column 1), *Ac* (column 2), *Ap* (column 3), *Fp* (column 4), and *S* (column 5) for *soc1 ful* (A), *rAP2* (B), and *ap2* (C) genotypes. Simulations (N = 100) were performed across three noise intensities (σ = 0.005, 0.01, and 0.05). Thick coloured lines represent the median trajectory flanked by first and third quartiles. Black dashed lines denote the corresponding deterministic simulations. Saddle-node bifurcation points and steady state of *S* as in Figure S16. See Methods and Suppl. Table 2 for further details.

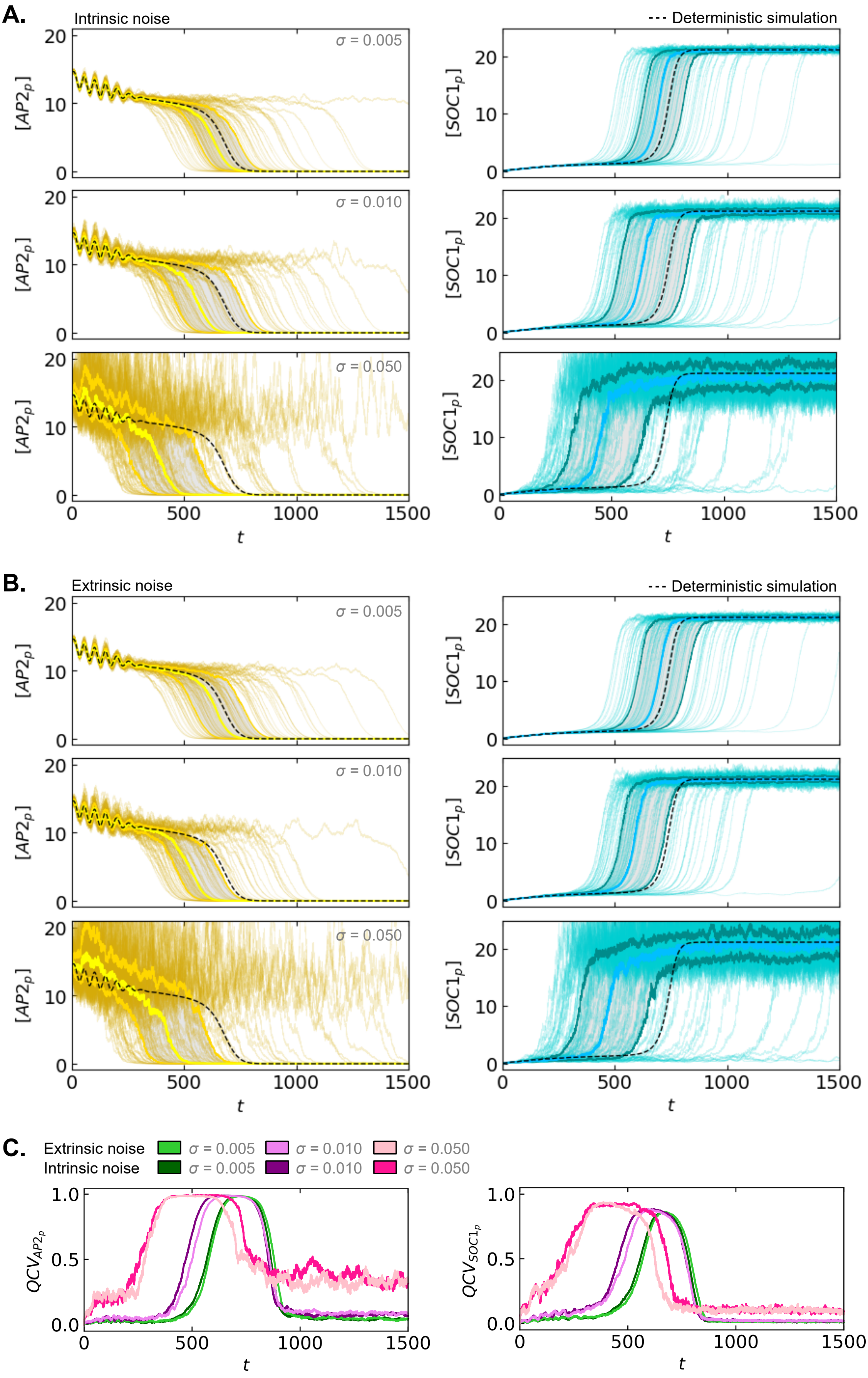
A.
Intrinsic noise
--- Deterministic simulation
σ = 0.005
σ = 0.010
σ = 0.050
[AP2p]
[SOC1p]
t
B.
Extrinsic noise
--- Deterministic simulation
σ = 0.005
σ = 0.010
σ = 0.050
C.
Extrinsic noise
Intrinsic noise
σ = 0.005
σ = 0.010
σ = 0.050
QCV_AP2p
QCV_SOC1p
t

**Figure S19. Stochastic numerical simulations in which the age variable saturated closer to the saddle node bifurcation under intrinsic and extrinsic noise in the extended toggle-switch model.** A and B) Temporal dynamics of model variables AP2p (column 1) and SOC1p (column 2) under intrinsic (A) and extrinsic noise (B). Simulations (N = 100) were performed across three noise intensities ($\sigma$ = 0.005, 0.01, and 0.05). Thick coloured lines represent the median trajectory, flanked by first and third quartiles. Black dashed lines denote the corresponding deterministic simulations. C) Quartile coefficient of variation (QCV) for AP2p (left) and SOC1p (right) from stochastic numerical simulations for the extrinsic and intrinsic noise intensities shown in A and B. In these simulations, the steady state of the signal is closer to the saddle-node bifurcation, and hence there is a stronger influence of the ghost attractor in the deterministic limit than in Figures S15–S18. Saddle-node bifurcation occurs at $S_{SN}$=156.01, and the steady state of $S$ reached in the simulations is $S^*$=175. See Methods and Suppl. Table 2 for further details.

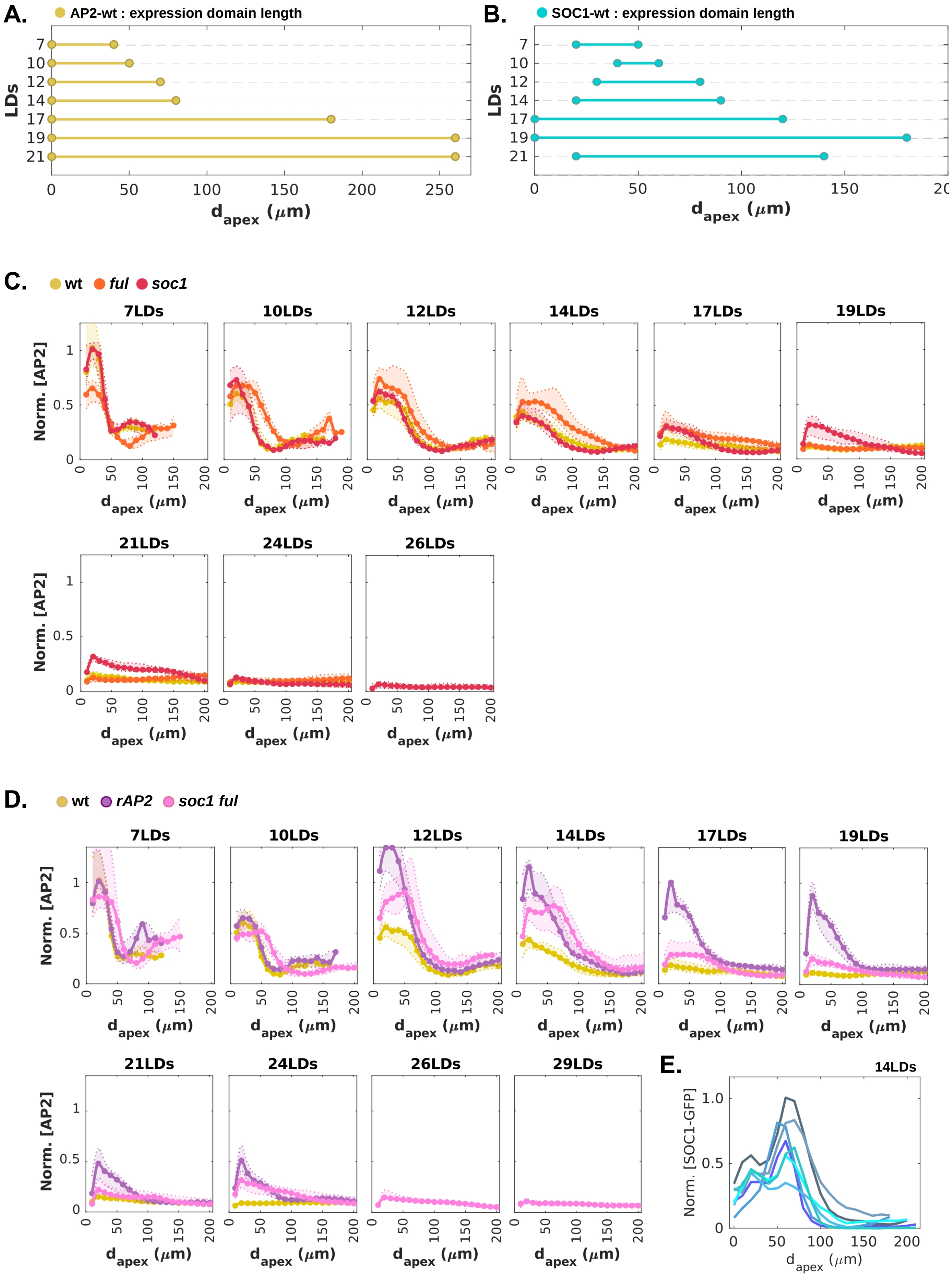

**Figure S20. Extended spatial dynamics data for AP2-Venus and SOC1-GFP.** A and B) Characteristic domain ranges for AP2-VENUS (A) and SOC1-GFP (B) of the median protein expression profiles for each time point during floral transition in wt-SAMs. C) Spatial protein dynamics of AP2-VENUS in wt, *ful,* and *soc1* during floral transition at the SAM. D) Spatial protein dynamics of AP2-Venus in wt, *soc1 ful,* and rAP2 during floral transition at the SAM.

Expression profiles of *soc1* were quantified from published data[47]. $d_{apex}$ represents the distance with respect to the SAM apex. Normalisation was performed with respect to the maximum value reached in wt. Individual normalisation was performed for AP2:rAP2::VENUS fluorescence because the transgene is inserted in a different genomic position, and the fluorescence intensity signal is therefore not directly comparable with AP2:AP2::VENUS of the other genotypes. Continuous lines represent the median value. Dashed lines indicate first and third quantiles. E) Single SAM spatial SOC1-GFP protein dynamics in wt versus distance to the SAM apex at 14LDs. Normalisation was performed with respect to the maximum value reached among all SAMs analysed at this time point.

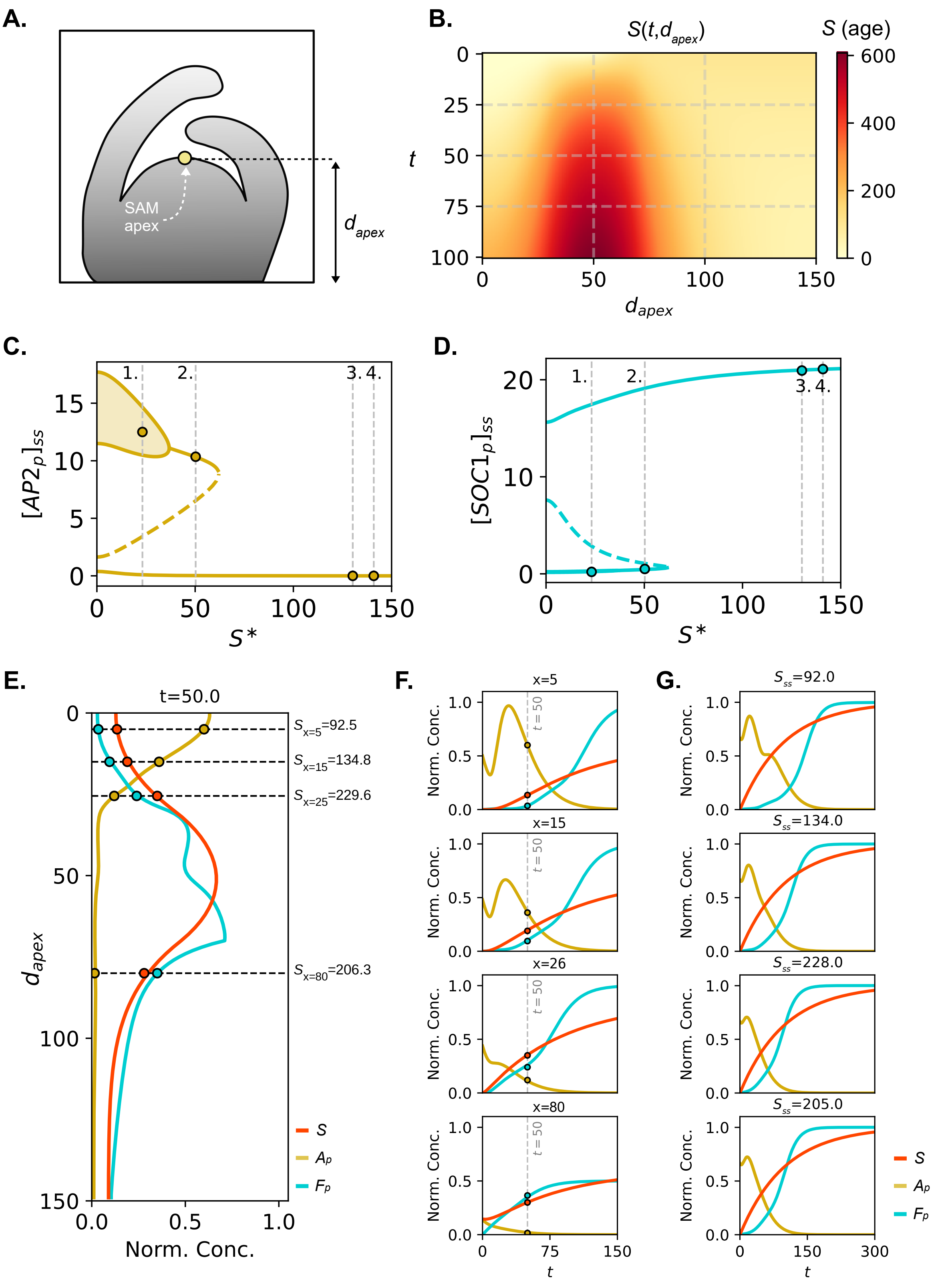

**Figure S21. Additional analysis of the spatial model.** A) Schematic diagram of a meristem indicating the apex and the distance to it along SAM's longitudinal axis. B) Heat map showing the simulated age-signal (S) spatio-temporal pattern in the extended model, $S(t,d_{apex})$. C and D) $AP2_p$ (C) and $SOC1_p$ (D) age-bifurcation diagrams. Coloured dots indicate $AP2_p$ and $SOC1_p$ values for different $S$ coordinates (grey vertical dashed lines) corresponding to the different scenarios shown in Figures 6C and 6D. Solid and dashed lines represent stable and unstable states, respectively. E) Numerically simulated spatial dynamics of $A_p$, $F_p$, and $S$ wt-model variables at t = 50 as a function of the distance to the apex. F) Temporal numerical simulations at different SAM positions (x = 5, 15, 25.5, and 80) along the SAM longitudinal axis. All variables were normalised by their global maximum. Coloured dots indicate the correspondence with panel (E). G) Temporal numerical simulations at different SAM positions (x = 5, 15, 25.5 and 80) along the SAM longitudinal axis, where the maximum $S$ value reached corresponds to the red-filled point in F. Normalisation of $A_p$ and $F_p$ was performed by dividing by their maximum values reached across the four displayed scenarios ($S_{ss}$ = 92, 134, 228 and 205). For graphical clarity, $S$ was normalised individually within each scenario. See Suppl. Tables 1 and 2 for parameter values and Suppl. Tables 3 and 4 for information about normalisations.

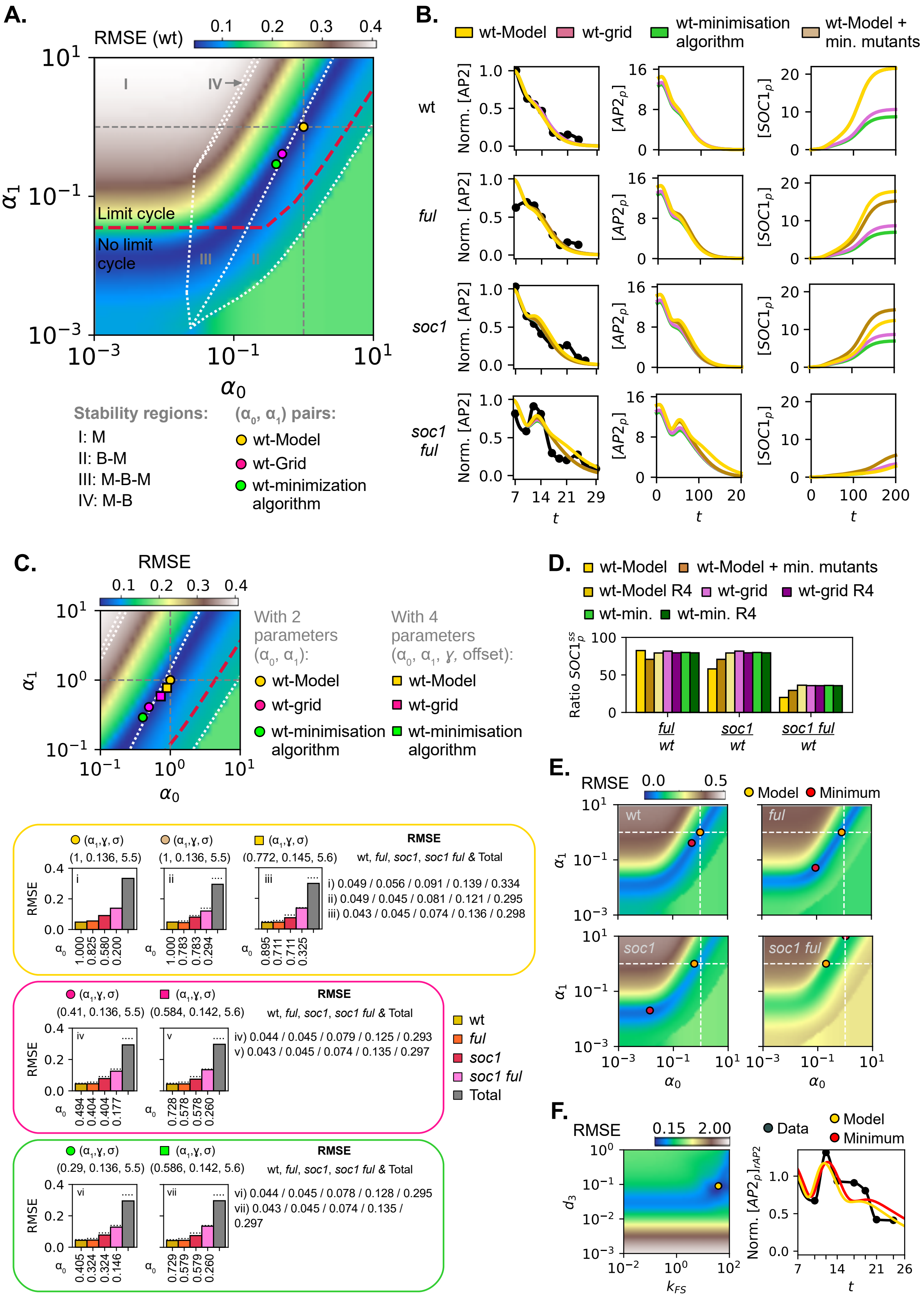

**Figure S22. Model parametrisation and parameter optimisation.** A) Heatmap of the ($\alpha_0$,$\alpha_1$) plane showing the root mean square error (RMSE) between AP2-VENUS data and Model 2 simulations for wt. Coloured dots indicate three parameter sets: yellow, Model 2 values used in the main text; pink, grid-minimum RMSE; and green, best fit via Nelder-Mead minimisation.

Roman numbers denote stability regimes along S*: monostable (I:M); bistable to monostable (II:B-M); two saddle-node bifurcations (III:M-B-M); monostable to bistable (IV: M-B) where $S^*$ is insufficient to reach the second saddle-node. The red line marks the limit cycle boundary. B) Temporal dynamics for wt and floral activator mutant genotypes associated with the parameter sets in A. For the mutant genotypes in pink, green, and light brown cases, $\alpha_1$ was fixed to the corresponding wt value in A, and $\alpha_0$ was optimised via Nelder-Mead minimisation of the RMSE under the biological constraint $\alpha_0^{\mathrm{wt}} > \alpha_0^{ful} > \alpha_0^{soc1} > \alpha_0^{soc1\ ful}$. First column: normalised AP2-VENUS data (black dots) and *Ap* simulations. Second and third columns: non-normalised inhibitor and activator model variable dynamics. C) Multi-parameter RMSE optimisation heatmap. The ($\alpha_0$,$\alpha_1$) plane shows how initial pairs (circles) shift after Nelder-Mead minimisation of the RMSE for AP2-wt model simulations on the quadruple ($\alpha_0$,$\alpha_1$,$\gamma$, offset), depicted by squares. Subsequent mutants optimisations as described in B. D) Predicted ratios (%) of steady-state *SOC1p* values in mutants versus wt for each parametrisation in C. E) Individual RMSE heat maps in the ($\alpha_0$,$\alpha_1$)-plane for AP2 wt, *ful*, *soc1*, and *soc1 ful* model simulations (Model 2) and their respective AP2-VENUS datasets. F) Optimisation of *rAP2* genotype. Left: RMSE heat map in the ($d_3$,$k_{FS}$)-plane. Right: comparison of rAP2 temporal dynamics using Model 2 parameters (yellow) and the ($d_3$,$k_{FS}$) pair yielding the global minimum RMSE value in the grid (red) versus rAP2-VENUS experimental data. See Methods and Suppl. Tables 2 and 4 for further information.

### Supplementary Tables

**Table S1. Parameter values and model specifications from the main figures**

| **Figure** | *Parameter and/or model specifications* |
|---|---|
| Fig. 2 (M1) | E–G) $\alpha_1$=0.81, $\alpha_4$=0.3, $\alpha_0$=0.22, $d_1$=0.08, $d_4$=0.08, $d_s$=0.008, $k_{AS}$=160.0, $k_{AF}$=0.9, $k_{FS}$=80.0, $k_{FA}$=1.0, $n$=2, and $\beta_s$=1.75.<br>H and I) Same as B with $\alpha_0$=0.34.<br>J) Same as H with $\beta_s$=1.535 (to be close to the saddle node).<br>K and L) Parameters wt-genotype: $\alpha_1$=0.58, $\alpha_0$=0.35, $\alpha_4$=0.30, $d_1$=0.08, $d_4$=0.08, $\beta_s$=1.3, $d_s$=0.008, $k_{AS}$=160.0, $k_{AF}$=0.9, $k_{FS}$=80.0, $k_{FA}$=1.0, $n$=2.<br>M) Same as J. To fit data and numerical simulation, $t_{exp}$= $t_{num}$·(10.75/125)+6.25.<br><br>Initial conditions for numerical simulations: $(A, F, S)\|_{t=0}$ = $(A^{ss(S=0)}, 0, 0)$, where $A^{ss(S=0)}$ is the AP2 steady state value reached when $S$ = 0 (i.e., $\beta_s$=0). |
| Fig. 3 (M1) | D, E, and J) Parameters as in Fig. 2J-L.<br>$\alpha_0^{ful}$=0.95·$\alpha_0^{wt}$, $\alpha_0^{soc1}$=0.95·$\alpha_0^{wt}$, $\alpha_0^{soc1ful}$=0.82·$\alpha_0^{wt}$.<br>E) Nullclines were computed at the wt-saddle node $S$-value ($S_{ss}^{wt}$=138.25).<br><br>Initial conditions: $(A, F, S)\|_{t=0}$ = $(A^{ss(S=0)}, 0, 0)$ where $A^{ss(S=0)}$ is the AP2 steady state value reached when $S$ = 0 (i.e., $\beta_s$=0). |
| Fig. 4 (M2) | Parameters wt-genotype: $\alpha_1$=1.0, $\alpha_2$=1.0, $\alpha_3$=1.0, $\alpha_0$=1.0, $\alpha_4$=0.3, $d_1$=0.1, $d_2$=0.065, $d_3$=0.065, $d_4$=0.06, $\beta_s$=7.5, $d_s$=0.01, $k_{AA}$=8, $k_{AS}$=80.0, $k_{AF}$=0.2, $k_{FS}$=50.0, $k_{FA}$=1.5, $n$=10, $n_2$=2.<br>In D, to fit the data and the numerical simulation, $t_{exp}$= $t_{num}$·(17/125)+5.5.<br><br>Initial conditions: $(Am, Ac, Ap, Fp, S)\|_{t=0}$ = $(Am^{ss(S=0)}, Ac^{ss(S=0)}, Ap^{ss(S=0)}, 0, 0)$ where $Ax^{ss(S=0)}$ is the $A$x steady state value reached when $S$ = 0 (i.e., $\beta_s$=0). |
| Fig. 5 (M2) | Parameters as in Fig. 4<br>$\alpha_0^{ful}$=0.825, $\alpha_0^{soc1}$=0.580, $\alpha_0^{soc1ful}$=0.200, $\alpha_1^{ap2}$=0.250;<br>For rAP2, $(\alpha_0^{rAP2},\alpha_1^{rAP2})=(\alpha_0^{wt},\alpha_1^{wt})$, $d_3$=0.09, $k_{FS}$=38.0, $\beta_s$=6.0, and no inhibition of age on $Ac$.<br><br>Initial conditions: $(Am, Ac, Ap, Fp, S)\|_{t=0}$ = $(Am^{ss(S=0)}, Ac^{ss(S=0)}, Ap^{ss(S=0)}, 0, 0)$, where $Ax^{ss(S=0)}$ is the $A$x steady state value reached when $S$ = 0 (i.e., $\beta_s$=0). For rAP2, $(Am, Ac, Ap, Fp, S)\|_{t=0}$ = (0, 2, 15, 0, 0). In the comparison with the experimental data, normalisation is done with respect to $Ap^{t=0}$. |

| | |
|---|---|
| Fig. 6 (M3) | Parameters as those of wt in Fig. 4.<br>$k_{AS}$=15.0, $\beta_s$=15, *D*=1 and *dt* = 0.01. L = 150 and *dx* = L/N with N = 150.<br>Zero flux boundary conditions.<br>$\beta_s > 0$ if x ∊ [30,70] or if *S*(x,t) > 0<br>$\alpha_i$=$\alpha_i$·(1+0.1·\|x-70\|)^(-1) for i =1 and i =3 if x > 70.<br><br>Initial conditions: $(Am, Ac, Ap, Fp, S)\|_{t=0}$ = (0, 0, *Ap*(x,0), 0, *S*(x,0)) where *Ap*(x,0) = 15*e*^(-0.0002$x^2$) and *S*(0,x≥50) = -0.25($x^2$) + 35x - 1125 & *S*(0,x≥70) = 100. In E, $\beta_s$=0.23 (1), 0.50 (2), 0.78 (3), and 1.39 (4). |

**Table S2. Parameter values and model specifications from supplementary figures**

| **Figure** | *Parameter and/or model specifications* |
|---|---|
| Fig. S3 (M1) | Same as Fig. 2E.<br>A-C) $\alpha_0$=0.22.<br>D-F) $\alpha_0$=0.34.<br>G-I) $\alpha_0$=0.34 and $\beta_s$=1.535.<br><br>Initial conditions: $(A, F, S)\|_{t=0}$ = $(A^{ss(S=0)}$, 0, 0) where $A^{ss(S=0)}$ is the AP2 steady state value reached when $S$ = 0 (i.e., $\beta_s$=0). |
| Fig. S4 (M1) | Same as Fig. 2E.<br>A) $\alpha_0$=0.22.<br>B and C) $\alpha_0$=0.34.<br>E and F) $\beta_s$=1.4 and $\alpha_0$=[0.10, 0.25, 0.40, 0.50, 0.75, 1.00].<br>G and H) $\beta_s$=2.4, $\alpha_0$=0.22, and $\alpha_1$=[0.40, 0.60, 0.80, 1.00, 1.20, 1.40].<br>Initial conditions: $(A, F, S)\|_{t=0}$ = $(A^{ss(S=0)}$, 0, 0) where $A^{ss(S=0)}$ is the AP2 steady state value reached when $S$ = 0 (i.e., $\beta_s$=0). |
| Fig. S5 (M1) | Same as Fig. 2K and 2L.<br>A) --<br>B and C) Different $S_{ss}$ values.<br>D and E) $\beta_s$=1.75. A N-square ($\alpha_0$, $\alpha_1$)-grid ($\alpha_0, \alpha_1 \in$ [0.1,1] with N=10) has been used for the simulations. For the rightmost panel in D and E, and for F, $\alpha_0$=0.34 and $\alpha_1$=0.81 for the bistable and $\alpha_0$=0.18 and $\alpha_1$=0.5 for the monostable case.<br><br>G) Same as in Fig. 2L.<br>Initial conditions: $(A, F, S)\|_{t=0}$ = $(A^{ss(S=0)}$, 0, 0) where $A^{ss(S=0)}$ is the AP2 steady state value reached when $S$ = 0 (i.e., $\beta_s$=0). |
| Fig. S7 | See Supp. Tables 5 and 6 contain information about the fits. |
| Fig. S8 (M1) | A–F) Same as Fig. 2K, 2L, 3D, and 3J. For rAP2, $(\alpha_0^{rAP2},\alpha_1^{rAP2})=(\alpha_0^{wt},\alpha_1^{wt})$ and no inhibition of age on AP2. $\beta_s^{rAP2}$=2.5.<br><br>C-F) $\tau_0$ = t($S^{sim}$=$S^{SN}$), $\tau_g$ ends when $\partial A/\partial t$=0 (second maximum). Thrs.($\tau_e$)= 0.5.<br>Initial conditions: $(A, F, S)\|_{t=0}$ = $(A^{ss(S=0)}$, 0, 0) where $A^{ss(S=0)}$ is the AP2 steady state value reached when $S$ = 0 (i.e., $\beta_s$=0).<br><br>G) Same as Fig. 2K, 2L, 3D, and 3J.<br><br>H–L) Experimental data. Splines have been computed using the *gradient* function in Python, together with a *gaussian_filter1d* (n = 4). |
| Fig. S9 (M1) | Same as Fig. 2K, 2L, 3D, and 3J.<br>For rAP2, $(\alpha_0^{rAP2},\alpha_1^{rAP2})=(\alpha_0^{wt},\alpha_1^{wt})$ and no inhibition of age on AP2. $\beta_s^{rAP2}$=2.5. |
| Fig. S13 (M2) | Same as Fig. 4.<br>A-C) Monostable: $\alpha_0$=0.025 and $\alpha_1$=0.25. Bistable: $\alpha_0$=0.25 and $\alpha_1$=0.25.<br>D and E) Monostable: $\alpha_0$=0.025, $\alpha_1$=0.25 and $\beta_s$=3.<br>F and G) Bistable: $\alpha_0$=0.25 and $\alpha_1$=0.25 and $\beta_s$=3.<br>H and I) Monostable: $\alpha_0$=0.025 and $\alpha_1$=0.25. $\beta_s$=[0, 2, 4, 6, 8, 10, 20].<br>J and K) Bistable: $\alpha_0$=0.25 and $\alpha_1$=0.25. $\beta_s$=[0, 2, 4, 6, 8, 10, 20]. |

| | |
|---|---|
| | Initial conditions: $(Am, Ac, Ap, Fp, S)\|_{t=0} = (Am^{ss(S=0)}, Ac^{ss(S=0)}, Ap^{ss(S=0)}, 0, 0)$, where $Ax^{ss(S=0)}$ is the $A$x steady state value reached when $S = 0$ (i.e., $\beta_s$=0). |
| Fig. S14 (M2) | A and B) Same as Fig. 4.<br>$\alpha_0^{wt\text{-}M}$=0.01, $\alpha_0^{ful\text{-}M}$=0.0075, $\alpha_0^{soc1\text{-}M}$=0.001 and $\alpha_0^{soc1\ ful\ \text{-}M}$=0.0001.<br>$\alpha_1^{wt\text{-}M}$= $\alpha_1^{ful\text{-}M}$= $\alpha_1^{soc1\text{-}M}$= $\alpha_0^{soc1\ ful\ \text{-}M}$=0.25. In B), $\beta_s$=20.<br>C-K) Same as in Fig.5.<br>I-K) Same as in Fig.5. For *ap2*, $\alpha_0^{ap2}=\alpha_0^{wt}$, and $\alpha_1^{ap2}=0.25\bullet\alpha_1^{wt}$.<br><br>Initial conditions: $(Am, Ac, Ap, Fp, S)\|_{t=0} = (Am^{ss(S=0)}, Ac^{ss(S=0)}, Ap^{ss(S=0)}, 0, 0)$, where $Ax^{ss(S=0)}$ is the $A$x steady state value reached when $S = 0$ (i.e., $\beta_s$=0). For rAP2, $(Am, Ac, Ap, Fp, S)\|_{t=0}$ = (0, 2, 15, 0, 0). |
| Fig. S15 (M2) | Same as in Fig. 5 for wt (A), *ful* (B) and *soc1*.<br><br>Initial conditions: $(Am, Ac, Ap, Fp, S)\|_{t=0} = (Am^{ss(S=0)}, Ac^{ss(S=0)}, Ap^{ss(S=0)}, 0, 0)$, where $Ax^{ss(S=0)}$ is the $A$x steady state value reached when $S = 0$ (i.e., $\beta_s$=0). |
| Fig. S16 (M2) | Same as in Fig. 5 for *soc1 ful* (A) and *rAP2* (B). For *ap2* (C), as in Fig. S12I-S12K.<br>For rAP2, $(Am, Ac, Ap, Fp, S)\|_{t=0}$ = (0, 2, 15, 0, 0). In the comparison with the experimental data, normalisation is done with respect to $Ap^{t=0}$. |
| Fig. S17 (M2) | Same as in Fig. S15 with extrinsic noise. |
| Fig. S18 (M2) | Same as in Fig. S16 with extrinsic noise. |
| Fig. S19 (M2) | Same as in Fig. 4 with $\beta_s$=1.75. |
| Fig. S21 (M3) | Same as Fig. 6.<br>In G, $\beta_s$=0.23 (1), 0.50 (2), 0.78 (3), and 1.39 (4). |
| Fig. S22 (M2) | A)<br>+ wt-Model: Same as in Fig. 4 for wt.<br>+ wt-grid: ($\alpha_0, \alpha_1$)-point with lowest RMSE error in square grid.<br>+ wt-minimisation algorithm: ($\alpha_0$, $\alpha_1$)-point with lowest RMSE obtained with Nelder–Mead optimisation method initialised with wt-Model parameters.<br><br>B)<br>+ wt-Model: Same as in Fig. 4 for the different genotypes.<br>+ wt-grid: $\alpha_1^{mutants} = \alpha_1^{wt}$ (as in A). $\alpha_0$ for floral activator mutants is computed with the Nelder–Mead optimisation method of the RMSE under the condition $\alpha_0^{wt} > \alpha_0^{ful} > \alpha_0^{soc1} > \alpha_0^{soc1\ ful}$.<br>+ wt-minimisation algorithm (wt): $\alpha_1^{mutants} = \alpha_1^{wt}$ (as in A). $\alpha_0$ for floral activator mutants is computed with the Nelder–Mead optimisation method of the RMSE under the condition $\alpha_0^{wt} > \alpha_0^{ful} > \alpha_0^{soc1} > \alpha_0^{soc1\ ful}$.<br>+ wt-Model + min. mutants: $\alpha_1^{mutants} = \alpha_1^{wt}$ (as in Fig. 4). $\alpha_0$ for floral activator mutants is computed with the Nelder–Mead optimisation method of the RMSE under the condition $\alpha_0^{wt} > \alpha_0^{ful} > \alpha_0^{soc1} > \alpha_0^{soc1\ ful}$. |

| | |
|---|---|
| | *Ap* model curves were interpolated at experimental data points (7–24LDs) using the *interp1d* Python function.<br>C)<br>+ With two parameters: same as A.<br>+ With four parameters: ($\alpha_0$, $\alpha_1$, $\gamma$, offset)-RMSE minimisation performed with Nelder–Mead optimisation method, initialised with the values derived in the analysis with two parameters only. The specific values obtained are found in the figure itself.<br><br>D) Same as in C.<br><br>E) ($\alpha_0$, $\alpha_1$)-point with lowest RMSE in square grid. Scaling variables ($\gamma$, offset) and $\alpha_1$ as in A. Yellow dots: ($\alpha_0$, $\alpha_1$) pairs used for Model 2 for each genotype (Fig. 5).<br><br>F) Model: rAP2 parameters as Fig. 5G.<br>Minimum: ($k_{FS}$, $d_3$)-point with the lowest RMSE obtained in the square grid. |

**Table S3. Details about the normalisation used in the main figures.**

| **Figure** | *Details about the normalisation* |
| --- | --- |
| Fig. 2 (M1) | C) AP2-VENUS: normalised by its maximum median value (i.e., first time point).<br>SOC1-GFP: normalised by its maximum median value.<br><br>F, I, J, and L) *A* and *F* model numerical simulations are normalised individually for each panel by its maximum value, respectively.<br><br>M) AP2-VENUS experimental datapoints are normalised by the maximum median value, as in C. Numerically simulated AP2 is normalised by its maximum value. To compare experiment and model results simultaneously, a temporal rescaling of the simulation data ($t_{exp}$= $t_{num}\cdot\gamma$+offset) was implemented taking as reference the first time point at which AP2 reduction is completed, i.e., 17LDs ($\gamma$ = 10.75/125 and offset = 6.25). Note that the maximum AP2 value in experimental data and model results always occurs at the first time point. |
| Fig. 3 (M1) | C) AP2-VENUS concentrations are normalised by the maximum median value reached during the transition in wt (i.e., first time point in wt).<br><br>D) *AP2* model numerical simulations are normalised individually by its maximum value, respectively, but note in Fig. S7B with non-normalised results, that the maximum is identical for all genotypes (i.e., all simulations are normalised by the maximum reached in wt).<br><br>H) AP2-VENUS single-cell nuclear concentrations in *soc1 ful* and *rAP2* are normalised individually by the maximum median value reached for each genotype. Individual normalisation by the median value at the first time point (7LDs) was performed for AP2:rAP2::VENUS fluorescence because the transgene is inserted at a different genomic position than for AP2-VENUS, and the fluorescence intensity signal is therefore not directly comparable with AP2:AP2::VENUS of the other genotypes.<br>I) AP2-VENUS single-cell nuclear concentrations in *soc1 ful* and *rAP2* are normalised individually by the maximum median value and the median value at 7LDs.<br><br>J) Same as in D. |
| Fig. 4 (M2) | C) *Ap* and *Fp* model numerical simulations are normalised individually for each panel by their respective maximum values.<br><br>D) AP2-VENUS experimental data is normalised by its maximum median value. Numerically simulated AP2 is normalised by its maximum value. To compare experiment and model results simultaneously, a temporal rescaling of the simulation data ($t_{exp}$= $t_{num}\cdot\gamma$+offset) was implemented taking as reference the first time point at which AP2 reduction is completed, i.e., 17LDs ($\gamma$ = 17/125 and offset = 5.5). |

| | |
|---|---|
| Fig. 5 (M2) | B) *AP2p* model numerical simulations are normalised individually by their maximum values, which is identical in wt, *ful*, *soc1*, and *soc1 ful* genotypes (i.e., all simulations are normalised by the maximum reached in wt).<br><br>Third column in C–G: *Ap* and *Fp* model numerical simulations are normalised individually for each panel by their respective maximum values.<br><br>Fourth column in C–G: AP2-VENUS experimental data is normalised by the maximum median value reached in wt. Numerically simulated AP2 curves are normalised by their maximum values (i.e., the maximum is the same for wt, *ful*, *soc1* and *soc1 ful* simulations). For rAP2, which has a different transgene insertion from AP2-VENUS, rAP2-VENUS experimental data are normalised by its first median value (at 7LDs). The numerically simulated rAP2 curve is also normalised by its initial concentration, i.e., $Ap(t=0)$.<br>To compare experiment and model results simultaneously, a temporal rescaling of the simulation data ($t_{exp}= t_{num}\cdot\gamma$+offset) was implemented taking as reference the first time point at which AP2 reduction is completed, i.e., 17LDs ($\gamma$ = 17/125 and offset = 5.5). For consistency, the same rescaling factor was used for all genotypes. |
| Fig. 6 (M3) | A) AP2-VENUS and SOC1-GFP spatial concentration profiles along the SAM longitudinal axes are normalised separately by the maximum median value reached among all time points.<br><br>B, C and D) *Ap*, *Fp* and *S* model spatiotemporal numerical simulations are normalised individually by their global maximum values reached along the simulation.<br><br>E) *Ap* and *Fp* are normalised by their maximum values across the four scenarios displayed; *S* has been normalised within each scenario. |

**Table S4. Details about the normalisation used in supplementary figures.**

| **Figure** | *Details about the normalisation* |
|---|---|
| Fig. S2 | A) AP2-VENUS: normalised by the maximum median value reached in the two experiments.<br><br>D–F) AP2-VENUS: normalised by the maximum median value<br>SOC1-GFP: normalised by its maximum median value.<br>Each characteristic domain definition (i.e., CD=50, CD=75%, CS=0-20 μm) has its own normalisation. |
| Fig. S3 | C, F, and I) *A* and *F* model numerical simulations are normalised individually for each panel by their respective maximum values. |
| Fig. S5 | C) *Ap* and *Fp* model numerical simulations are normalised individually for each panel by their respective maximum values.<br><br>D and E) Each curve is individually normalised by its maximum - the maximum of the absolute value of the curve in E.<br><br>G) *A*, *F*, and *S* model variables are normalised by their maximum values reached in the simulation. |
| Fig. S6 | C, D, H, and I) Normalisation of fluorescence concentration measurements is performed with respect to the maximum median value in wt (which is also the initial (7LDs) wt-concentration). For rAP2-VENUS, since the transgene insertion is different to that of AP2-VENUS, fluorescence measurements were normalised with respect to its initial concentration (at 7LDs). |
| Fig. S7 | A, B, and E): Normalisation of AP2-VENUS fluorescence concentration measurements is performed with respect to the maximum median value in wt (which is also the initial (7LDs) wt-concentration). |
| Fig. S8 | C) The absolute value of the derivative of AP2 numerical simulation is normalised with respect to its maximum.<br><br>F) Normalisation is performed with respect to the total sum of times for each genotype.<br><br>H–L) Analyses are computed on normalised data shown in Figure 3. |
| Fig. S10 | C) Each intensity profile corresponds to a different SAM and is individually normalised by its maximum value. |
| Fig. S12. | A and B) Normalisations performed with respect to the maximum *AP2p* reached in each genotype (the maximum is the same for the four genotypes).<br><br>C–E) Normalisations are performed individually for each genotype with respect to the maximum *AP2p* reached in either the bifurcation diagram or the numerical simulation. The age signal is normalised by its own maximum. |

| | |
|---|---|
| | H) SOC1-GFP experimental data is normalised by its maximum median value. Numerically simulated *SOC1p* is also normalised by its maximum value. To compare experiment and model results simultaneously, a temporal rescaling of the simulation data ($t_{exp}= t_{num}\cdot\gamma$+offset) was implemented taking as reference the first time point at which AP2 reduction is completed, i.e., 17LDs ($\gamma$ = 17/125 and offset = 5.5). |
| Fig. S20 | C and D) AP2-VENUS spatial concentration profiles along the SAM longitudinal axes in wt, *ful*, *soc1*, and *soc1 ful* genotypes are normalised by the maximum median value reached among all time points in wt. rAP2-VENUS spatial concentration profile is normalised by its own maximum. |
| Fig. S21. | E and F) As in Fig. 6. *Ap*, *Fp*, and S model spatiotemporal numerical simulations are normalised individually by their global maximum values reached along the simulation.<br><br>G) *Ap* and *Fp* are normalised by their maximum values across the four scenarios displayed; S has been normalised within each scenario. |
| Fig. S22 | B) AP2-VENUS experimental data is normalised by the maximum median value reached in wt. Numerically simulated AP2 curves are normalised by their maximum values (i.e., the maximum is the same for wt, *ful*, *soc1*, and *soc1 ful* simulations).<br><br>To compare experiment and model results simultaneously, a temporal rescaling of the simulation data ($t_{exp}= t_{num}\cdot\gamma$+offset) was implemented taking as reference the first time point at which AP2 reduction is completed, i.e., 17LDs ($\gamma$ = 17/125 and offset = 5.5). For consistency, the same rescaling factor was used for all genotypes.<br><br>F) rAP2-VENUS experimental data is normalised by its first median value (at 7LDs). The numerically simulated rAP2 curve is also normalised by its initial concentration, i.e., *Ap*(*t*=0). |

**Table S5. Parameters and results from the fits in Figure S7 for wt.** Comparison of a single exponential, a sum of two exponentials, and a sum of two arctangent functions fitted to AP2-VENUS fluorescence data in wt. The table shows the exact formulas, initial parameter guesses, and final optimised parameters. The last four columns present different statistical criteria to evaluate the goodness of each fit. The sum of squared errors (SSE) is computed as $SSE = \sum_{i=1}^{N}(y_i - \hat{y}_i)^2$, where $y_i$ and ($\hat{y}_i$) correspond to the observed and predicted values, respectively. The root mean squared error (RMSE) is given by $RMSE = \sqrt{SSE/N}$. $R^2$ (R-square) is the coefficient of determination, representing the proportion of the variance in the dependent variable that is explained by the independent variable and is calculated as $R^2 = 1 - SSE/SST$, where SST is the total sum of squares, $SST = \sum_{i=1}^{N}(y_i - \bar{y}_i)^2$, with $\bar{y}_i$ the sample mean. $R^2_{adj}$ (adjusted R-square) is a modified version of the $R^2$ that accounts for the number of predictors in a regression model, providing a more accurate measure of the goodness of the fit; $R^2_{adj} = 1-((1-R^2)(n-1))/(n-p-1)$, where n is the number of samples and p is the number of parameters to predict. (*) indicates parameters that were fixed during the optimisation to represent the normalised initial AP2 concentration at 7LDs. Fits were performed using the function *fit* in MATLAB with no imposed parameter bounds. Optimisation settings were configured with *MaxFunEvals* set to 2000, and *MaxIter* set to 10000; all other fit parameters were maintained at MATLAB default values. Parameter initialisation and definitions: for the sum of two exponentials, initial guesses were selected to identify two distinct timescales ($b_1$>$b_2$) departing from the initial AP2 concentration ($a_1$+$a_2$=1). In the sum of two arctangent functions: *H* represents the normalised initial AP2 concentration; $a_i$ accounts for the magnitudes of the two drops; $x_i$ denotes the midpoints of the transition (where the distance $t_2$-$t_1$ defines the duration of the critical slowing down, $\Delta\tau$); and $k_i$ determines the steepness of each decline.

| **wt** | | | | | | |
|---|---|---|---|---|---|---|
| **Function** | **Parameter start point** | **Parameter values** | **$R^2$** | **$R^2_{adj}$** | **SSE** | **RMSE** |
| Single exponential: $y = ae^{-b(t-7)}$ | $a = 1.0$<br>$b = 0.1$ | $a = 1.008$<br>$b = 0.140$ | 0.9674 | 0.9609 | 0.0020 | 0.0630 |
| Sum of 2 exponentials: $y = \sum_{i=1}^{n=2} a_i e^{-b_i(t-7)}$ | $a_1 = 0.7$<br>$b_1 = 0.1$<br>$a_2 = 0.3$<br>$b_2 = 0.01$ | $a_1 = 2.608$<br>$b_1 = 0.083$<br>$a_2 = -1.617$<br>$b_2 = 0.059$ | 0.9725 | 0.9451 | 0.0168 | 0.0747 |
| Sum of 2 arctangents: $y = H - \sum_{i=1}^{n=2} a_i \left(\frac{1}{2} + \frac{1}{\pi} arctan(k_i(t-t_i)\right)$ | $H = 1.1$*<br>$a_1 = 1.0$<br>$k_1 = 0.5$<br>$t_1 = 12.0$<br>$a_2 = 0.1$<br>$k_2 = 0.35$<br>$t_2 = 15.0$ | $H = 1.1$*<br>$a_1 = 0.559$<br>$k_1 = 1.102$<br>$t_1 = 8.723$<br>$a_2 = 0.433$<br>$k_2 = 1.299$<br>$t_2 = 14.80$ | 0.9990 | 0.9942 | 0.0006 | 0.0244 |

**Table S6. Parameters and results from the fits in Figure S7 for *soc1*.** Comparison of a single exponential, a sum of two exponentials, and a sum of two arctangent functions fitted to AP2-VENUS fluorescence data in *soc1*. The table shows the exact formulas, initial parameter guesses, and final optimised parameters. The last four columns present different statistical criteria to evaluate the goodness of each fit, as in Table S5. Asterisks (*) indicate parameters that were fixed during the optimisation to represent the normalised initial AP2 concentration at 7LDs.

| ***soc1*** | | | | | | |
|---|---|---|---|---|---|---|
| **Function** | **Parameter start point** | **Parameter values** | **$R^2$** | **$R^2$ adj.** | **SSE** | **RMSE** |
| Single exponential: $y = ae^{-b(t-7)}$ | $a = 1.0$<br>$b = 0.1$ | $a = 0.979$<br>$b = 0.127$ | 0.9832 | 0.9808 | 0.0012 | 0.0414 |
| Sum of 2 exponentials: $y = \sum_{i=1}^{n=2} a_i e^{-b_i(t-7)}$ | $a_1 = 0.7$<br>$b_1 = 0.1$<br>$a_2 = 0.3$<br>$b_2 = 0.01$ | $a_1 = -0.245$<br>$b_1 = 0.135$<br>$a_2 = 1.224$<br>$b_2 = 0.129$ | 0.9832 | 0.9731 | 0.0120 | 0.0489 |
| Sum of 2 arctangents: $y = H - \sum_{i=1}^{n=2} a_i \left(\frac{1}{2} + \frac{1}{\pi} arctan(k_i(t-t_i)\right)$ | $H = 1.1$*<br>$a_1 = 1.0$<br>$k_1 = 0.1$<br>$t_1 = 10.0$<br>$a_2 = 0.15$<br>$k_2 = 1.0$<br>$t_2 = 22.0$ | $H = 1.1$*<br>$a_1 = 0.920$<br>$k_1 = 0.406$<br>$t_1 = 8.723$<br>$a_2 = 0.179$<br>$k_2 = 2.68$<br>$t_2 = 23.54$ | 0.9739 | 0.9304 | 0.0186 | 0.0787 |

**Table S7. Mann-Whitney-Wilcoxon-test values for wt-AP2 comparison in Figure S2A** Mann-Whitney-Wilcoxon-test tables for the median values of AP2-Venus wt-concentration from two independent experiments. Significance level, $\alpha$ = 0.05. Red values indicate non-significant differences between the medians, i.e., those cases in which the null hypothesis cannot be rejected**.**

| | 7LDs | 10LDs | 12LDs | 14LDs | 17LDs | 19LDs | 21LDs |
|---|---|---|---|---|---|---|---|
| wt-2 & wt-3 | --------- | 0.5941 | 0.6730 | **0.0170** | 0.1128 | 0.7913 | **0.0098** |

**Table S8. Mann-Whitney-Wilcoxon-test values for height and width in wt-AP2 and wt-SOC1 comparison in Figures S2B and S2C.** Mann-Whitney-Wilcoxon-test tables for the median values of AP2-Venus and SOC1-GFP wt-height and width from three independent experiments. Significance level, $\alpha$ = 0.05. Red values indicate non-significant differences between the medians, i.e., those cases in which the null hypothesis cannot be rejected**.**

| **Height** | 7LDs | 10LDs | 12LDs | 14LDs | 17LDs | 19LDs | 21LDs |
|---|---|---|---|---|---|---|---|
| wt-1 & wt-2 | 0.2031 | 0.0829 | 0.7577 | 0.3638 | 0.4363 | 0.1457 | 0.2359 |
| wt-1 & wt-3 | 0.8125 | 0.6038 | 0.5358 | 0.4757 | 0.0535 | 0.0545 | **0.0091** |
| wt-2 & wt-3 | 0.2810 | 0.1292 | 1.0000 | 0.1824 | **0.0021** | 0.2730 | **0.0062** |

| **Width** | 7LDs | 10LDs | 12LDs | 14LDs | 17LDs | 19LDs | 21LDs |
|---|---|---|---|---|---|---|---|
| wt-1 & wt-2 | 0.1166 | 0.6993 | 0.0848 | **0.0250** | 0.2224 | 0.3473 | 0.6551 |
| wt-1 & wt-3 | 0.7931 | 0.7802 | 0.6362 | 0.2500 | **0.0030** | **0.0013** | **0.0001** |
| wt-2 & wt-3 | 0.0541 | 0.9530 | 0.1590 | **0.0235** | **0.0006** | **0.0028** | **0.0003** |

**Table S9. Mann-Whitney-Wilcoxon-test values for wt-AP2 and wt-SOC1 comparison in Figures S2D–S2F.** Mann-Whitney-Wilcoxon-test tables for the median values of AP2-Venus and SOC1-GFP concentration for 50 and 75% thresholds characteristic domains and first 20 µm from the SAM. Significance level, $\alpha$ = 0.05. Red values indicate non-significant differences between the medians, i.e., those cases in which the null hypothesis cannot be rejected.

| **CD = 50%** | 7-10 | 10-12 | 12-14 | 14-17 | 17-19 | 19-21 | 21-24 |
|---|---|---|---|---|---|---|---|
| AP2-wt | **0.0089** | **0.0453** | 0.4436 | **$3.609 \cdot 10^{-8}$** | **0.0036** | **0.0223** | **$3.711 \cdot 10^{-5}$** |
| SOC1-wt | **0.0057** | **0.0115** | **0.0262** | 0.7577 | 0.8148 | 0.7209 | --------- |

| **CD = 75%** | 7-10 | 10-12 | 12-14 | 14-17 | 17-19 | 19-21 | 21-24 |
|---|---|---|---|---|---|---|---|
| AP2-wt | **0.0061** | **0.0102** | 0.6418 | **$9.708 \cdot 10^{-8}$** | **0.0001** | **0.0016** | **0.0005** |
| SOC1-wt | **0.0057** | **0.0021** | **0.0070** | 1.0000 | 0.1672 | 0.1304 | --------- |

| **20 µm** | 7-10 | 10-12 | 12-14 | 14-17 | 17-19 | 19-21 | 21-24 |
|---|---|---|---|---|---|---|---|
| AP2-wt | **0.0129** | **0.0102** | 0.8267 | **$9.708 \cdot 10^{-8}$** | **0.0003** | 0.4094 | **0.0102** |
| SOC1-wt | **0.0653** | **0.0418** | **0.0070** | 0.1738 | 0.6730 | **0.0379** | --------- |

**Table S10. Mann-Whitney-Wilcoxon-test values for height and width comparison between wt, *ful,* and *soc1* in Figures S6A and S6B.** Mann-Whitney-Wilcoxon-test tables for the median values of wt*, ful,* and *soc1* height and width (wt-data grouped from wt-2 and wt-3). Significance level, $\alpha$ = 0.05. Red values indicate non-significant differences between the medians, i.e., those cases in which the null hypothesis cannot be rejected.

| **Height** | 7LDs | 10LDs | 12LDs | 14LDs | 17LDs | 19LDs | 21LDs | 24LDs |
|---|---|---|---|---|---|---|---|---|
| wt & *ful* | 0.2184 | 0.5280 | **0.0135** | 0.4441 | 0.3839 | **0.0178** | 0.0950 | 0.0770 |
| wt & *soc1* | 0.1857 | **0.0056** | 0.0858 | **$3.3 \cdot 10^{-5}$** | **0.0001** | **0.0204** | **$1.5 \cdot 10^{-5}$** | **$2.2 \cdot 10^{-5}$** |
| *ful* & *soc1* | **0.0140** | 0.1672 | 0.1286 | **0.0021** | **0.0010** | 0.9682 | **0.0002** | **0.0003** |

| **Width** | 7LDs | 10LDs | 12LDs | 14LDs | 17LDs | 19LDs | 21LDs | 24LDs |
|---|---|---|---|---|---|---|---|---|
| wt & *ful* | 0.6816 | 0.3383 | 0.0021 | 0.0857 | 0.4170 | 0.4863 | 0.2816 | 0.3281 |
| wt & *soc1* | 0.0076 | 0.0180 | 0.1294 | 0.0084 | 0.0474 | **$2.7 \cdot 10^{-5}$** | **$1.0 \cdot 10^{-5}$** | **$2.2 \cdot 10^{-5}$** |
| *ful* & *soc1* | 0.0056 | 0.3213 | 0.0024 | 0.1502 | **0.0817** | 0.0015 | **0.0002** | **$2.2 \cdot 10^{-5}$** |

**Table S11. Mann-Whitney-Wilcoxon-test values for AP2-Venus comparison in wt, *ful,* and *soc1* from Figures S6C-S6E.** Mann-Whitney-Wilcoxon-test tables for the median values of AP2-Venus concentration for 50 and 75%-threshold characteristic domains and first 20 µm from the SAM between wt, *ful* and *soc1*. Significance level, $\alpha$ = 0.05. Red values indicate non-significant differences between the medians, i.e., those cases in which the null hypothesis cannot be rejected**.**

| **CD=50%** | 7LDs | 10LDs | 12LDs | 14LDs | 17LDs | 19LDs | 21LDs | 24LDs |
|---|---|---|---|---|---|---|---|---|
| wt & *ful* | **0.0003** | 0.5832 | **0.0055** | 0.3573 | 0.0126 | **0.6760** | 0.9506 | **$4.1 \cdot 10^{-5}$** |
| wt & *soc1* | 0.7984 | 0.9051 | 0.5899 | 0.1218 | **0.0027** | **0.0008** | **0.0008** | 0.2428 |
| *ful* & *soc1* | **0.0003** | 0.6058 | **0.0276** | 0.0640 | 0.3374 | **0.0028** | **0.0049** | **0.0133** |

| **CD=75%** | 7LDs | 10LDs | 12LDs | 14LDs | 17LDs | 19LDs | 21LDs | 24LDs |
|---|---|---|---|---|---|---|---|---|
| wt & *ful* | **0.0003** | 0.9229 | **$9.5 \cdot 10^{-5}$** | 0.2642 | **0.0019** | 0.3910 | 0.1543 | 0.0770 |
| wt & *soc1* | 0.1049 | 0.9525 | **0.0177** | 0.0628 | **0.0079** | **0.0001** | **0.0003** | 0.1564 |
| *ful* & *soc1* | 0.0059 | 0.6730 | **0.0402** | **0.0211** | 0.8590 | **0.0017** | **0.0002** | 0.7197 |

| **20 µm** | 7LDs | 10LDs | 12LDs | 14LDs | 17LDs | 19LDs | 21LDs | 24LDs |
|---|---|---|---|---|---|---|---|---|
| wt & *ful* | **0.0012** | 0.6283 | **0.0084** | 0.5437 | **0.0015** | 0.4679 | 0.6949 | **0.0078** |
| wt & *soc1* | 1.0000 | 0.8580 | **0.0177** | 0.0848 | **0.0304** | **0.0006** | **0.0003** | **0.0220** |
| *ful* & *soc1* | **0.0006** | 0.6730 | 0.6485 | **0.0452** | 0.4138 | **0.0036** | **0.0002** | 0.9048 |

**Table S12. Mann-Whitney-Wilcoxon-test values for height and width comparison between wt, *rAP2,* and *soc1 ful* in Figures S6F and S6G.** Mann-Whitney-Wilcoxon-test tables for the median values of wt, *rAP2,* and *soc1 ful* height and width (wt-data grouped from wt-2 and wt-3). Significance level, $\alpha$ = 0.05. Red values indicate non-significant differences between the medians, i.e., those cases in which the null hypothesis cannot be rejected**.**

| **Height** | 7LDs | 10LDs | 12LDs | 14LDs | 17LDs | 19LDs | 21LDs | 24LDs |
|---|---|---|---|---|---|---|---|---|
| wt & *rAP2* | 0.3446 | 0.1573 | **0.0019** | 0.7883 | **0.0014** | **0.0001** | **0.0002** | **0.0006** |
| wt & *soc1 ful* | 0.5996 | 0.6915 | **0.0001** | **$3.1 \cdot 10^{-6}$** | **0.0007** | **0.0063** | **0.0018** | **0.0010** |
| *rAP2* & *soc1 ful* | 0.2824 | 0.2031 | 0.3599 | **0.0010** | **0.0002** | **0.0170** | 0.0673 | 0.8691 |

| **Width** | 7LDs | 10LDs | 12LDs | 14LDs | 17LDs | 19LDs | 21LDs | 24LDs |
|---|---|---|---|---|---|---|---|---|
| wt & *rAP2* | 0.7210 | 0.0583 | 0.3382 | 0.2234 | **$2.6 \cdot 10^{-5}$** | **$3.7 \cdot 10^{-6}$** | **$5.2 \cdot 10^{-6}$** | **0.0001** |
| wt & *soc1 ful* | 1.0000 | 0.5580 | **0.0148** | **0.0291** | 0.3533 | **$9.0 \cdot 10^{-6}$** | **0.0009** | **0.0032** |
| *rAP2* & *soc1 ful* | 0.6620 | 0.0676 | 0.3599 | 0.0550 | **0.0002** | **0.0002** | **0.0003** | 0.0927 |

**Table S13. Mann-Whitney-Wilcoxon-test values for AP2-Venus comparison in wt and *soc1 ful* from Figures S6H and S6I.** Mann-Whitney-Wilcoxon-test tables for the median values of AP2-Venus concentration for 50 and 75%-threshold characteristic domains between wt and *soc1 ful*. Significance level, $\alpha$ = 0.05. Red values indicate non-significant differences between the medians, i.e., those cases in which the null hypothesis cannot be rejected**.**

| **CD=50%** | 7LDs | 10LDs | 12LDs | 14LDs | 17LDs | 19LDs | 21LDs | 24LDs |
|---|---|---|---|---|---|---|---|---|
| wt & *soc1 ful* | 0.3450 | 0.3601 | **$3.4 \cdot 10^{-5}$** | **0.0010** | **0.0004** | **$1.5 \cdot 10^{-5}$** | **0.0258** | **0.0001** |

| **CD=75%** | 7LDs | 10LDs | 12LDs | 14LDs | 17LDs | 19LDs | 21LDs | 24LDs |
|---|---|---|---|---|---|---|---|---|
| wt & *soc1 ful* | 0.4136 | 0.0907 | **$2.8 \cdot 10^{-5}$** | **0.0010** | **0.0036** | **$5.4 \cdot 10^{-6}$** | **0.0002** | **0.0001** |

**Table S14. Sample size for each experiment and genotype.** Number of sample size by experiment and genotype. Each sample is a different SAM grown under the same conditions.

| **Experiment 01: SOC1-GFP in wt and *ap2*** | | | | | | | | | | |
|---|---|---|---|---|---|---|---|---|---|---|
| List of figures: Figures 1B, 1C, 6A, S2B-S2F, S14H, S20B and S20E | | | | | | | | | | |
| | Number of samples analysed (n) by data points (LDs) | | | | | | | | | |
| Genotype | 7 | 10 | 12 | 14 | 17 | 19 | 21 | 24 | 26 | 29 |
| wt | 10 | 9 | 7 | 7 | 9 | 8 | 8 | - | - | - |
| *ap2* | 10 | 8 | 7 | 7 | 8 | 9 | 6 | - | - | - |
| **Experiment 02: AP2-VENUS in wt, *soc1* and *rAP2*** | | | | | | | | | | |
| List of figures: Figures 2C, 2M, 3B, 3C, 3G–I, 4D, 5C, 5E, 5G, 6A, S2, S6, S7A, S7E, S8H, S8I, S10A–C, S11A, S11C, S11E, S12A, S12C, S12E, S20A, S20C, S20D and S22B. | | | | | | | | | | |
| | Number of samples analysed (n) by data points (LDs) | | | | | | | | | |
| Genotype | 7 | 10 | 12 | 14 | 17 | 19 | 21 | 24 | 26 | 29 |
| wt | 8 | 5 | 9 | 10 | 9 | 10 | 9 | 9 | - | - |
| *soc1* | 8 | 9 | 9 | 9 | 9 | 10 | 9 | 10 | 10 | - |
| *rAP2* | 8 | 8 | 8 | 8 | 10 | 10 | 10 | 10 | - | - |
| **Experiment 03: AP2-VENUS in wt, *ful* and *soc1 ful*** | | | | | | | | | | |
| List of figures: Figure 2A, 2C, 2M, 3A, 3C, 3F, 3H, 3I, 4D, 5C, 5D, 5F, S2, S6, S7B, S7E, S8H, S8I, S11B, S11D, S11D, S20C, S20D and S22B. | | | | | | | | | | |
| | Number of samples analysed (n) by data points (LDs) | | | | | | | | | |
| Genotype | 7 | 10 | 12 | 14 | 17 | 19 | 21 | 24 | 26 | 29 |
| wt | 7 | 10 | 8 | 13 | 10 | 10 | 11 | - | - | - |
| *ful* | 7 | 8 | 11 | 10 | 12 | 10 | 11 | 9 | - | - |
| *soc1 ful* | 6 | 10 | 10 | 13 | 10 | 13 | 13 | 12 | 12 | 10 |
